# Engineering HIV antibodies with enhanced breadth and potency of neutralization through multistate affinity maturation

**DOI:** 10.1101/2025.10.22.684031

**Authors:** Mateusz Kędzior, Swastik Phulera, Monica L. Fernández-Quintero, Johannes R. Loeffler, Collin Joyce, Jordan Woehl, Abdolrahim Abbasi, Maryam Karimi, Arash Aslanabadi, Mahsa Hojabri, Roza Zareidoodeji, Ben Atkinson, Eduar Fernando Pinzon Burgos, Alonso Heredia, Karen Saye-Francisco, Quoc Tran, Yoojin Kim, Fernando Acosta-Puente, Lara Shahin, Amelia Zhou, Pilar X. Altman, Nathaniel R. Felbinger, Brian G. Pierce, Devin Sok, Michael S. Seaman, Dennis R. Burton, Anthony L. DeVico, Andrew B. Ward, Gabriel Ozorowski, Mohammad M. Sajadi, Joseph G. Jardine

## Abstract

Broadly neutralizing antibodies (bnAbs) against HIV hold promise as therapeutic and prophylactic agents, but realizing this potential requires antibodies that function across the antigenic heterogeneity of the HIV envelope glycoprotein (Env). Although numerous bnAbs have been isolated from infected individuals, their breadth and potency may not be sufficient to tackle global viral diversity, motivating efforts to further improve their neutralization capacity. Here, we address this challenge using a multistate antibody engineering approach integrating deep mutational scanning with combinatorial library screening across diverse Env variants. This strategy enables identification of mutation patterns that confer improved binding across antigenically distinct targets. Starting from one of the best-in-class CD4-binding site bnAbs, we performed iterative optimization to increase binding affinity across diverse Env variants. The resulting lead candidate exhibited improved breadth and up to 100-fold higher potency against pseudoviruses from large cross-clade historical and contemporary panels while maintaining biophysical and pharmacokinetic profiles conducive to clinical development. Structural and molecular dynamics analyses revealed a unique tri-tyrosine aromatic triad and reinforced electrostatic contacts that stabilized the bnAb/Env interface. These findings demonstrate that systematic *in vitro* engineering can generate bnAbs with enhanced breadth and potency, providing a generalizable strategy for developing therapeutic antibodies against highly diverse pathogens.

## Introduction

Antibody-based therapeutics have emerged as powerful tools against infectious diseases, as demonstrated by recent clinical successes targeting Ebola virus^1^, respiratory syncytial virus^2^, and SARS-CoV-2^3^. For HIV, combination antiretroviral therapy (ART) remains the standard of care, effectively suppressing viral replication and reducing transmission^4,5,6^. However, ART does not eliminate latent viral reservoirs and thus requires lifelong adherence^7,8^. Broadly neutralizing antibodies (bnAbs) offer an additional means of viral control, capable of maintaining suppression during analytic treatment interruptions or in situations where ART delivery is challenging (e.g., prevention of mother-to-child transmission)^9,10,11^. Beyond neutralizing free virus, bnAbs can engage Fc-dependent effector functions to drive clearance of infected cells, an approach actively explored in HIV cure-focused strategies^12^.

A central challenge for using bnAbs as HIV countermeasures is the extraordinary antigenic diversity of the HIV envelope glycoprotein (Env), which can vary by up to 35% between isolates^13^ and undergoes rapid diversification within an individual during the course of infection^14^, creating many opportunities for viral escape^9^. Clinical trials of bnAb monotherapy during ART interruption have shown only transient viral control in most participants, with the eventual emergence of resistant variants and viral rebound^15,16,17,18^. In contrast, combinations of bnAbs targeting distinct epitopes have achieved more durable viral suppression and delayed escape^19,20,21^. These studies underscore the importance of maximizing bnAb neutralization breadth to reduce the risk of viral resistance. Moreover, the Antibody Mediated Prevention (AMP) trials demonstrated that prophylactic bnAb administration can establish sterilizing immunity, but only against strains highly sensitive to the bnAb^22^. Taken together, these studies demonstrate the potential of bnAbs while highlighting the necessity of maximizing their neutralization breadth and potency.

The discovery of potent HIV bnAbs has relied on screening for rare individuals with outstanding broadly neutralizing serum responses and then isolating the monoclonal antibodies responsible for that activity. Over the past two decades, hundreds of such bnAbs have been defined, converging on a relatively limited set of conserved Env epitopes^23,24,25,26^. Early discovery efforts frequently identified bnAbs within each epitope class that exhibited marked improvements in neutralization breadth or potency; however, as efforts have continued, such gains have plateaued, and antibodies surpassing current best-in-class examples are now rarely reported^24,27^. This stagnation likely reflects intrinsic limits of natural B cell affinity maturation.

During germinal center maturation, antibodies often achieve low-nanomolar to high-picomolar binding affinities. Beyond this range, selective pressure diminishes: on-rates are limited by both diffusion and the restricted density of antigen on follicular dendritic cells, off-rates slower than the timescale of antigen internalization provide little added benefit, and high-affinity B cells preferentially differentiate into plasma cells, curtailing further evolution^28,29,30^. In addition, somatic hypermutation samples sequence space unevenly, favoring hotspot motifs and preferentially generating mutations accessible by a single nucleotide change from the starting codon. A further limitation is the restricted viral diversity encountered during natural infection. Infection in most individuals is established by a limited number of closely related founder viruses, and only a minority ever become superinfected with a second, genetically distinct strain^31,32,33^. Consequently, the breadth of viral variants that a maturing bnAb is exposed to within a single host is far narrower than global HIV diversity, potentially constraining the breadth achievable through selection in that individual.

To overcome these natural maturation constraints, we and others have pursued rational engineering approaches to address HIV antigenic diversity and achieve greater breadth and potency in existing bnAbs. Previously, we showed how grafting the elongated framework 3 (FRH3) loop from one anti-CD4-binding site (CD4bs) bnAb, VRC03, onto other bnAbs of this class improves their neutralization and biochemical profiles^34,35^. We also reported that other, more generic engineering strategies can optimize clinical developability while preserving the enhanced bnAb function^36^.

In this study, we implemented a novel, multistate affinity maturation strategy designed to push HIV bnAb binding affinities beyond that which germinal centers typically produce. *In vitro* directed evolution techniques have previously produced protein binders with extraordinarily high affinities, including into the femtomolar range^37,38^. Since enhanced antibody/antigen affinity often translates into greater neutralization potency and resistance to viral escape, we hypothesized that artificially boosting bnAb affinity through directed evolution could significantly improve therapeutic breadth and potency^39,40,41,42^. Here, antibody libraries were screened in parallel against a panel of diverse Env immunogens representing global variability. By applying selection pressure simultaneously across multiple Env variants, we aimed to enrich mutations that promote improved generalizable Env binding, rather than affinity restricted to a single strain.

As a test case, we began with N49P7-FR, a CD4bs bnAb variant that was previously improved by grafting the elongated VRC03 FRH3 loop into the parental N49P7 antibody as noted above^35^. We carried out multistate affinity maturation of this bnAb in two phases: an initial deep mutational scanning stage to delineate the impact of all single mutations across the variable regions, followed by the screening of combinatorial libraries to identify synergistic combinations of globally beneficial mutations. Both phases employed parallel yeast display selections against a diverse panel of HIV gp120 variants. Deep sequencing of enriched populations enabled rapid identification of mutations that were consistently favored across all Env targets. Using this strategy, we developed engineered variants of N49P7-FR (denoted as eN49P7-FR) that achieved markedly enhanced binding affinity across our panel of gp120 variants. Correspondingly, the engineered bnAb exhibited substantially improved neutralization breadth and potency. Structural analyses revealed that the enriched mutations introduced a distinct mode of Env engagement, including the insertion of a triad of tyrosine residues into a conserved pocket of the CD4bs, thereby stabilizing antibody/antigen interactions. Molecular dynamics simulations further showed that these mutations enhanced electrostatic and van der Waals interactions, rigidified the binding interface, particularly the CDRH3 loop, and stabilized key contacts with Env residues and glycans. Together, these results establish a broadly applicable platform for engineering antibodies with optimized activity against structurally conserved yet antigenically heterogeneous targets. In the context of HIV, our findings highlight the potential of such engineered antibodies to advance therapeutic and functional cure strategies.

## Results

### 1. Optimization of N49P7-FR

To map the sequence/function landscape of N49P7-FR, we generated double barcode-enhanced saturated mutagenesis scanning (dbSMS) libraries of the variable heavy (VH) and variable light (VL) chains for yeast surface display (Fig. 1). These libraries incorporated all possible single amino acid substitutions, excluding cysteines, and included synonymous mutations flanking each targeted codon to serve as molecular barcodes for high-throughput deconvolution. These molecular barcodes enabled robust differentiation of true mutations from sequencing errors and allowed accurate normalization of mutation frequencies relative to the parental N49P7-FR residue. Yeast displaying these libraries as molecular Fab fragments were incubated with sub-saturating concentrations of His-tagged gp120 proteins, followed by fluorescently conjugated antibodies targeting the epitope tags used for Fab display and gp120 binding quantification. Each library was simultaneously screened by fluorescence-activated cell sorting (FACS) against a panel of six antigenically diverse gp120 monomers, alongside an unselected control population (collected in triplicate) used to establish baseline mutation frequencies (Supplementary Figs. 1–3). Following two rounds of FACS enrichment per antigen, antibody-encoding DNA from the sorted populations was deep sequence analyzed, generating high-resolution mutational enrichment profiles that quantitatively captured beneficial amino acid substitutions across VH and VL domains (Fig. 2 and Supplementary Figs. 8–10). Enrichment was calculated in two ways: (1) as normalized amino acid frequencies at each position and (2) as fold-change comparisons of sorted populations relative to the control. These analyses produced a comprehensive mutational fingerprint, allowing classification of each amino acid substitution as beneficial, neutral, or detrimental for Env binding.

**Figure 1.**
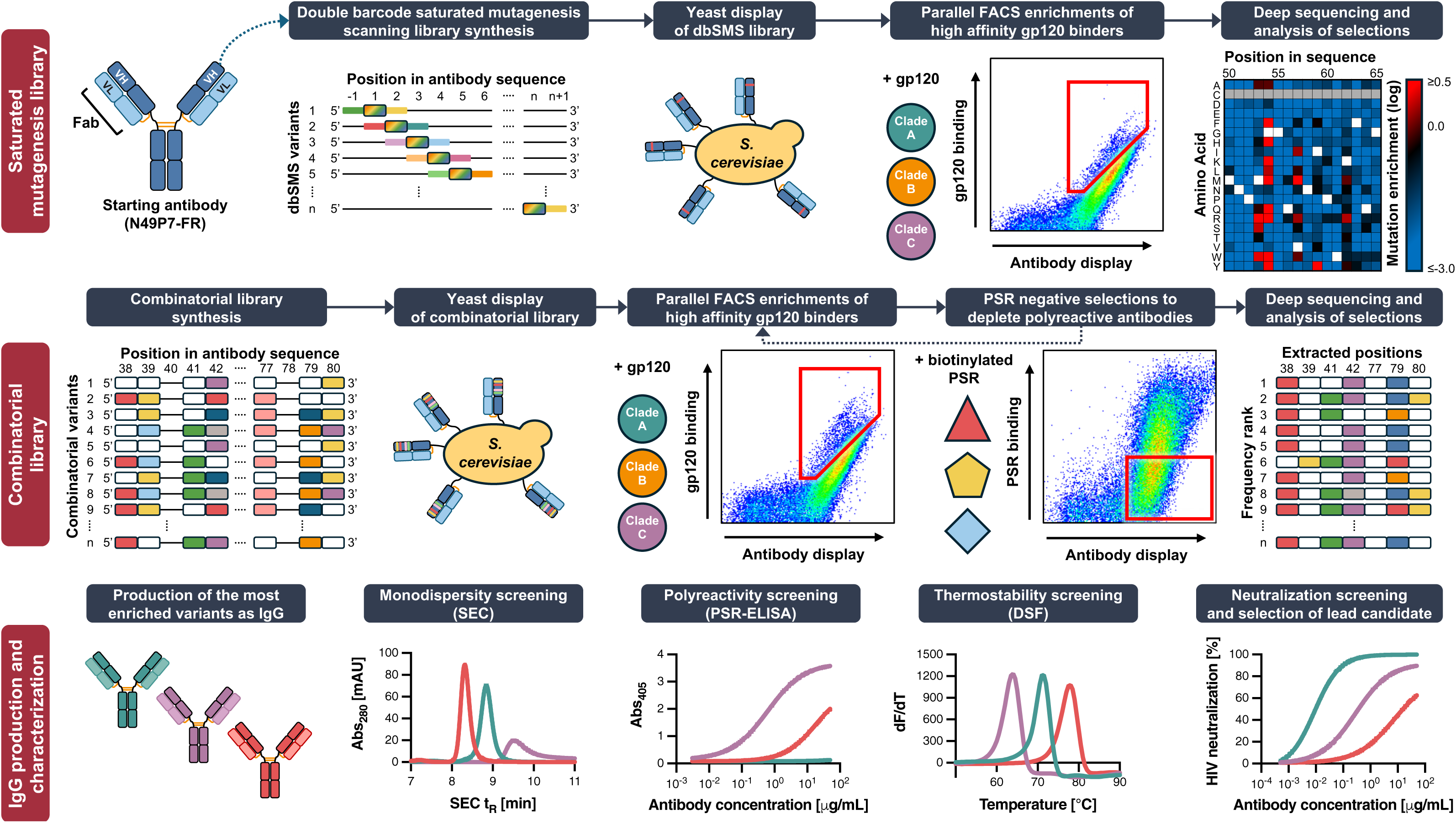
Workflow for *in vitro* bnAb affinity maturation via yeast display. *Top panel:* A dbSMS library was constructed separately for VH and VL regions of N49P7-FR. This library individually incorporated degenerate codons at all positions, sampling all amino acids except cysteine, with silent “barcode” mutations introduced into adjacent codons to enable precise identification of single amino acid substitutions. The libraries were displayed on the surface of yeast as molecular Fab fragments fused to a synthetic mucin-like domain and a GPI anchor. The yeast-displayed libraries were screened via FACS to select top 5-10% populations with improved binding to six diverse HIV gp120 proteins in parallel. The antibody-encoding vectors were recovered from enriched and unselected control populations and analyzed via deep sequencing. Amino acid frequencies in enriched datasets were normalized against the control to develop a sequence/binding profile for each single mutation in the antibody variable domains. *Middle panel:* Combinatorial libraries were then constructed to sample all possible combinations of the original amino acid and select favorable substitutions identified in the dbSMS screening. These combinatorial libraries were displayed on yeast and screened via FACS against twelve gp120 proteins in parallel. To avoid selecting for sticky antibody variants, positive selections were interspersed with negative selections using a polyspecificity reagent (PSR) to deplete polyreactive variants. Antibody-encoding DNA from enriched cells was deep sequenced to identify variants with improved binding across all twelve gp120 proteins. *Bottom panel:* The most enriched combinatorial variants were produced as IgGs and tested for neutralization potency and biochemical properties to identify a lead candidate.

**Figure 2.**
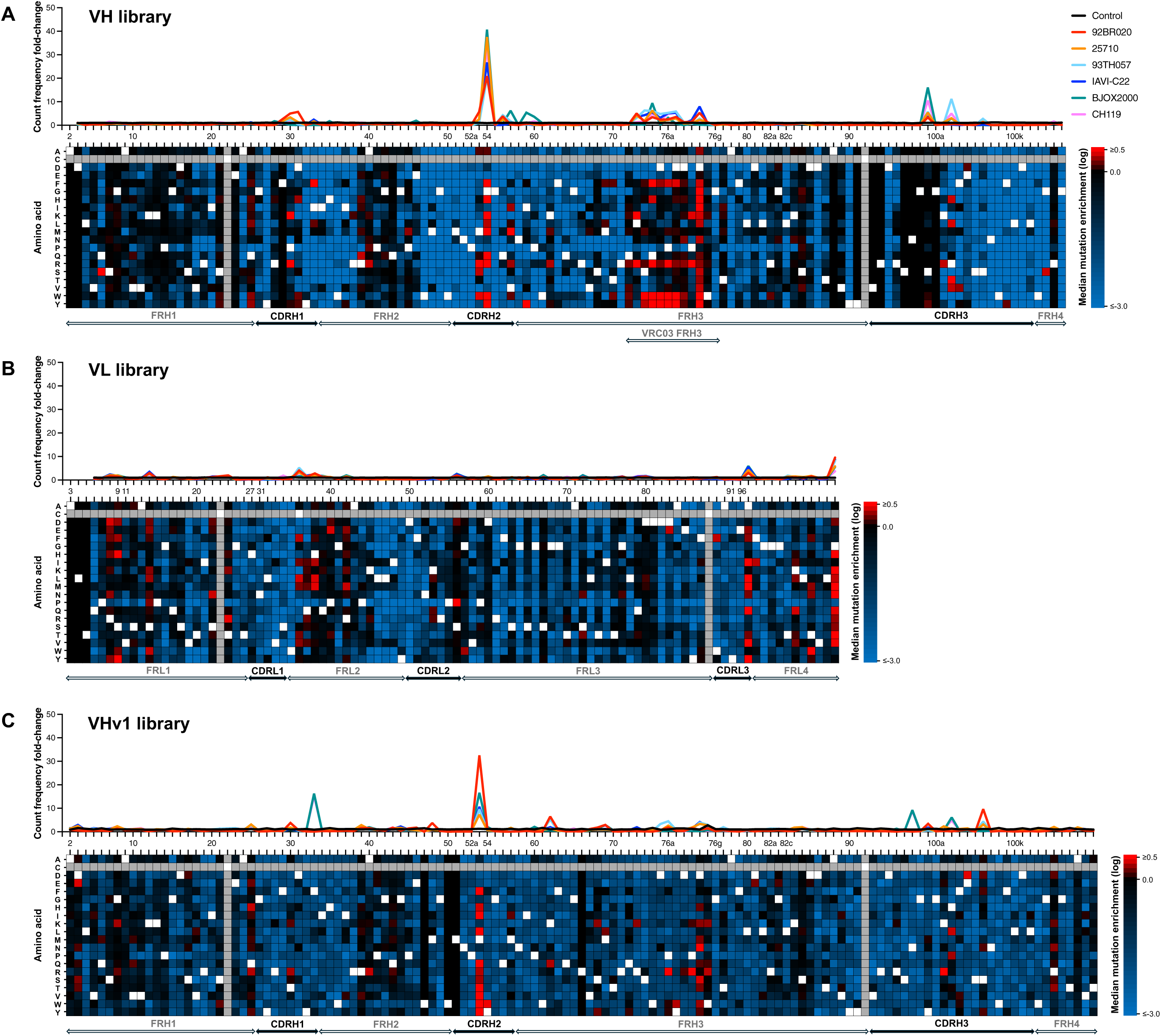
Mutation enrichment in the gp120-selected N49P7-FR dbSMS library variants. **A)** N49P7-FR initial VH library with fixed wild-type VL, **B)** VL library with fixed initially optimized VH, **C)** and subsequent VH library (VHv1) with fixed optimized VL were screened to identify the mutations enhancing the Fab affinity to six gp120s. *Upper part of each panel:* total variant count frequency per amino acid position in the gp120-selected dbSMS library normalized to total variant count frequency at the same amino acid position in the unselected (control) population expressed as count frequency fold-change. *Lower part of each panel:* heat map showing median amino acid enrichment (in log scale) across all variable region positions in the dbSMS library selected with six gp120s. Amino acids enhancing the gp120 interaction (red), diminishing it (blue), having a neutral effect (black), wild-type (white), and not sampled in the library (gray) are indicated. Amino acid positions are numbered according to the Kabat scheme. A threshold for the most significantly enriched mutations (≥0.5) was established based on the distribution of mutation enrichment values in the gp120-selected populations compared to the unselected control population (Supplementary Fig. 7).

Analysis of the deep sequencing data identified a small subset of mutations in N49P7-FR that were reproducibly enriched across all gp120 selections. Mutation enrichment profiles were more pronounced in the HC dataset (Fig. 2A), consistent with the binding mode of N49P7, in which the majority of antigen contacts are mediated through the HC^43^. The strongest enrichments were observed in CDRH2 and the FRH3 region, both located at the bnAb/Env interface. The most enriched mutations occurred at position 54 in the HC, where glycine was replaced by aromatic residues—phenylalanine, tyrosine, and tryptophan (Fig. 2A, Supplementary Fig. 8). The aromatic mutations at position 54 replicate a critical aromatic interaction between CD4 and HIV gp120, and have previously been associated with enhanced neutralization potency in various VRC01-class bnAbs^35,44,45^. However, engineered introduction of aromatic residues at position 54 in VRC01-class bnAbs frequently induces polyreactivity and reduces biochemical stability^46,47^. In contrast, several naturally occurring VRC01-class bnAbs isolated from HIV-infected donors evolved aromatic residues at this position and do not exhibit these detrimental effects, likely due to compensatory mutations elsewhere in the antibody^36,48,49^.

Unlike the CDRH2 region that directly contacts the CD4bs, pronounced enrichments in the FRH3 insertion were less expected. Structural analyses of other VRC01-class bnAbs with the FRH3 insertion show it interacts with the base of V1V2 and V3 loops, making both intra- and inter-protomer contacts^35^. However, our screening utilized monomeric gp120 rather than trimeric Env to avoid avidity-driven enrichments. Monomeric gp120 lacks ordered V1V2 loops and would recapitulate intra-protomer contacts. It is possible that transient interactions with flexible V1V2 loops could have contributed to the observed enrichments, though an alternative explanation is that these mutations represent subtle structural adjustments optimizing the FRH3 insertion in the N49P7 context. Given that the FRH3 insertion originated from VRC03 where it co-evolved within the context of the other VRC03 mutations, it is possible that the initial loop transfer into the N49P7 heavy chain is suboptimal and required compensatory mutations in this new context.

In contrast to the HC enrichments, the LC enrichments were far less pronounced. This is in agreement with the observation that the LC makes far fewer contacts with gp120 than the HC, with most of the interactions coming from CDRL3 and CDRL1^43^. The highest increase of mutation count frequency across all gp120-sorted populations compared to the control was observed in FRL2, CDRL3, and FRL4 (Fig. 2B). Within these regions, the most significantly enriched mutations were found at positions 36, 97, and 108. The original unpaired cysteine at position 36 (FRL2) mediates packing of the hydrophobic core at the VL-VH interface. Alanine at position 97 (CDRL3) and glycine at position 108 (FRL4) interact with the N-terminus of VL and CL, respectively. These three positions are involved in the intra- or interchain stabilization of the bnAb.

After identifying beneficial individual mutations through dbSMS, we next designed combinatorial libraries to assemble sets of mutations that together corrected distinct deficiencies in the antibody. In many cases, several substitutions could resolve the same limitation, meaning that only one was necessary while others became redundant or incompatible. Library design was guided primarily by the dbSMS datasets (Fig. 2). When multiple enriched mutations were identified at a single site, we also examined other naturally occurring VRC01-class bnAbs to assess whether such substitutions had arisen in different donors (Supplementary Fig. 11). To proactively enhance developability, we aimed to reduce hydrophobic surface patches, eliminate liabilities such as aspartic acid isomerization sites, and minimize potential immunogenicity by favoring mutations commonly found in germline VH genes.

In our optimization strategy, we first sought to identify the optimal HC variant while retaining the original LC, and then to screen LC variants for compatibility with the improved HC. For the HC, we targeted 16 positions, sampling between 1 and 4 substitutions at each site while always including the original amino acid (Supplementary Fig. 12A), resulting in a total library diversity of 2.2 × 10^6^ variants. The combinatorial HC library was displayed on yeast and divided into 12 fractions, each screened to enrich clones with improved binding to a different gp120 variant. The library underwent five rounds of selection (Supplementary Fig. 4): three positive selections with gp120 to isolate the highest affinity binders (AFF sorts), interspersed with two negative selections using the polyspecificity reagent (PSR) to eliminate polyreactive variants (PSR sorts). After the selections, the antibody-encoding DNA from the enriched cells was extracted and deep sequenced. We then analyzed the data to identify HC variants that showed the strongest enrichment across all datasets (Supplementary Fig. 13A). In total, we selected 24 sequences (Supplementary Table 1), each differing by at least two amino acids, for reformatting as IgG1 to assess their neutralization potency and biochemical properties.

Engineered VRC01-class bnAbs are prone to polyreactivity and developability issues^46,47^, so we began by characterizing the biophysical properties of the 24 HC variants. Of those 24, only five showed minimal polyreactivity and no aggregation (Supplementary Table 4), and four of these displayed increased neutralization potency against a small pseudovirus panel (Supplementary Fig. 14). The strongest variant, VHv1, owed much of its improved activity to a G54Y mutation, as reverting this substitution diminished neutralization potency. Despite these gains, all variants exhibited poor developability, specifically delayed SEC retention times, precipitation, and low purification yields (Supplementary Table 4), necessitating further optimization.

We selected VHv10, one of the most potent HC variants and the one with the fewest biochemical liabilities, as the starting point for LC optimization. A combinatorial LC library targeting 15 positions (Supplementary Fig. 12B) was generated, yielding a theoretical diversity of 3.3 x 10^6^ variants. Screening was performed in the same manner as for the HC library (Supplementary Fig. 5), and sequencing identified enriched LC variants across all datasets (Supplementary Fig. 13B). Twenty-one candidates were reformatted as IgG1 (Supplementary Table 2), none of which exhibited polyreactivity (Supplementary Table 5). Seventeen displayed monodisperse SEC profiles, and three showed substantially improved thermal stability and production yield while maintaining neutralization breadth and potency across the global pseudovirus panel (Supplementary Fig. 15).

After identifying LC variants that improved the biochemical behavior of VHv10, we revisited two other promising HC variants, VHv1 and VHv13, which had higher neutralization potency than VHv10 but carried more pronounced liabilities. When paired with VLv3, VHv1 largely retained the neutralization potency gains across the pseudovirus panel (Supplementary Fig. 16). It also showed substantial improvements in the overall biochemical behavior, though notable liabilities remained compared to the parental N49P7-FR and our clinical antibody controls (Supplementary Table 6).

To try to address these remaining liabilities, we used VHv1/VLv3 as the starting point for a final round of HC optimization. We generated a new dbSMS library based on the VHv1 HC and screened it as before. Compared to the first HC screen, fewer mutations showed strong enrichment, suggesting that most major improvements had already been captured in the initial round (Fig. 2C, Supplementary Fig. 10). When constructing the subsequent combinatorial library, we prioritized the few strongly enriching mutations but also incorporated a broad set of neutral mutations. This approach allowed us to sample different antibody surfaces in search of variants with improved biochemical properties. In total, the HC library targeted 28 positions, with 1–3 substitutions tested at each site while always retaining the parental residue (Supplementary Fig. 12C). Following selection (Supplementary Fig. 6) and sequencing, 96 enriched HC variants were reformatted as IgG1 (Supplementary Fig. 13C; Supplementary Table 3). None exhibited polyreactivity (Supplementary Table 7), and 10 showed SEC retention times within the range of clinical antibodies. Among these, three variants—VHv1-23/VLv3, VHv1-65/VLv3, and VHv1-85/VLv3—displayed the greatest neutralization breadth and potency, with substantial improvement over the starting bnAb and developability comparable to the parental antibody (Supplementary Fig. 17; Supplementary Table 8). Neutralization against AMP study-derived pseudoviruses further confirmed broad and potent activity (Supplementary Figs. 18–19). Based on its slightly superior overall potency and developability, VHv1-23/VLv3, hereafter designated eN49P7-FRv1-23, was selected as the lead candidate.

### 2. Functionality and developability of the optimized N49P7-FR

Through stepwise optimization of N49P7, including the initial incorporation of the FRH3 insertion^35^ and subsequent affinity maturation, we observed progressive gains in neutralization breadth and potency (Fig. 3A). The final lead candidate, eN49P7-FRv1-23, was on average 2.5-fold more potent than N49P7-FR, 5-fold more potent than the original N49P7, and an order of magnitude more potent than VRC01 against the 119-pseudovirus cross-clade panel (Fig. 3A). Using the IC_80_ ≤0.3 μg/mL cutoff (associated with >80% prevention efficacy in the VRC01 AMP trials^22^), eN49P7-FRv1-23 achieved 82% breadth, compared to 57% for N49P7-FR and 26% for N49P7. Only four viral strains in the panel were fully resistant (Supplementary Fig. 20). eN49P7-FRv1-23 also showed the highest gains in neutralization potency against AMP study-derived pseudoviruses, achieving near pan-coverage of the tested strains. Specifically, 93% and 100% clade B viruses from the placebo group and breakthrough clade C viruses, respectively, were neutralized at IC_80_ ≤0.3 μg/mL, compared to 38% and 9% neutralized by VRC01 (Fig. 3A). Furthermore, eN49P7-FRv1-23 demonstrated exceptional breadth and one of the highest mean potencies among best-in-class HIV bnAbs targeting diverse epitopes across the 119-pseudovirus global panel (Fig. 3B,C).

**Figure 3.**
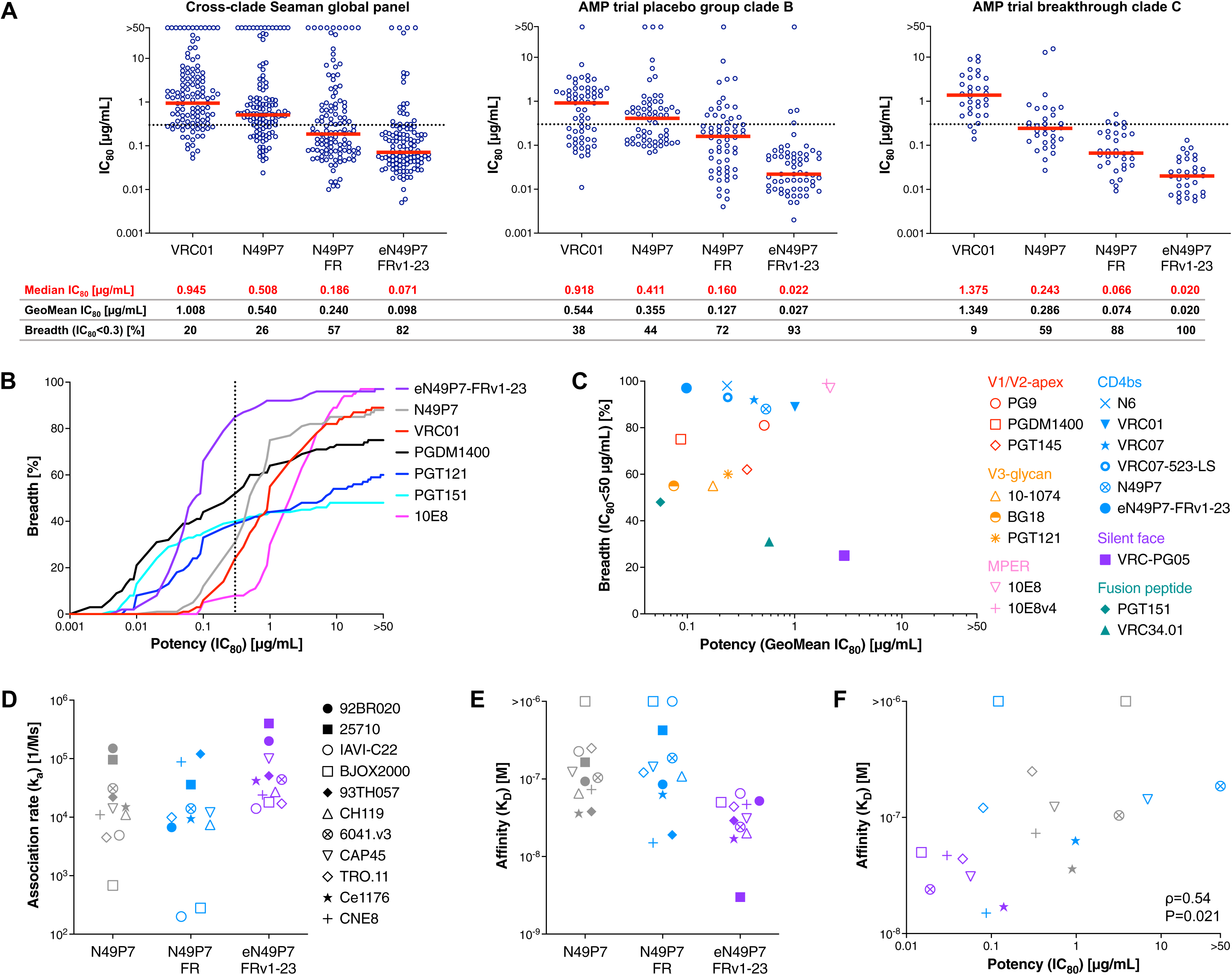
Evaluation of the optimized bnAb variant eN49P7-FRv1-23. **A**) *In vitro* neutralization potency and breadth of VRC01 and N49P7 variants against the indicated panels of HIV pseudoviruses. Each circle represents an IC_80_ value for one pseudovirus strain. Horizontal red bars indicate the median IC_80_ values, and the horizontal dotted line marks IC_80_=0.3 μg/mL, a cutoff estimated to correlate with >80% prevention efficacy in the VRC01 AMP trials^22^. **B**) Potency and corresponding breadth of N49P7 variants and other best-in-class bnAbs targeting different HIV epitopes against the 119-virus cross-clade Seaman global panel. The vertical dotted line marks IC_80_=0.3 μg/mL. **C**) Mean potency and breadth of N49P7 variants and other best-in-class bnAbs targeting different HIV epitopes against the 119-virus cross-clade Seaman global panel. The neutralization data for bnAbs other than VRC01, N49P7, and eN49P7-FRv1-23 were derived from the CATNAP database^23^. **D–F**) Binding kinetics of N49P7 variants against a panel of HIV cross-clade gp120s as determined by SPR. Association rate constant k_a_ (**D**), binding affinity expressed as equilibrium dissociation constant K_D_ (**E**), and significant positive correlation (Spearman’s ρ=0.54, P=0.021) between neutralization potency and binding affinity (**F**) are shown. Colors indicate specific bnAbs and symbols indicate interacting gp120s (**D**).

The hypothesis motivating our directed evolution campaign was that higher binding affinity would translate into increased neutralization potency. To better understand this correlation, we measured the binding affinities of N49P7-FR and its engineered variant against a panel of monomeric gp120s matched to pseudoviruses with varying neutralization sensitivities (Fig. 3D–F, Supplementary Fig. 21, Supplementary Table 9). Although monomeric gp120 does not fully capture the quaternary structure of native Env on the pseudovirus, it was the bait used in the directed evolution campaign that ultimately produced the eN49P7-FR variants. The neutralization potency gains observed for eN49P7-FRv1-23 corresponded closely with increased gp120 binding affinity, including against TRO.11 and Ce1176, two strains not used in the maturation campaign (Fig. 3F). Binding kinetics revealed that the enhanced affinity of eN49P7-FRv1-23 was primarily conferred by a faster association rate (Fig. 3D, Supplementary Fig. 21, Supplementary Table 9). By contrast, other engineered HIV bnAbs, such as VRC01-class antibodies carrying a potency-enhancing aromatic substitution at position 54, achieve improved affinities predominantly through slower dissociation rates^44,46,50^.

eN49P7-FRv1-23 acquired 15 VH mutations and 11 VL mutations during the *in vitro* affinity maturation of N49P7-FR. Notably, many of these mutations (12 in VH and 3 in VL) are also found in other VRC01-class bnAbs (Supplementary Fig. 11), highlighting the different maturation trajectories a bnAb can take to achieve high affinity binding. These mutations neither induced polyreactivity (Supplementary Fig. 22A,B) nor impaired SEC monodispersity (Supplementary Fig. 22C), thermal stability (Supplementary Fig. 22D), or solubility (Supplementary Fig. 22E). While they slightly increased hydrophobicity, it remained within the range observed for clinical antibodies (Supplementary Fig. 22F). To assess pharmacokinetics (PK), we evaluated the plasma half-life (t_1/2_) of eN49P7-FRv1-23 in homozygous human neonatal Fc receptor (hFcRn) mice, a well-established model for predicting the t_1/2_ of human antibodies, which correlates more closely with human PK than wild-type mice, hemizygous hFcRn mice, or non-human primates^51^. The Fc region of eN49P7-FRv1-23 was modified with the LS mutations (M428L, N434S), which enhance binding affinity to hFcRn and thereby extend plasma t_1/2_^52,53^. Following intraperitoneal administration, eN49P7-FRv1-23-LS had a plasma t_1/2_ of 10 days (Supplementary Fig. 22G), comparable to that of previously tested VRC01-LS^36^. While this was shorter than the half-lives of N49P7-LS (14 days) and N49P7-FR-LS (12 days), it exceeded that of VRC07-523-LS (6 days), a long-acting clinical bnAb candidate for HIV prevention and treatment^36,54,55^. These findings indicate that eN49P7-FRv1-23 maintains a favorable PK profile comparable to leading clinical candidates, despite the extensive mutations acquired to enhance potency and breadth.

### 3. Structural analysis of the optimized N49P7-FR

To understand the enhanced activity of eN49P7-FRv1-23 at the atomic level, we generated cryo-electron microscopy (cryo-EM) structures of N49P7-FR and eN49P7-FRv1-23 Fab, each in complex with BG505 MD39 SOSIP^56^, with estimated global reconstruction resolutions of 2.7 Å and 2.6 Å, respectively (Fig. 4A, Supplementary Fig. 23, and Supplementary Table 10). Overall, the Fv regions of the two antibodies have high overlap, with a peptide backbone Root Mean Square Deviation (RMSD) of 1.0 Å with respect to gp120 alignment (Fig. 4B). In addition, the FRH3 insertion in the heavy chain extends to the neighboring gp120 protomer in both structures (Fig. 4C). A D76bY^eVH^ mutation in eN49P7-FRv1-23 FRH3 results in a hydrogen bond (HB) with P118^Env^ (estimated distance 2.8 Å) while also improving packing through its bulkier side chain. The non-mutated D76b^VH^ in N49P7-FR is nearby potential HB acceptors/donors in the backbones of P206-K207^Env^ but with distances ∼3.3 Å, suggesting weaker or transient interactions (Fig. 4C).

**Figure 4.**
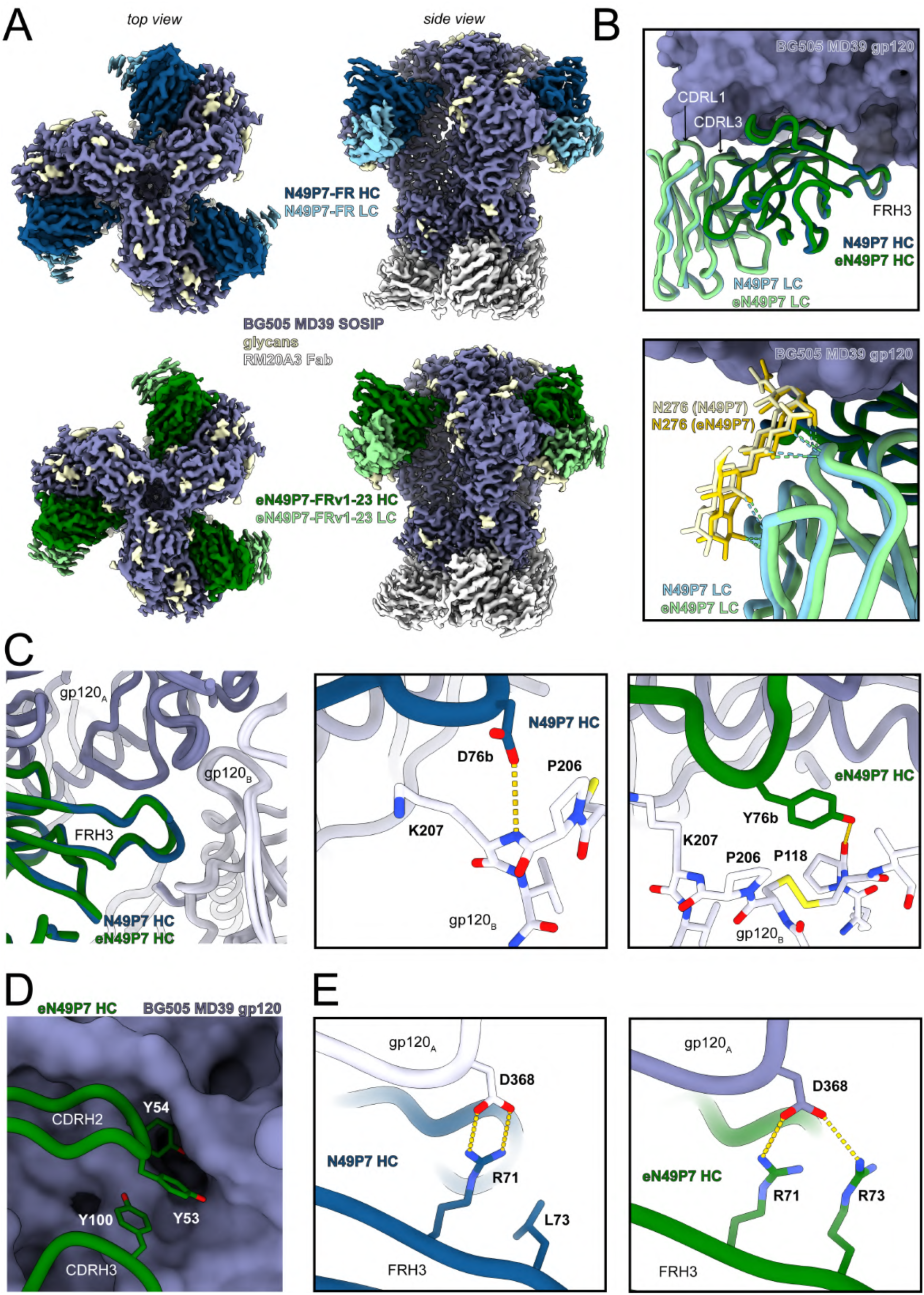
Structural analysis of eN49P7-FRv1-23 mutations via cryo-EM. **A**) Cryo-EM reconstructions, colored by component, of BG505 MD39 SOSIP in complex with N49P7-FR (*upper panels*) or eN49P7-FRv1-23 (*lower panels*). **B**) Overlay of N49P7-FR (N49P7) and eN49P7-FRv1-23 (eN49P7) Fv regions after gp120 alignment (*upper panel*). The backbone Root Mean Square Deviation (RMSD) between the two antibodies is 1.0 Å. Interaction of the light chains of each antibody with Env gp120 N276 glycan (*Iower panel*). Putative hydrogen bonds are depicted as dashed lines. **C**) Overlay of the FRH3 regions of each antibody, which includes an insertion that extends to the adjacent gp120 (chain B). Specific interactions of each antibody are shown in the middle and right panels with a focus on the D76bY mutation. Putative hydrogen bonds are shown as dashed lines. **D**) Three tyrosine mutations (Y53, Y54, and Y100) in eN49P7-FRv1-23 insert into a groove in gp120. **E**) Comparison of the conserved D368 salt bridge with FRH3 residue R71, and the L73R mutation in eN49P7-FRv1-23. Amino acid positions are numbered according to the Kabat scheme.

Using a 3.4 Å distance cutoff criteria for HB and considering only the primary gp120 protomer, both N49P7-FR and eN49P7-FRv1-23 heavy chains are predicted to form 10 HB with BG505 MD39 SOSIP peptides, while the light chains contribute only 1-2 each via the conserved “YE” motif in CDRL3. The light chains also form multiple HB with the sugars of the N276^Env^ glycan but involve residues that are conserved between the two antibodies and therefore the increased potency of eN49P7-FRv1-23 is not thought to be due to glycan-specific contacts (Fig. 4B).

Focusing on the heavy chain, Y54^eVH^ was the most enriched mutation in mutagenesis scanning across all gp120-sorted populations (Fig. 2A, Supplementary Fig. 8). The presence of Y54^eVH^ strongly favored the M53Y^eVH^ mutation (Fig. 2C, Supplementary Fig. 10). Combined with the side chain of Y100^eVH^, which was also enriched for high affinity binding to a subset of gp120s in mutagenesis scanning (Fig. 2A, Supplementary Fig. 8), the three mutant tyrosines in eN49P7-FRv1-23 (Y53^eVH^, Y54^eVH^, and Y100^eVH^) spatially occupy a void in the hydrophobic cavity of the Env CD4bs; their aromatic rings stack with one another to form a bulky triad that fills up the CD4bs pocket, thus adding to the cumulative increase in the interaction strength (Fig. 4D). R71^(e)VH^ in FRH3 of both bnAbs forms a salt bridge with D368^Env^ (Fig. 4E). Importantly, an L73R^eVH^ mutation in eN49P7-FRv1-23 provides a second salt bridge with D368^Env^, further strengthening the interaction (Fig. 4E). This mutation was highly enriched across most of the gp120-sorted populations (Fig. 2A, Supplementary Fig. 8). D368^Env^ and Q428^Env^, which are engaged by the heavy chain of eN49P7-FRv1-23 through salt bridges and/or a HB, are highly conserved across HIV clades, as indicated by sequence analysis of Envs from the 206 pseudoviruses used in our *in vitro* neutralization assay (Supplementary Table 11). Among these, 26 pseudoviruses contained no mutations at any of the identified bnAb contact positions and were neutralized by eN49P7-FRv1-23 (IC_80_ <4 μg/mL), though some remained resistant to N49P7-FR. Additionally, most pseudoviruses neutralized by eN49P7-FRv1-23 harbored mutations at some of these contact positions, while several strains with identical mutations exhibited divergent neutralization outcomes (e.g., 3301.v1.c24 was sensitive, whereas ZM233M.PB6 was resistant to eN49P7-FRv1-23, despite both carrying the N279D^Env^ and A281V^Env^ mutations). These observations suggest the involvement of yet-undetermined additional interactions and/or structural features at the Env/bnAb interface that contribute to viral sensitivity.

A canonical feature of VRC01-class antibodies, the conserved “YE” motif in CDRL3^57^, maintains crucial contacts via HB to the gp120 D-loop (residues N279^Env^ and/or N280^Env^) and the backbone of G459^Env^ in gp120 V5 (Supplementary Fig. 24A). In addition, an unpaired cysteine at position 36 in FRL2 of N49P7-FR was mutated to leucine. This mutation was highly enriched across all gp120-sorted populations (Fig. 2B, Supplementary Fig. 9). It may enable improved packing of the hydrophobic core, being surrounded by the hydrophobic side chains of L46^eVL^, W87^eVL^, F89^eVL^, F98^eVL^, L100k^eVH^, and W103^eVH^ (Supplementary Fig. 24B).

To further characterize the effect of the introduced mutations in eN49P7-FRv1-23 on the binding interface, we performed molecular dynamics (MD) simulations to capture differences in binding site flexibility and interactions (direct and water-mediated) (Fig. 5). Overall, eN49P7-FRv1-23 shows more favorable electrostatic and van der Waals interaction energies than N49P7-FR, accompanied by a rigidification of the binding interface. In particular, the CDRH3 loop exhibits reduced flexibility.

**Figure 5.**
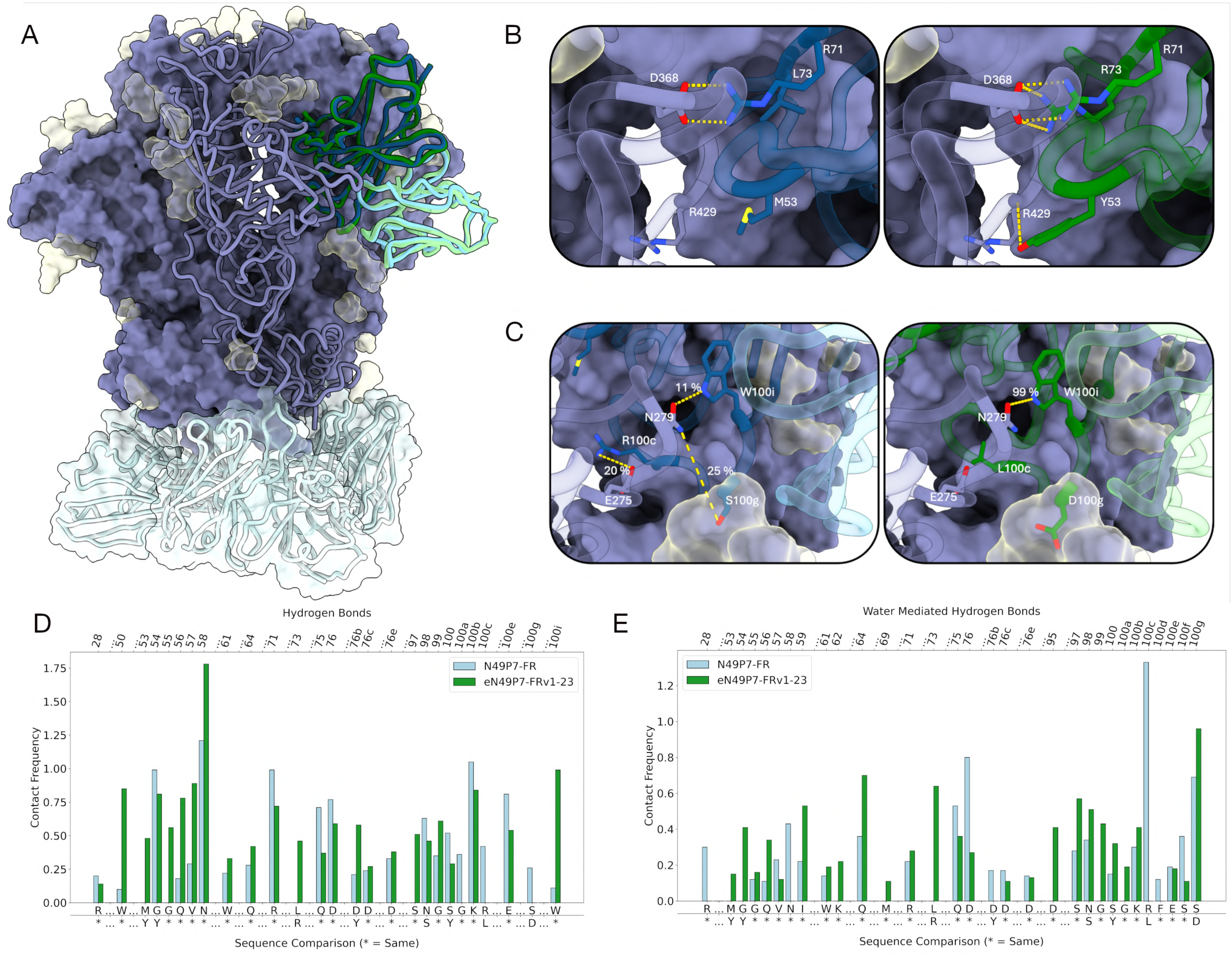
MD simulations of eN49P7-FRv1-23 mutations. **A**) Overlay of the cryo-EM starting structures for MD simulations (PDB accession codes: 9P6E and 9P6G). N49P7-FR and eN49P7-FRv1-23 are shown in blue and green, respectively. **B–C**) Changes in interface interactions and contact frequencies resulting from the introduced mutations. **D–E**) Hydrogen bond and water-mediated hydrogen bond frequencies for the paratope residues quantifying similarities and differences between N49P7-FR and eN49P7-FRv1-23. Amino acid positions are numbered according to the Kabat scheme.

Mutations such as M53Y^eVH^ in CDRH2 and L73R^eVH^ in FRH3 introduce new charged HB and salt bridge interactions that directly strengthen the binding interface (Fig. 5B). M53Y^eVH^ forms HB with W427^Env^ and R429^Env^, while L73R^eVH^ forms long-lasting HB and salt bridge interactions with Q428^Env^ and D368^Env^, respectively. A critical residue in the binding interface, W100i^(e)VH^, makes hydrophobic interactions in both N49P7-FR and eN49P7-FRv1-23, but is substantially more stabilized in eN49P7-FRv1-23, allowing for a long-lasting HB (99% of the simulation time) with N279^Env^ (Fig. 5C,D). This stabilization is facilitated by surrounding CDRH3 mutations, R100cL^eVH^ and S100gD^eVH^. R100cL^eVH^ disrupts a transient salt bridge with E275^Env^ (20% of the simulation time) but reduces repulsion with K282^Env^ and displaces water from the binding site (Fig. 5E), collectively contributing to interface stabilization. In N49P7-FR, S100g^VH^ forms a HB with N279^Env^, in addition to water-mediated contacts with the N276^Env^ glycan and intramolecular interactions with D50^VL^ of the light chain. By contrast, S100gD^eVH^ forms stronger, sustained interactions with both the N276^Env^ glycan and N279^Env^, stabilizing the glycan and thereby the binding site. The rigidification of S100gD^eVH^ and W100i^eVH^ is reflected in B-factor analysis (196 Å² → 124 Å² / 195 Å² → 118 Å²), indicating reduced flexibility, and in cluster analysis of the antibody binding site, which reveals fewer conformations (Supplementary Fig. 25). Cluster analysis of the N276^Env^ glycan also shows reduced flexibility, evidenced by a markedly lower number of clusters.

## Discussion

The clinical utility of HIV bnAbs relies critically on their ability to potently neutralize diverse circulating viral variants. Developing antibodies with the necessary breadth and potency is inherently challenging due to the extreme antigenic variability of HIV Env^9,13^. bnAbs isolated from infected individuals evolve under multiple constraints: exposure to only a fraction of circulating viral diversity and intrinsic B-cell programs, whereby germinal center kinetics and plasma cell differentiation impose a functional ceiling on binding affinities^28,29,30,32,33^. *In vitro* affinity maturation offers a powerful strategy to overcome these limitations by enabling systematic and targeted exploration of antibody sequence space against a broader spectrum of antigenic diversity and applying selective conditions designed to more effectively enrich high-affinity antibodies^58^.

In this study, we established a robust yeast surface display-based platform for multistate optimization of bnAbs targeting HIV Env (Fig. 1). Leveraging dbSMS, we comprehensively defined the mutational landscape of the CD4bs bnAb N49P7-FR, identifying amino acid substitutions that significantly enhanced gp120 binding (Fig. 2). Crucially, introducing molecular barcodes into the dbSMS libraries enabled precise normalization of mutational enrichment data, greatly improving variant selection accuracy by distinguishing true enrichments from sequencing errors and normalizing relative to starting residues. This approach also yielded relatively compact initial libraries, facilitating parallel selections against a diverse panel of gp120 variants and enabling identification of universally beneficial mutations rather than those improving binding in a strain-specific manner.

The strongest mutational enrichments were observed in the heavy chain (Fig. 2A,C), consistent with the binding mode of VRC01-class bnAbs. Many of the substitutions identified in our scans mirrored mutations found in naturally occurring VRC01-class bnAbs, reinforcing their functional relevance. Enrichment within the grafted FRH3 insertion from VRC03 further highlighted the need for context-specific optimization: while this loop was advantageous in its native framework, its transfer to N49P7 required additional adaptations for stable integration (Fig. 2A).

Combinatorial libraries then allowed us to explore optimal combinations of these mutations by sampling across diverse gp120s in parallel (Supplementary Fig. 12). This strategy enabled us to distinguish broadly beneficial substitutions from those that were strain-restricted, while reducing, though not eliminating, the number of variants with unfavorable biophysical properties through integrated negative selections. In practice, many candidates still required small-scale IgG production and screening to confirm potency and biochemical behavior. By systematically combining enriched and neutral substitutions, we could traverse multiple potential evolutionary trajectories and identify variants that maximized breadth and potency without sacrificing stability. This approach ultimately produced antibody variants with substantially improved neutralization profiles and favorable developability characteristics.

The final engineered variant, eN49P7-FRv1-23, demonstrated substantially improved neutralization potency and breadth against a large panel of HIV pseudoviruses (Fig. 3A), validating the efficacy of our multistate optimization strategy. Structural analyses revealed that the enhanced potency primarily arose from novel and stabilized interactions at the antibody/Env interface, particularly a distinctive tri-tyrosine aromatic wedge (Y53/Y54/Y100) penetrating deeply into the conserved hydrophobic cavity of the CD4bs (Fig. 4D). Notably, this tri-aromatic wedge represents a unique structural adaptation not previously observed in other VRC01-class antibodies, whether naturally derived or engineered^43,44,47,59,60,61,62^. Molecular dynamics simulations provided additional mechanistic insights, indicating that affinity-enhancing mutations improved electrostatic and van der Waals interactions and substantially rigidified key loops, notably the CDRH3, stabilizing critical antibody–Env contacts (Fig. 5B,C). Despite extensive sequence modifications, eN49P7-FRv1-23 retained excellent biochemical characteristics, including low polyreactivity, high thermal stability, and favorable developability metrics comparable to clinical-stage antibodies (Supplementary Fig. 22).

The ability to systematically engineer bnAbs that combine high potency, breadth, and robust developability opens new opportunities for HIV therapeutics. In the context of cure-focused strategies, such antibodies could strengthen immune clearance of latent reservoirs and synergize with small-molecule regimens to achieve lasting control^20,63,64,65,66,67,68^. Although ebnAbs could have potential as prophylactic agents, long-acting antiretrovirals have set a high standard for cost, efficacy and ease of delivery, and will almost certainly remain the first-line option^69,70,71,72^. Nevertheless, bnAbs with expanded coverage, such as those described here, could pave the way for truly effective alternatives in circumstances where ART is difficult to deploy, including prevention of mother-to-child transmission and other specialized contexts^9,10,11^.

In summary, our findings establish that systematic, multistate optimization can yield HIV bnAbs with markedly improved potency, breadth, and favorable drug-like properties. The generation of eN49P7-FRv1-23 illustrates the practical value of this approach, achieving neutralization performance that substantially exceeds its parental lineage. More broadly, the platform we describe offers a generalizable framework for mapping and assembling optimal mutational solutions across heterogeneous antigens, providing a versatile path toward next-generation antibody therapeutics.

## Acknowledgements

This work was supported by Gates Foundation grants: INV-005284 to M.M.S. and INV-036842 to M.S.S.; NIH grants: R01AI147870 to M.M.S., 1R01AI155150-01A1 to M.M.S. and A.L.D., and R35 GM144088 to B.G.P. We thank the Genomics Core Facility at Scripps Research (La Jolla, CA, USA) for deep sequencing using Illumina platforms, and Christopher A. Cottrell (Department of Immunology and Microbiology, Scripps Research, La Jolla, CA, USA) for providing HIV Env proteins.

## Data Availability

Cryo-EM maps have been deposited to the Electron Microscopy Data Bank (EMDB) under accession codes EMD-71307 and EMD-71308, and associated atomic models have been deposited to the Protein Data Bank (PDB) under accession codes 9P6E and 9P6G.

## Author Contributions

M.Kę.: designed and executed antibody optimization via *in vitro* affinity maturation; performed deep sequencing and bioinformatical data processing; performed IgG1 variant production, purification, and characterization (functional: *in vitro* neutralization, polyreactivity; biochemical: analytical SEC, HIC, DSF, PEG solubility); analyzed and interpreted data; contributed to writing the manuscript. S.P. and G.O.: performed cryo-EM; analyzed and interpreted structural data; contributed to writing the manuscript. M.F.-Q. and J.L.: performed MD simulations; analyzed and interpreted simulations data; contributed to writing the manuscript. C.J.: performed bioinformatical processing of deep sequencing data. J.W.: performed SPR; analyzed and interpreted binding kinetics data; assisted with analytical SEC. A.Ab., M.Ka., A.As., M.H., R.Z., B.A., E.F.P.B., and A.H.: performed PK studies in humanized mice; analyzed and interpreted data. K.S.-F.: produced HIV gp120 proteins. Q.T.: cultured mammalian cells and produced Fab fragments. Y.K.: performed IgG1 production and endotoxin-testing. F.A.-P.: performed IgG1 production. L.S. and A.Z.: cultured mammalian cells, confirmed HIV pseudovirus Env DNA sequences, and produced pseudovirus backbone DNA. P.X.A.: produced CHO-SMP, biotinylated PSR, and HIV Env proteins. N.R.F. and B.G.P.: performed structure-guided designing of VH variants. D.S.: provided input on the manuscript. M.S.S.: provided a global cross-clade panel of HIV pseudoviruses. D.R.B.: provided input on the manuscript; A.B.W.: supervised and provided input on cryo-EM and MD simulations. A.L.D.: provided input on studies and on the manuscript. M.M.S.: supervised and provided input on studies and on the manuscript. J.G.J.: conceived, designed, supervised, and provided input on antibody optimization via *in vitro* affinity maturation; provided input on antibody functional and biochemical characterization; interpreted data; ensured financial support; contributed to writing the manuscript.

## Materials and Methods

### Optimization of N49P7-FR

A double-barcoded saturated mutagenesis scanning (dbSMS) library was generated for both VH and VL regions of N49P7-FR. This library was synthesized by Twist Bioscience, where each amino acid position was individually mutated by replacing the original codon with a degenerate codon encoding all amino acids except cysteine to avoid creating additional disulfide bonds. Silent “barcode” mutations were introduced into adjacent codons flanking the degenerate codon to enable precise identification of amino acid substitutions during sequencing. For amino acids encoded by a single codon (e.g., methionine or tryptophan), barcode mutations were shifted to neighboring codons.

The linearized yeast display vector pYDSI2u (with a bidirectional Gal1-10 promoter and URA3 marker)^73^ and the dbSMS library, containing ∼100 bp flanking regions for overlap with the vector, were co-transformed into *Saccharomyces cerevisiae* YVH10 cells (ATCC, cat. no. MYA-4940) by electroporation, as previously described^74^. Constructs were assembled via homologous recombination in yeast. Fab fragments of N49P7-FR variants were displayed on the yeast cell surface, fused to a synthetic mucin-like domain and GPI anchor via the CH1 domain. The CH1 domain was tagged with V5, and the Cλ domain was tagged with c-Myc for detection.

Yeast cultures were grown in selective SD-Ura medium (Sunrise Science Products, cat. no. 1703) and induced in SGCAA+Trp medium, prepared by dissolving 20 g galactose, 1 g glucose, 6.7 g Difco yeast nitrogen base without amino acids, 5 g Bacto casamino acids, 5.4 g sodium phosphate dibasic, 8.6 g sodium phosphate monobasic monohydrate, and 85.6 mg L-tryptophan in DI water to a final volume of 1 L. Yeast cells displaying Fab fragments were incubated in parallel with His-tagged gp120 proteins from diverse viral clades (92BR020, 25710, 93TH057, IAVI-C22, BJOX2000, and CH119) at non-saturating concentrations. Cells were labeled with anti-c-Myc-FITC (ICL, cat. no. CMYC-45F), anti-V5-AF405 (produced in-house), and anti-His-APC (Miltenyi Biotec, cat. no. 130-119-782) antibodies. A BD FACSMelody™ cell sorter was used to select the top 5–10% gp120 binders (∼50,000 cells) and the top 5–10% Fab-displaying cells not exposed to gp120 (∼500,000 cells as a control). Sorted cells were cultured in SD-ura medium, induced in SGCAA+Trp medium, and subjected to another round of sorting to further enrich for variants with improved gp120 binding (Supplementary Figs. 1 and 2). All library screening steps were performed in duplicate.

Display vectors were then isolated from the sorted and grown yeast, and antibody-encoding DNA was PCR-amplified and indexed using Nextera XT v2 adapters. Libraries were pooled and sequenced on the Illumina MiSeq system with a 600-cycle v3 reagent kit (cat. no. MS-102-3003). Paired-end reads were merged, trimmed, translated, and analyzed to exclude sequences with stop codons, insufficient length, multiple mutations, or incorrect barcoding. Remaining reads were counted by position and amino acid type. Enrichment of amino acid substitutions in top gp120-binding variants relative to the control dataset was calculated:

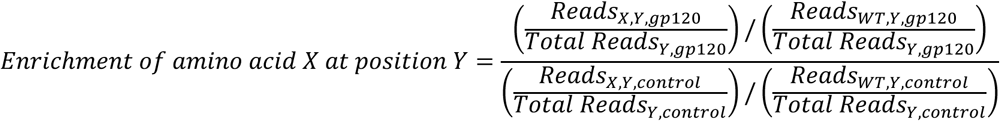

Where:

- Reads_X,Y,gp120_ represents the number of reads for amino acid X at position Y in the gp120 dataset,
- Total Reads_Y,gp120_ represents the total number of reads at position Y in the gp120 dataset,
- Reads_WT,Y,gp120_ represents the number of reads for the wild-type amino acid (WT) at position Y in the gp120 dataset,
- Similarly, Reads_X,Y,control_, Total Reads_Y,control_, and Reads_WT,Y,control_ represent the corresponding values for the control dataset.

Using these enrichment data, combinatorial libraries for VH and VL were constructed. Positions with highly enriched mutations (>3-fold) were targeted, and degenerate codons encoding favorable amino acids, including wild-type residues, were introduced at 16 VH and 15 VL positions (Supplementary Fig. 12A,B). Each library was synthesized by Integrated DNA Technologies (IDT) as overlapping ultramers, which were PCR-assembled and combined with the linearized display vector in yeast after electroporation as described above. The theoretical library sizes were 2.2 x 10^6^ for VH and 3.3 x 10^6^ for VL, with ∼3 x 10^8^ and ∼1 x 10^8^ yeast clones obtained for VH and VL libraries, respectively, providing >100-fold and >30-fold coverage of their theoretical diversity.

The VH combinatorial library was expressed on yeast with the wild-type VL and screened for improved gp120 binding. Subsequently, the VL combinatorial library was expressed with one of the most enriched VH variants (VHv10), which exhibited increased potency as an IgG1 in HIV neutralization assay. Both libraries underwent five and six rounds of selection, respectively, alternating positive selections with His-tagged gp120s and negative selections with biotinylated polyspecificity reagent (PSR)^75^ to eliminate polyreactive variants (Supplementary Figs. 4 and 5). Positive selections used 12 gp120s (92BR020, 25710, 93TH057, IAVI-C22, BJOX2000, CH119, 398F1, CAP45, 94UG103, 6041.v3, T278-50, X1632) at non-saturating concentrations, including resistant strains to increase bnAb breadth, collecting the top 5-10% gp120 binders (∼50,000 cells). Negative selections were performed with membrane-derived PSR at 5 μg/mL or cytosolic PSR at 50 μg/mL, collecting the bottom 20-25% PSR binders (∼50,000 cells).

Sorted yeast cells were grown, display vectors isolated, and DNA sequenced. Enriched variants were ranked by median frequency across datasets. Select VH (24) and VL (21) variants were reformatted and expressed as IgG1 in Expi293F cells (described below); VH variants were paired with the wild-type VL and, subsequently, VL variants with one of the most enriched VH variants, VHv10. These variants were evaluated for neutralization potency and biochemical properties as described below (‘Antibody characterization’). Based on these assessments, the most potent VH variant, VHv1, was paired with VLv3 for further HC optimization.

A dbSMS library was generated for VHv1 to improve its biochemical properties while maintaining enhanced neutralization potency. Variants were expressed on yeast, labeled, and sorted for gp120 binding (Supplementary Fig. 3). Additional selections were performed to identify thermally stable variants by preincubating the library at 65 °C for 1 hour prior to gp120 binding and sorting. Indexed libraries were sequenced on the Illumina NextSeq2000 system with a 600-cycle P1 reagent kit (cat. no. 20075294). Enrichment data guided construction of two VHv1 combinatorial libraries sampling 21 and 28 positions (Supplementary Fig. 12C), with theoretical sizes of 9.4 x 10^6^ and 1.2 x 10^9^. Approximately 6 × 10^7^ yeast clones were obtained per each library transformation, providing >6-fold and ∼5% coverage of theoretical diversity, respectively. The libraries were screened in six selection rounds alternating positive gp120 binding and negative PSR sorting, including thermal challenges as described above (Supplementary Fig. 6). DNA from sorted libraries was sequenced, and enriched variants ranked. Ninety-six most enriched variants with the highest median frequency across datasets were evaluated as IgG1, identifying VHv1-23/VLv3 as the final optimized candidate with superior neutralization potency and developability.

### Bioinformatical analysis

The deep sequencing datasets (FASTQ files) generated from the sorted dbSMS libraries and combinatorial libraries were parsed and analyzed bioinformatically as described above (‘Optimization of N49P7-FR’). Briefly, paired-end MiSeq and NextSeq reads were merged using PANDAseq^76^. The merged sequences were subsequently processed using custom Python 3.8 scripts (https://github.com/jardinelab/Multistate-optimization-of-HIV-bnAbs), which performed the following steps: removal of primers and low-quality reads, filtering of sequences with insertions or deletions, translation of nucleotide sequences, exclusion of variants containing stop codons or unintended amino acid substitutions, and counting of remaining variants. The amino acid enrichment at each position in dbSMS libraries (calculated using the equation provided in the section ‘Optimization of N49P7-FR’) and the variant frequency in combinatorial libraries were then computed.

### Protein production

#### gp120 production

Each gp120 protein was produced in a suspension culture of FreeStyle™ 293F cells (Invitrogen, cat. no. R79007) through transient transfection with a pHL-sec plasmid^77^ (a gift from Edith Yvonne Jones; Addgene plasmid #99845) using 293fectin™ (Invitrogen, cat. no. 12347019) as the transfection reagent. The plasmid encoded a mammalian codon-optimized gp120 protein with a C-terminal 6x His affinity tag. After 96 hours, gp120 proteins were harvested from the supernatants and purified by affinity chromatography using a HisTrap™ HP column (Cytiva, cat. no. 29051021), followed by size-exclusion chromatography with a Superdex™ 200 Increase 10/300 GL column (Cytiva, cat. no. 28990944) on an ÄKTAxpress chromatography system (Cytiva).

#### Antibody production

Each IgG1 was produced in a suspension culture of Expi293F™ cells (Invitrogen, cat. no. A14527) through transient co-transfection with two mammalian expression vectors encoding the heavy and light chains (1:2.5 ratio of HC:LC DNA), synthesized by GenScript Biotech. FectoPRO^®^ (Polyplus-transfection, cat. no. 116-040) was used as the transfection reagent. After 24 hours, the cultures were supplemented with 300 mM valproic acid (Sigma-Aldrich, cat. no. P4543) and 45% D-glucose (Sigma-Aldrich, cat. no. G8769). After additional 96 hours, antibodies were harvested from the supernatants and purified by affinity chromatography using Praesto^®^ AC agarose resin (Purolite, cat. no. PR000200-164) or, for high-throughput purification, Pierce™ protein A/G magnetic agarose beads (Thermo Scientific, cat. no. 78610). Antibodies were eluted from the chromatographic resin using a low-pH citrate buffer (9.38 mM sodium citrate dihydrate, 90.62 mM citric acid monohydrate, 150 mM sodium chloride, pH 3.0), followed by a 1-hour room-temperature incubation for low-pH viral inactivation—a routine step for antibody-based therapeutics to ensure viral clearance. The eluates were neutralized with 2 M Tris, pH 10.0, buffer-exchanged into PBS, centrifuged, visually inspected for precipitation, and quantified by absorbance at 280 nm using a NanoDrop One spectrophotometer (Thermo Scientific). Small aliquots (5 μg) were analyzed by SDS-PAGE to confirm the proper assembly of the heavy and light chains.

### Antibody characterization

#### Analytical size-exclusion chromatography (SEC)

Control and test antibodies (20 µg) in PBS were prepared in a 96-well plate (Corning, cat. no. 3788) and analyzed for retention time using the 1260 Infinity II LC System (Agilent). Samples were loaded onto a 30 cm TSKgel SuperSW mAb HR column (Tosoh, cat. no. 22854). PBS was used as the mobile phase, applied for 20 minutes at a flow rate of 1 mL/min. UV absorbance was monitored at 280 nm. Adalimumab, ipilimumab, and golimumab—FDA-approved antibodies with favorable physicochemical properties—were included as positive controls, while bococizumab, known for its aggregation-prone behavior, was included as a negative control. Control antibodies were produced in-house using Expi293F cell cultures. The experiment was performed in duplicate.

#### Polyspecificity reagent (PSR) ELISA

Solubilized CHO cell membrane proteins (CHO-SMP) were prepared in-house following the previously published protocol^75^. CHO-SMP, human insulin (Sigma-Aldrich, cat. no. I2643), and single-stranded DNA (ssDNA, Sigma-Aldrich, cat. no. D8899) were coated onto 96-well half-area high-binding plates (Corning, cat. no. 3690) at a concentration of 5 μg/mL in PBS and incubated overnight at 4 °C. After three washes with PBS containing 0.05% Tween 20 (PBST), the plates were blocked with PBS containing 3% BSA for 1 hour at room temperature (RT). Test and control antibodies were diluted to a concentration of 50 μg/mL in PBS containing 1% BSA, followed by additional 4-fold serial dilutions. The diluted antibodies were added to the blocked plates and incubated for 1 hour at RT. After three washes with PBST, an alkaline phosphatase-conjugated goat anti-human IgG Fcγ secondary antibody (Jackson ImmunoResearch, cat. no. 109-055-008), diluted 1:1000, was added and incubated for 1 hour at RT. Following the final three washes with PBST, phosphatase substrate (Sigma-Aldrich, cat. no. S0942–200TAB), reconstituted in staining buffer (10 mM magnesium chloride hexahydrate, 100 mM sodium bicarbonate, pH 9.8), was added to each well, and absorbance was measured within 20 min at 405 nm using a Synergy H1 microplate reader (BioTek). Adalimumab, a non-polyreactive antibody, and 4E10, a highly polyreactive antibody, were included as controls. Both were produced in-house using Expi293F cell cultures. Each antibody sample was tested in duplicate.

#### HEp-2 cell reactivity

Reactivity to human epithelial type 2 (HEp-2) cells was assessed by indirect immunofluorescence using HEp-2 slides (Hemagen, cat. no. 902360) according to the manufacturer’s instructions. Briefly, test antibodies were diluted to 50 μg/mL in PBS, applied to the wells on the slides containing immobilized HEp-2 cells, and incubated for 30 minutes at RT. After washing with PBS, one drop of FITC-conjugated goat anti-human IgG was added to each well on the slides and incubated in the dark for 30 minutes at RT. Following another wash, a drop of glycerol was added to each well, the slides were covered with coverslips, and images were captured using an EVOS f1 fluorescence microscope for FITC detection at 40x magnification. Reactivity to HEp-2 cells was interpreted based on the observed staining patterns. Positive and negative control sera, provided by the manufacturer, were included in the assay. The experiment was performed in duplicate.

#### Differential scanning fluorimetry (DSF) – thermal shift

The thermal stability of test antibodies was evaluated by determining their melting temperature (T_m_) using a real-time PCR system. Test antibodies (18 μL at 0.5 mg/mL) were mixed with 2 μL of 10x GloMelt™ dye (Biotium, cat. no. 33021-1) and transferred to a 96-well skirted PCR plate (Bio-Rad, cat. no. HSP9601), which was sealed with an optical adhesive film. Melt curves were generated using the CFX96 Touch Real-Time PCR System (Bio-Rad, cat. no. 1855195) equipped with CFX Maestro Software. Fluorescence was detected using the SYBR^®^ Green channel as the temperature was incrementally increased from 25 °C to 99 °C in 0.5 °C steps, with a 10-second hold at each step. T_m_ values were calculated from the peaks of the melt curves, derived from the first derivatives of the fluorescence intensity plotted against temperature. Antibody samples were tested in triplicate.

#### Solubility with polyethylene glycol (PEG)

PEG is a precipitant that lowers the intrinsic solubility maxima of proteins, enabling the prediction of antibody solubility in highly concentrated solutions by analyzing them at low concentrations. Antibody solubility was assessed using PEG as described previously^78,79^, with modifications. Briefly, 25 μL of test and control antibodies (1 mg/mL) were mixed with 75 μL of PEG solution in PBS to achieve a final PEG concentration of 8% (w/v). Blank samples for each antibody were prepared by adding PBS instead of PEG solution. Samples were incubated for 1 hour at RT in a 96-well non-treated (medium-binding) flat-bottom plate (Corning, cat. no. 9017) and pipetted up and down several times to resuspend any possible precipitate. Optical density (OD) was immediately measured at 350 nm using a Synergy H1 microplate reader (BioTek). Antibodies were predicted to be soluble at high concentrations (>50 mg/mL) if their OD_350_ values in PEG-containing samples did not increase by more than 0.03 compared to corresponding blank samples, as an OD_350_ increase of 0.03 indicates the onset of visible precipitation. Adalimumab (highly soluble antibody) and bococizumab (poorly soluble antibody) were used as controls. Each antibody sample was tested in duplicate.

#### Hydrophobic interaction chromatography (HIC)

Control and test antibodies (20 µg) in PBS were prepared in a 96-well plate (Corning, cat. no. 3788) and analyzed for retention time using the 1260 Infinity II LC System (Agilent). Samples were loaded onto a 10 cm TSKgel Butyl-NPR column (Tosoh, cat. no. 42168). A linear salt gradient was applied, transitioning from mobile phase A (1.5 M ammonium sulfate, 50 mM sodium phosphate, pH 7.0, 5% v/v isopropanol) to mobile phase B (50 mM sodium phosphate, pH 7.0, 20% v/v isopropanol) over 15 minutes at a flow rate of 0.5 mL/min. UV absorbance was monitored at 280 nm. Adalimumab, an FDA-approved antibody, and 4E10, a highly hydrophobic antibody, were included as controls. The experiment was performed in duplicate.

#### Pseudovirus production and neutralization

To produce HIV Env-pseudotyped viruses, plasmids encoding Env were co-transfected with an Env-deficient genomic backbone plasmid (pSG3ΔEnv) at a 1:2 ratio using the transfection reagent PEI MAX^®^ (Polysciences, cat. no. 24765) into 293T cells (ATCC, cat. no. CRL-3216). Pseudoviruses were harvested 72 hours post-transfection and used in a luciferase-based assay to assess the neutralizing activity of test antibodies. This assay measures a single round of pseudovirus replication in TZM-bl target cells (NIH AIDS Reagent program, cat. no. 8129), as described previously^80,81,82^. Neutralization results were reported as IC_80_ values, which indicate the antibody concentrations required to achieve 80% neutralization of HIV infection in TZM-bl cells. All antibody samples were tested in duplicate.

#### Surface plasmon resonance (SPR)

SPR measurements were carried out on a Biacore 8K instrument at 25 °C. All experiments were performed at a flow rate of 30 μL/min in a mobile phase of HBS-EP+ (0.01 M HEPES, pH 7.5, 0.15 M NaCl, 3 mM EDTA, 0.0005% (v/v) Surfactant P20) supplemented with 1 mg/mL BSA. For conventional kinetic/dose-response measurements, IgG1 samples were captured at 50 to 100 resonance units (RU) on a Series S Sensor Chip Protein A (Cytiva, cat. no. 29127556) before analyte injection. A dilution series of gp120 molecules was injected over the IgG1 and control surfaces for 2 min, followed by a 20-min dissociation phase using a multi-cycle method. The surface was regenerated between cycles with three 120 s injections of 10 mM glycine (pH 1.7). Kinetic analysis of reference-subtracted injection series was conducted using BIAEvaluation software (Cytiva). Sensorgram series were fit to either a 1:1 (Langmuir) binding model or a heterogeneous ligand model.

#### Pharmacokinetics (PK) and plasma half-life (t_1/2_) in hFcRn transgenic mice

Animal studies were approved by the University of Maryland School of Medicine IACUC. PK analysis and t_1/2_ determination were performed in homozygous human FcRn transgenic mice (FcRn⁻/⁻ hFcRn⁺/⁺, Jackson Laboratory), which lack the murine FcRn α-chain (Fcgrt^tm1Dcr^) and instead express the human FcRn α-chain. This model has been extensively validated for evaluating the *in vivo* stability of human IgG antibodies, including Fc-engineered variants, and shows strong correlation with human PK outcomes^51^. Mice (9 weeks old; similar numbers of males and females; n = 4–7 per group) received a single intraperitoneal injection of 10 mg/kg of the test antibody. Blood samples were collected at days 1, 3, 7, 14, 21, 28, 35, and 42 post-injection. Plasma concentrations of the test antibodies were quantified using a human IgG-specific ELISA. Terminal half-life estimates were calculated using non-compartmental analysis (NCA), following established PK modeling principles. Briefly, the elimination rate constant (λz) was derived from the log-linear phase of the concentration-time profile by performing linear regression of log-transformed plasma concentrations versus time. All available data points from day 7 onward were used to define the terminal phase, and values below the assay’s lower limit of quantification were excluded from the analysis. Individual t_1/2_ values were then averaged for each antibody group to report the mean plasma half-life.

#### Cryo-electron microscopy (cryo-EM)

0.1 mg of either N49P7-FR or eN49P7-FRv1-23 Fab was mixed with 0.1 mg of BG505 MD39 SOSIP.664 and 0.1 mg RM20A3 Fab (a Rhesus macaque mAb that recognizes the base of BG505 MD39 SOSIP.664 and increases angular sampling in cryo-EM). The complexes were incubated overnight at 22 °C. The following morning, the samples were concentrated using Amicon 100 kDa MWCO centrifugal filters to about 5.9 mg/mL. Detergent lauryl maltose neopentyl glycol (LMNG; Anatrace) was added to the sample to a final concentration of 0.005 mM shortly before freezing. Cryo grids were prepared using a Vitrobot Mark IV (Thermo Fisher Scientific) and the following settings: 4 °C temperature, 100% humidity, blot force 1, and wait time 10 s. Blotting time varied from 3 to 6 s. A 3.5 µL drop of either complex was applied to glow-discharged UltrAufoil 1.2/1.3-300 (Quantifoil Micro Tools GmbH) grids. Following blotting, the grids were plunge-frozen into liquid nitrogen-cooled liquid ethane. Sample grids were clipped, loaded into an Autoloader cassette, and screened using a Thermo Fisher Scientific Glacios TEM operating at 200 kV and equipped with a Thermo Fisher Scientific Falcon 4i direct electron detector. Exposure magnification was set to 190,000x with a pixel size at the specimen plane of 0.718 Å. EPU software (Thermo Fisher Scientific) was used for automated data collection. CryoSPARC Live^83^ was used for micrograph movie frame motion correction, dose weighting, and CTF correction. The remainder of data processing took place in cryoSPARC. Particle picking was initially performed using blob picker, followed by template picker. Multiple rounds of 2D classification were performed, followed by 3D *ab initio* reconstruction. The remaining particles were then re-extracted, and Fourier cropped by a factor of ∼1.4x (to reduce memory requirements of large box sizes), resulting in image pixel sizes of 1.034 Å. Non-uniform 3D refinement was performed with C3 symmetry enforced and with global CTF refinement of beamtilt and trefoil. Global resolution was estimated using half maps and a Fourier shell correlation cutoff of 0.143. Final data collection and processing statistics are summarized in Supplementary Figure 23 and Supplementary Table 10. Atomic modeling was performed in ChimeraX^84^ and Coot^85^, followed by real space refinement in Phenix^86^. Starting atomic coordinates for BG505 MD39 SOSIP were taken from PDB 6DFG, while PDB 6BCK was used for N49P7-FR and eN49P7-FRv1-23, with mutations performed in Coot to match the sequences of the samples. Final models were validated using MolProbity and EMRinger in the Phenix suite, and statistics are summarized in Supplementary Figure 23 and Supplementary Table 10. Figures were generated using UCSF ChimeraX.

#### Molecular dynamics (MD) simulations

We used the cryo-EM structures of N49P7-FR and eN49P7-FRv1-23 in complex with BG505 MD39 SOSIP as starting structures for MD simulations. Starting structures were prepared in Molecular Operating Environment (Chemical Computing Group, version 2024.06) using the Protonate3D tool^87^, followed by system setup with CHARMM-GUI^88,89^. Each structure model was solvated in a cubic TIP3P water box with a minimum wall distance to the protein of 12 Å, and the charge was neutralized with K^+^/Cl^−^ ions up to a concentration of 0.15 mM^90^. Simulations employed the AMBER force field 19SB^91^ and the GLYCAM-06j force field for glycans^92^. For each antibody–gp120 complex, three independent 1-µs classical MD simulations were performed with Amber24^93^ in an NpT ensemble using pmemd.cuda^94^. Bonds involving hydrogen atoms were constrained with SHAKE^95^, allowing a 2 fs timestep. Temperature (300 K) was maintained with a Langevin thermostat (collision frequency 2 ps^−1^)^96,97^, and pressure with a Monte Carlo barostat (1 volume change attempt per 100 steps)^98^. Electrostatic and van der Waals interaction energies were calculated with CPPTRAJ (linear interaction energy method)^99^, averaged across all simulation frames. Binding interface interactions and their frequencies were analyzed using GetContacts (https://getcontacts.github.io/). Cluster analysis was performed in CPPTRAJ, and structures were visualized in ChimeraX^84^.

## Figure descriptions

**Supplementary Figure 1.**
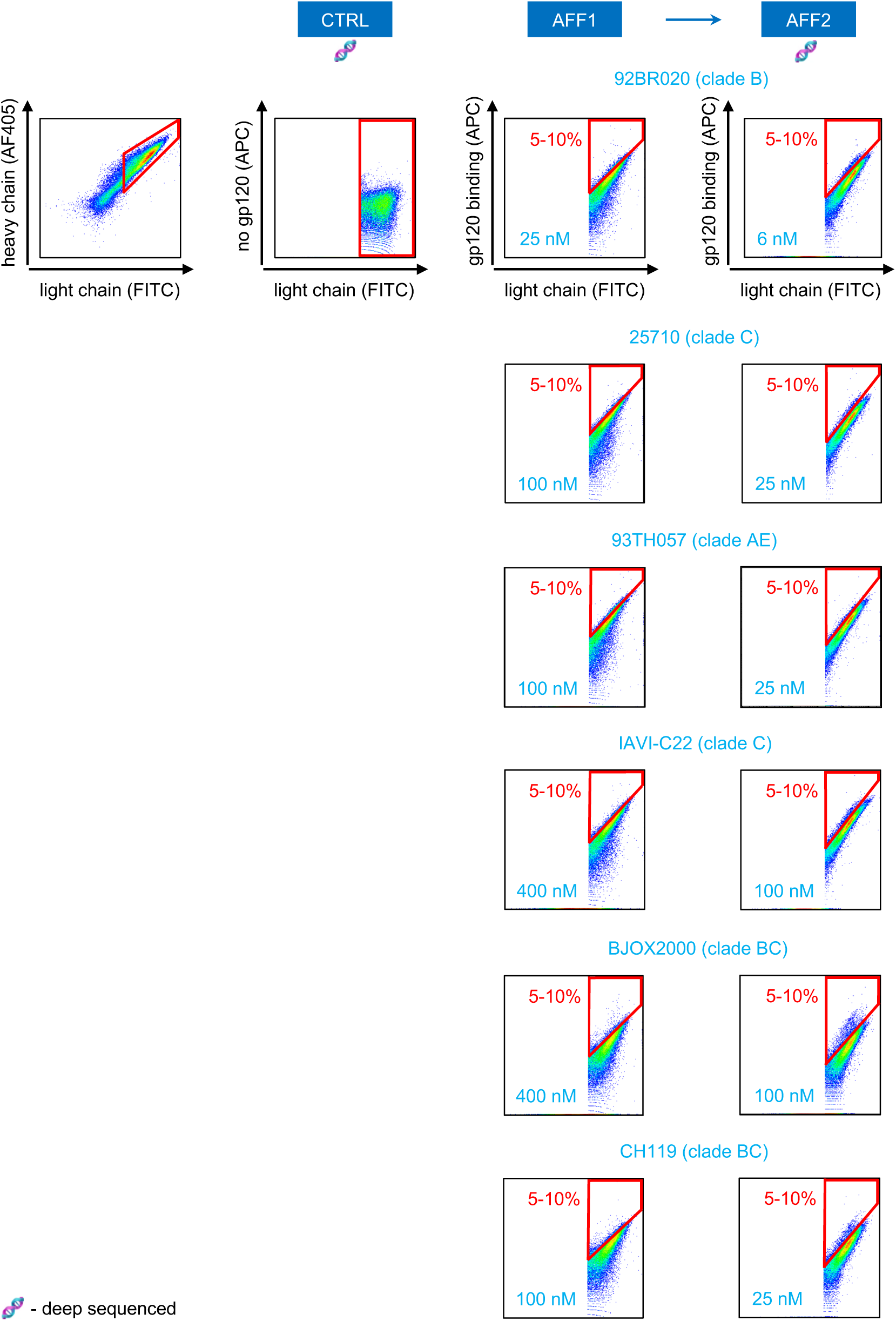
FACS screening strategy to enrich N49P7-FR VH dbSMS library variants with the highest affinity to six HIV gp120s. All VH variants were paired with the wild-type VL. FACS plots with sorting gates (red) and the percentage of the displayed library population selected for sorting are shown for each gp120 used at the indicated concentration. CTRL – control (unselected) population of displayed library variants not incubated with gp120 (deep sequenced for normalization to calculate mutation enrichment); AFF1 – affinity screening round 1; AFF2 – affinity screening round 2 (a sorted population was deep sequenced to calculate mutation enrichment). The experiment was performed in duplicate.

**Supplementary Figure 2.**
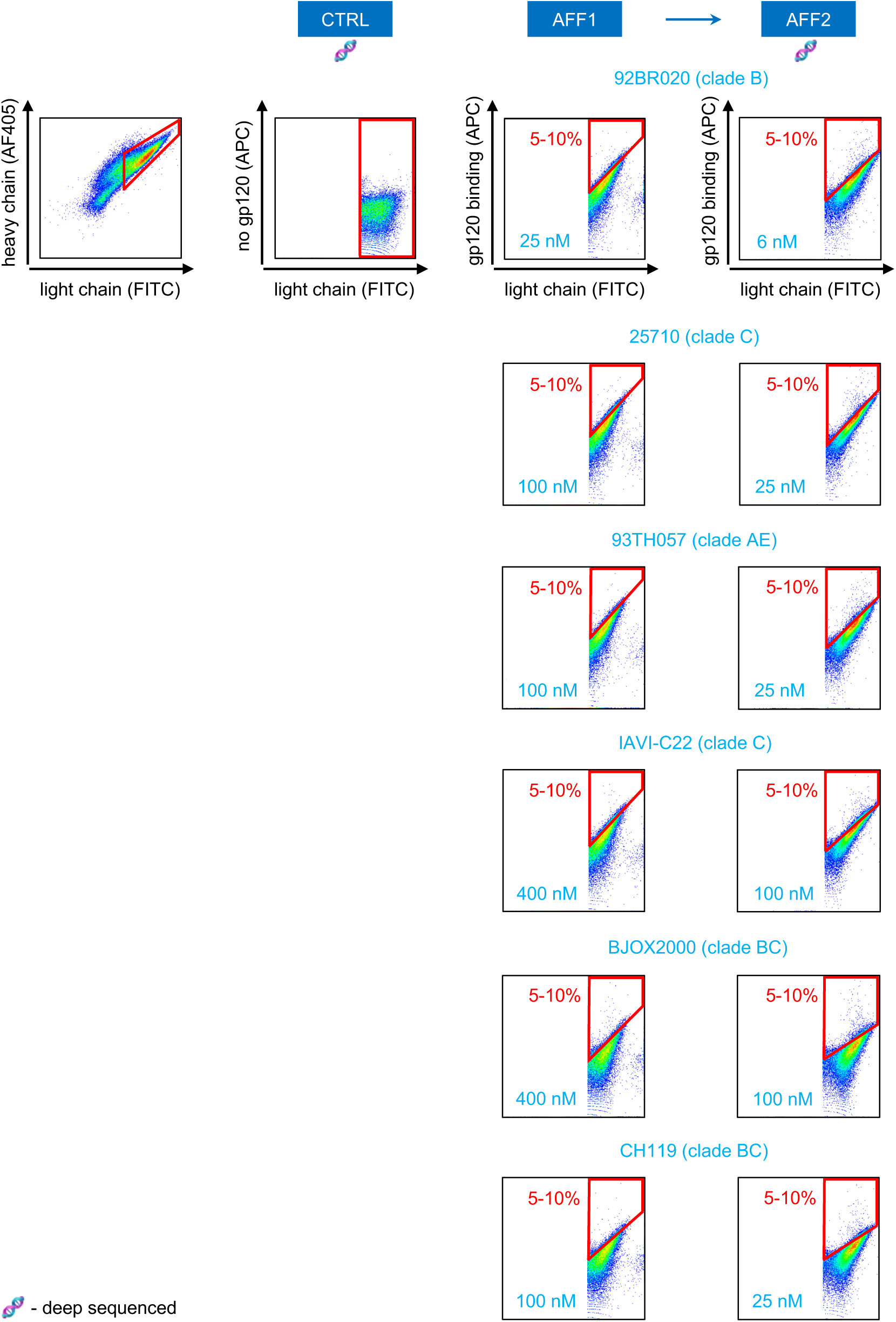
FACS screening strategy to enrich N49P7-FR VL dbSMS library variants with the highest affinity to six HIV gp120s. All VL variants were paired with the wild-type VH. FACS plots with sorting gates (red) and the percentage of the displayed library population selected for sorting are shown for each gp120 used at the indicated concentration. CTRL – control (unselected) population of displayed library variants not incubated with gp120 (deep sequenced for normalization to calculate mutation enrichment); AFF1 – affinity screening round 1; AFF2 – affinity screening round 2 (a sorted population was deep sequenced to calculate mutation enrichment). The experiment was performed in duplicate.

**Supplementary Figure 3.**
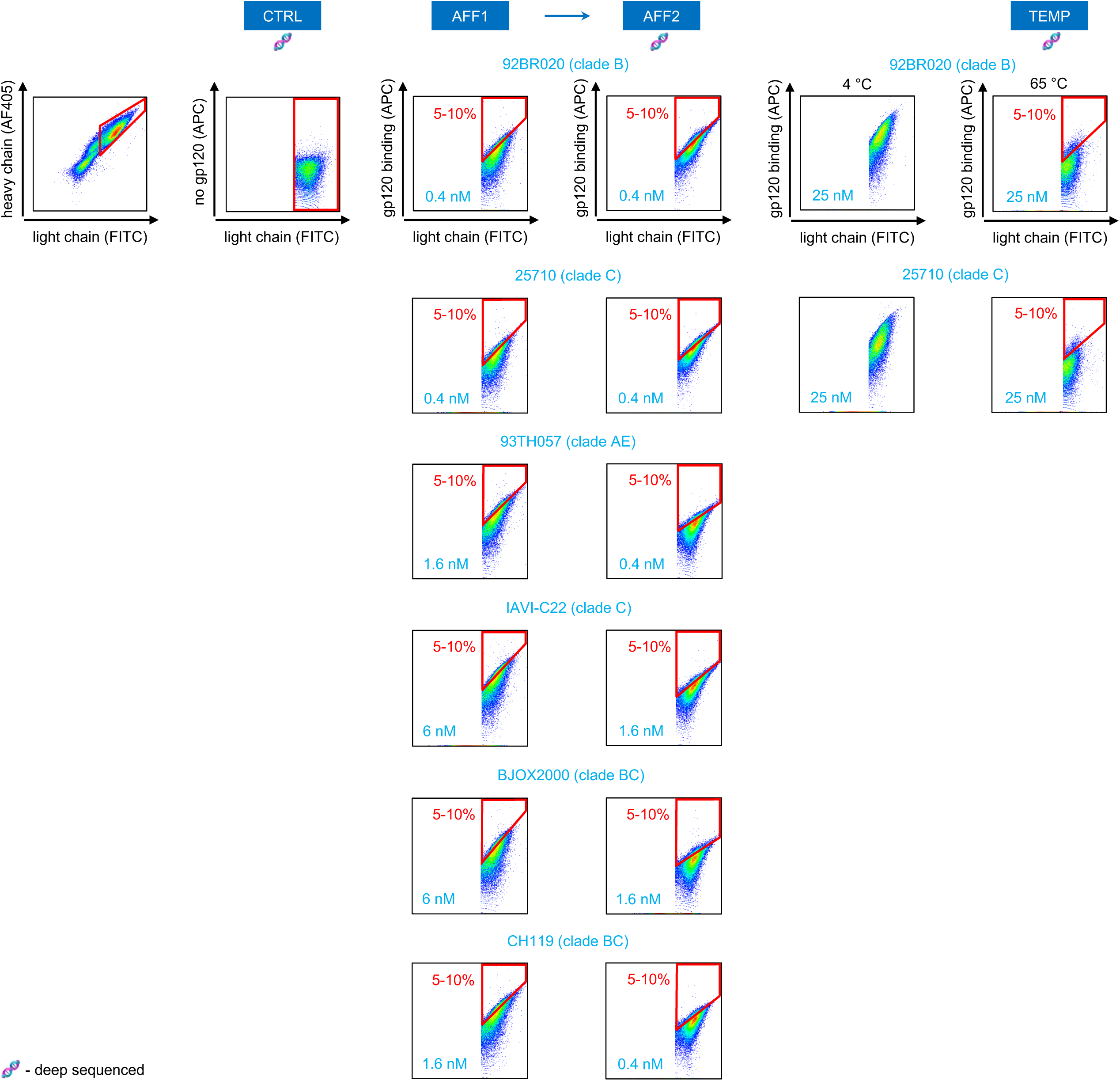
FACS screening strategy to enrich N49P7-FR VHv1 dbSMS library variants with the highest affinity to six HIV gp120s and enhanced thermal stability. All VHv1 variants were paired with the optimized VL (VLv3). FACS plots with sorting gates (red) and the percentage of the displayed library population selected for sorting are shown for each gp120 used at the indicated concentration. CTRL – control (unselected) population of displayed library variants not incubated with gp120 (deep sequenced for normalization to calculate mutation enrichment); AFF1 – affinity screening round 1; AFF2 – affinity screening round 2 (a sorted population was deep sequenced to calculate mutation enrichment); TEMP – affinity screening following library incubation at 65 °C (a sorted population was deep sequenced to calculate mutation enrichment). The experiment was performed in duplicate.

**Supplementary Figure 4.**
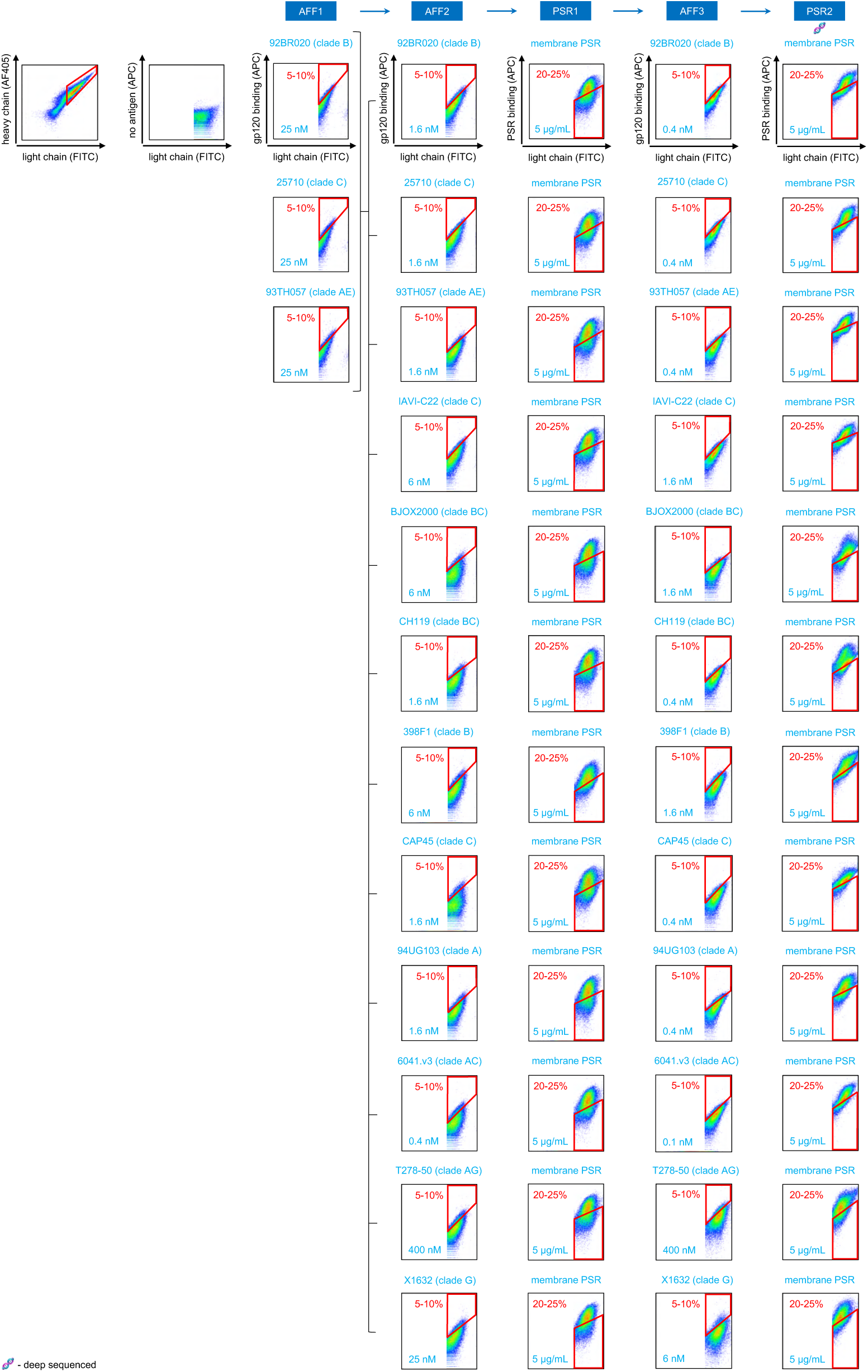
FACS screening strategy to enrich N49P7-FR VH combinatorial library variants with the highest affinity to twelve HIV gp120s and the lowest polyreactivity with PSR. All VH variants were paired with the wild-type VL. FACS plots with sorting gates (red) and the percentage of the displayed library population selected for sorting are shown for each gp120 and PSR used at the indicated concentration. AFF1-3 – affinity screening rounds 1-3; PSR1-2 – polyreactivity screening rounds 1-2. After AFF1, all sorted populations were combined, grown, induced, and split to perform subsequent sorts with twelve gp120s in parallel. After the last sort (PSR2), populations were deep sequenced to calculate variant frequency.

**Supplementary Figure 5.**
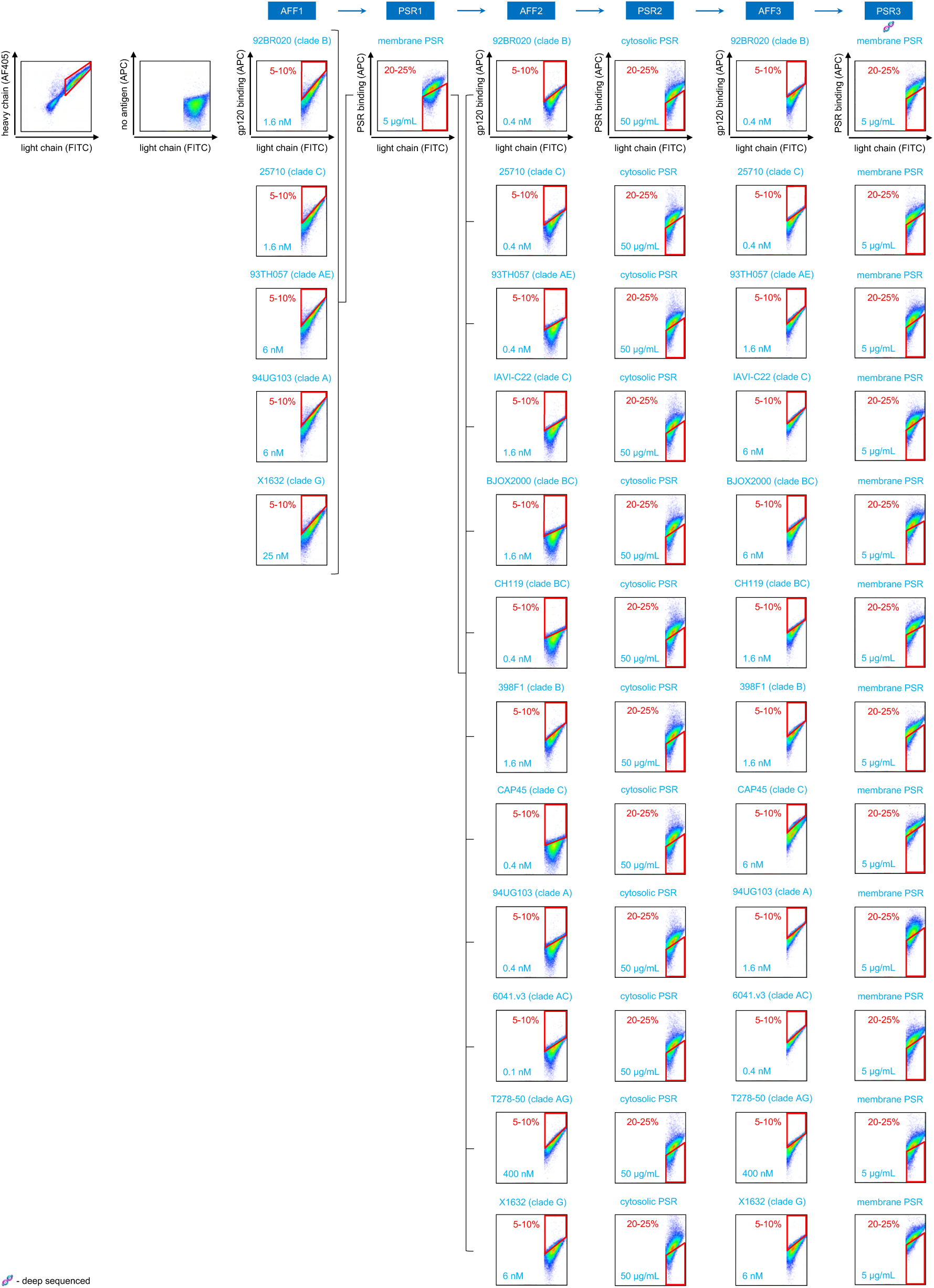
FACS screening strategy to enrich N49P7-FR VL combinatorial library variants with the highest affinity to twelve HIV gp120s and the lowest polyreactivity with PSR. All VL variants were paired with one of the optimized VH variants (VHv10). FACS plots with sorting gates (red) and the percentage of the displayed library population selected for sorting are shown for each gp120 and PSR used at the indicated concentration. AFF1-3 – affinity screening rounds 1-3; PSR1-3 – polyreactivity screening rounds 1-3. After AFF1, all sorted populations were combined, grown, induced, screened for polyreactivity (PSR1), grown, induced, and split to perform subsequent sorts with twelve gp120s in parallel. After the last sort (PSR3), populations were deep sequenced to calculate variant frequency.

**Supplementary Figure 6.**
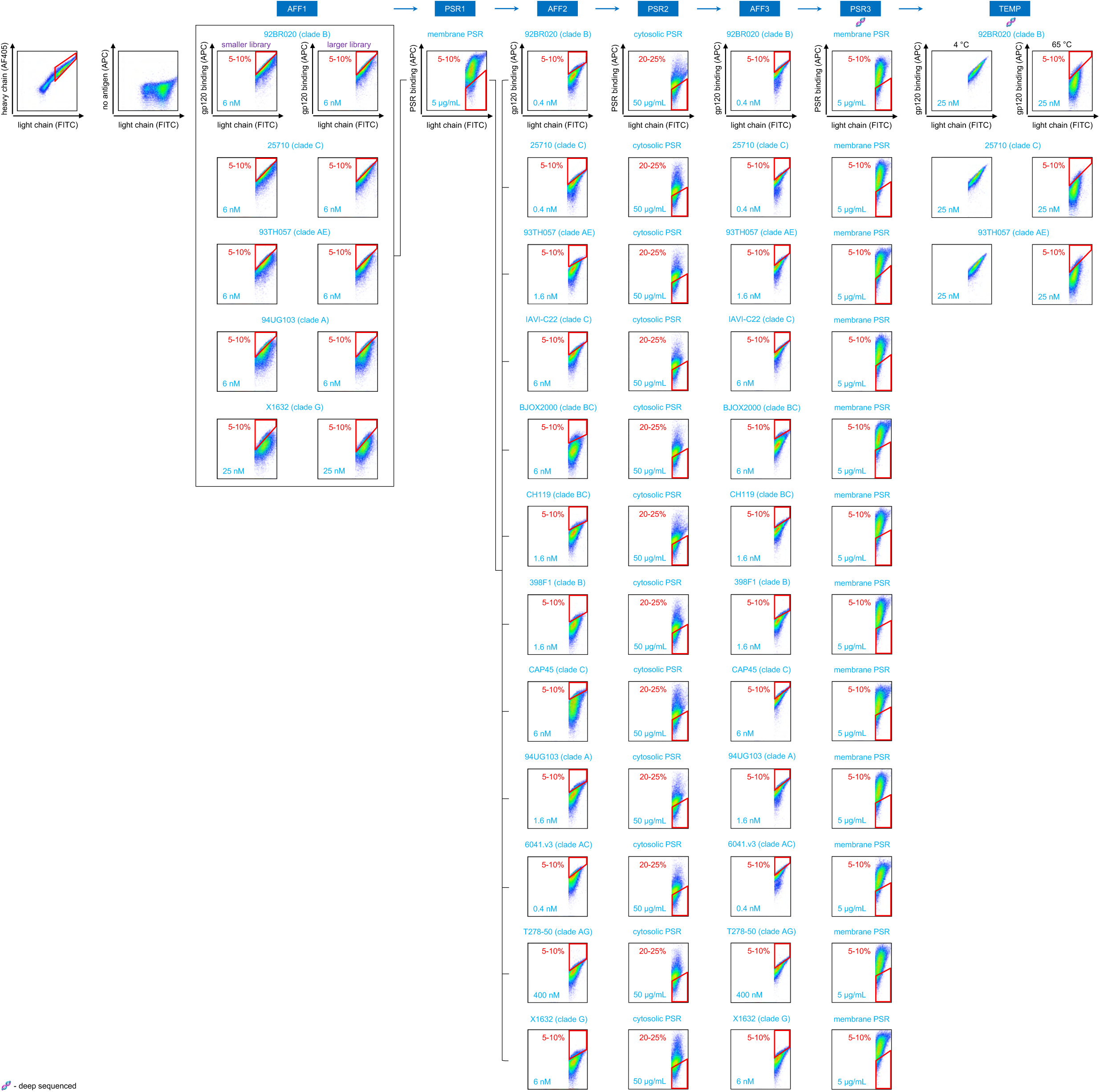
FACS screening strategy to enrich N49P7-FR VHv1 combinatorial library variants with the highest affinity to twelve HIV gp120s and the lowest polyreactivity with PSR. All VHv1 variants were paired with the optimized VL (VLv3). FACS plots with sorting gates (red) and the percentage of the displayed library population selected for sorting are shown for each gp120 and PSR used at the indicated concentration. AFF1-3 – affinity screening rounds 1-3; PSR1-3 – polyreactivity screening rounds 1-3. TEMP – affinity screening following library incubation at 65°C. After AFF1, all sorted populations from a smaller library (sampling 21 positions) and a larger library (sampling 28 positions) were combined, grown, induced, screened for polyreactivity (PSR1), grown, induced, and split to perform subsequent sorts with twelve gp120s in parallel. After PSR3 and TEMP, sorted populations were deep sequenced to calculate variant frequency.

**Supplementary Figure 7.**
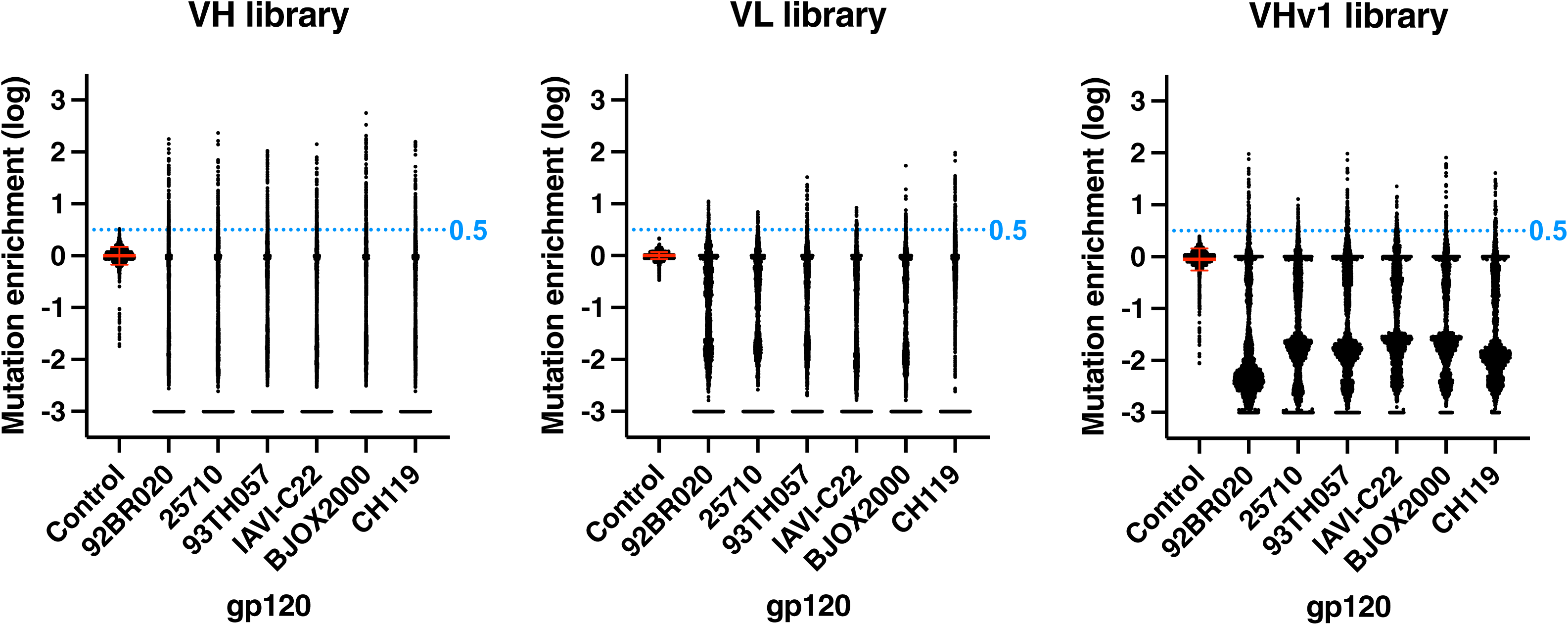
Distribution of mutation enrichment values across all N49P7-FR VH/VL positions in gp120-selected and unselected (control) dbSMS libraries. Each datapoint (dot) represents the enrichment of a single amino acid substitution at a specific VH or VL position, normalized to the reference control and averaged across two replicates. Horizontal and vertical red bars indicate the mean and standard deviation (SD), respectively, of mutation enrichment in the control. A threshold for identifying significantly enriched mutations in the gp120-selected populations (≥0.5; horizontal blue dotted line) was set to distinguish gp120-driven enrichment from background variability.

**Supplementary Figure 8.**
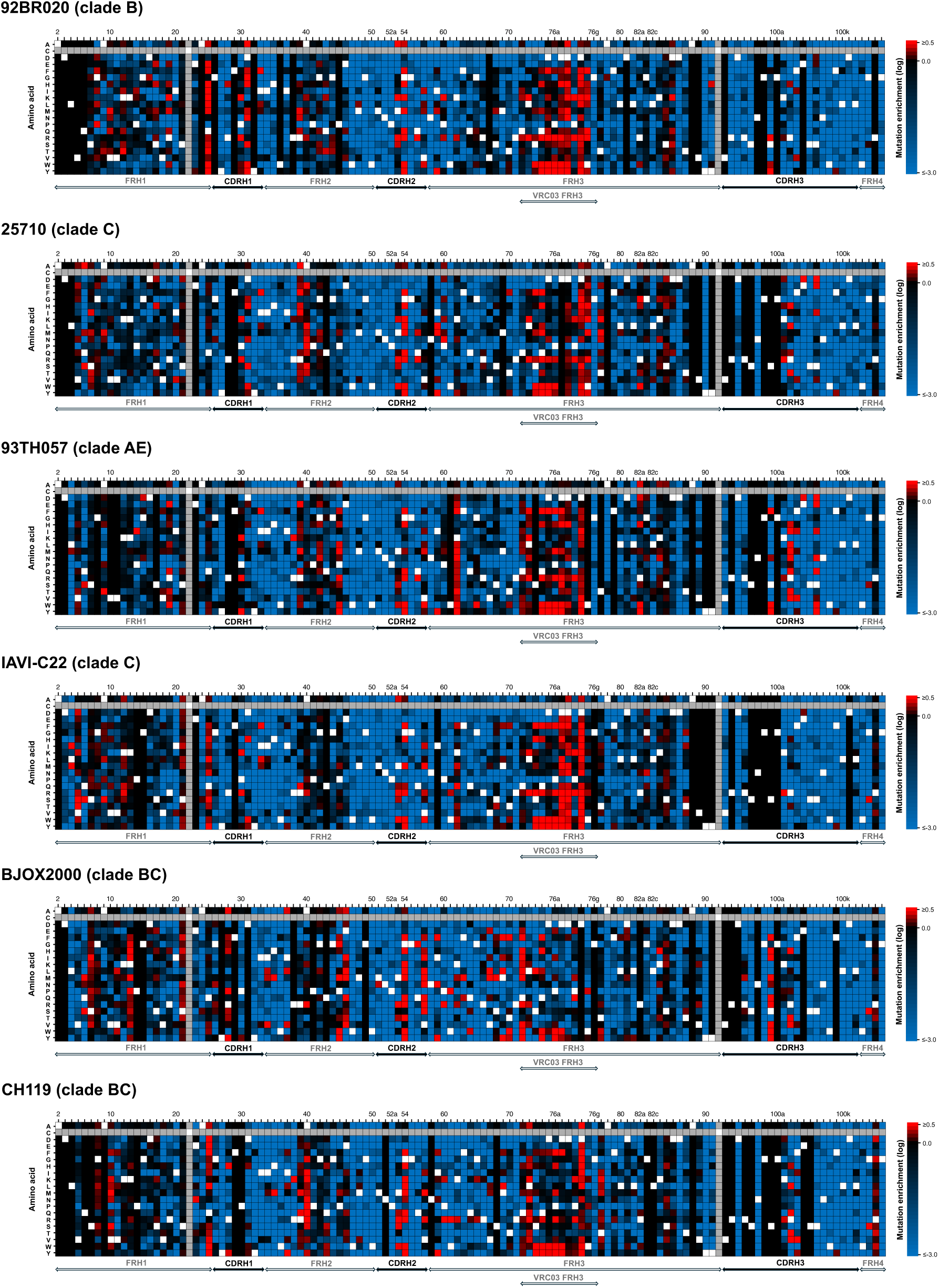
Heat maps showing amino acid enrichment across all VH positions in the dbSMS library selected with six gp120s in parallel. Amino acids enhancing the gp120 interaction (red), diminishing it (blue), having a neutral effect (black), wild-type (white), and not sampled in the library (gray) are indicated. Amino acid positions are numbered according to the Kabat scheme.

**Supplementary Figure 9.**
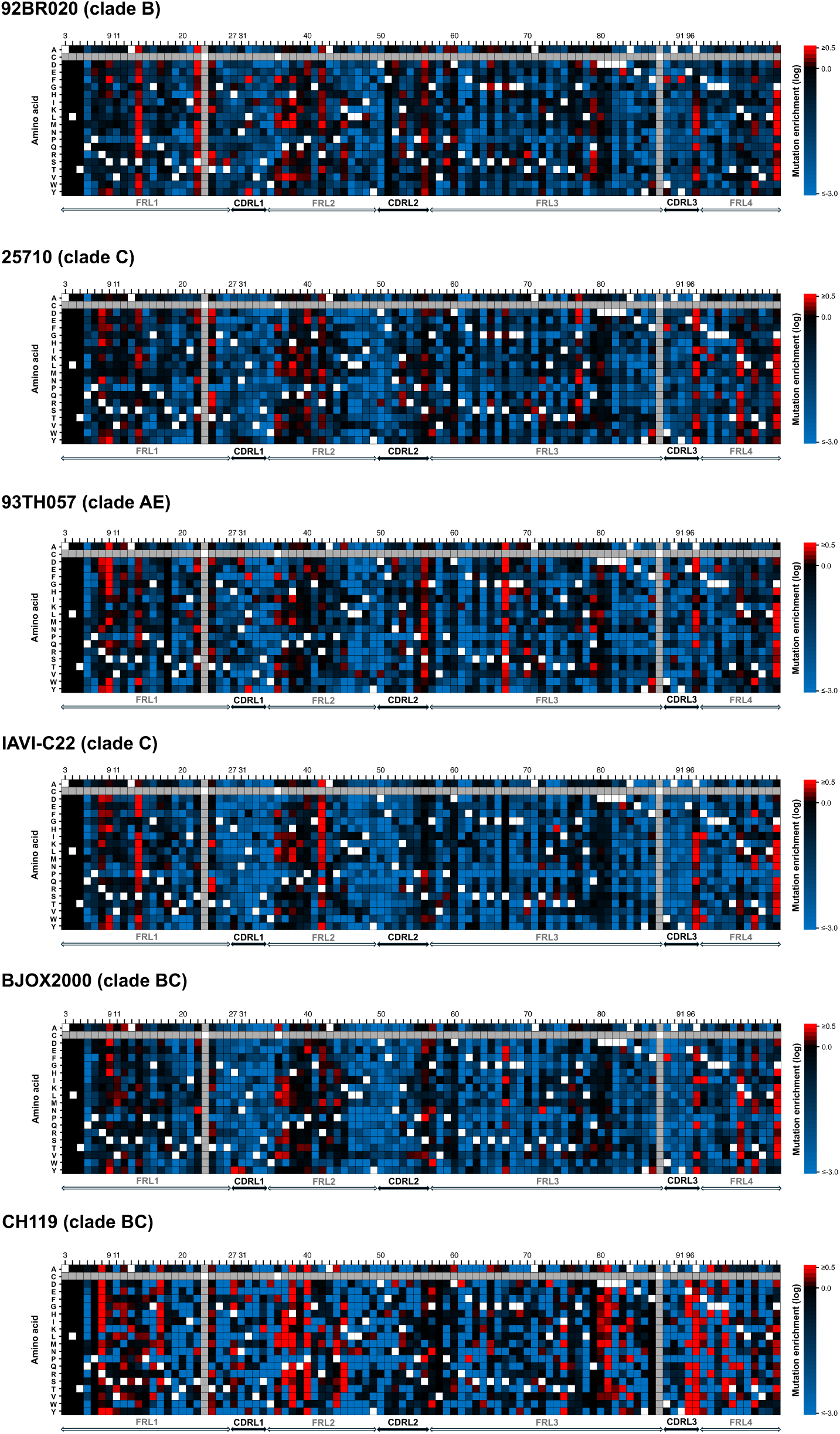
Heat maps showing amino acid enrichment across all VL positions in the dbSMS library selected with six gp120s in parallel. Amino acids enhancing the gp120 interaction (red), diminishing it (blue), having a neutral effect (black), wild-type (white), and not sampled in the library (gray) are indicated. Amino acid positions are numbered according to the Kabat scheme.

**Supplementary Figure 10.**
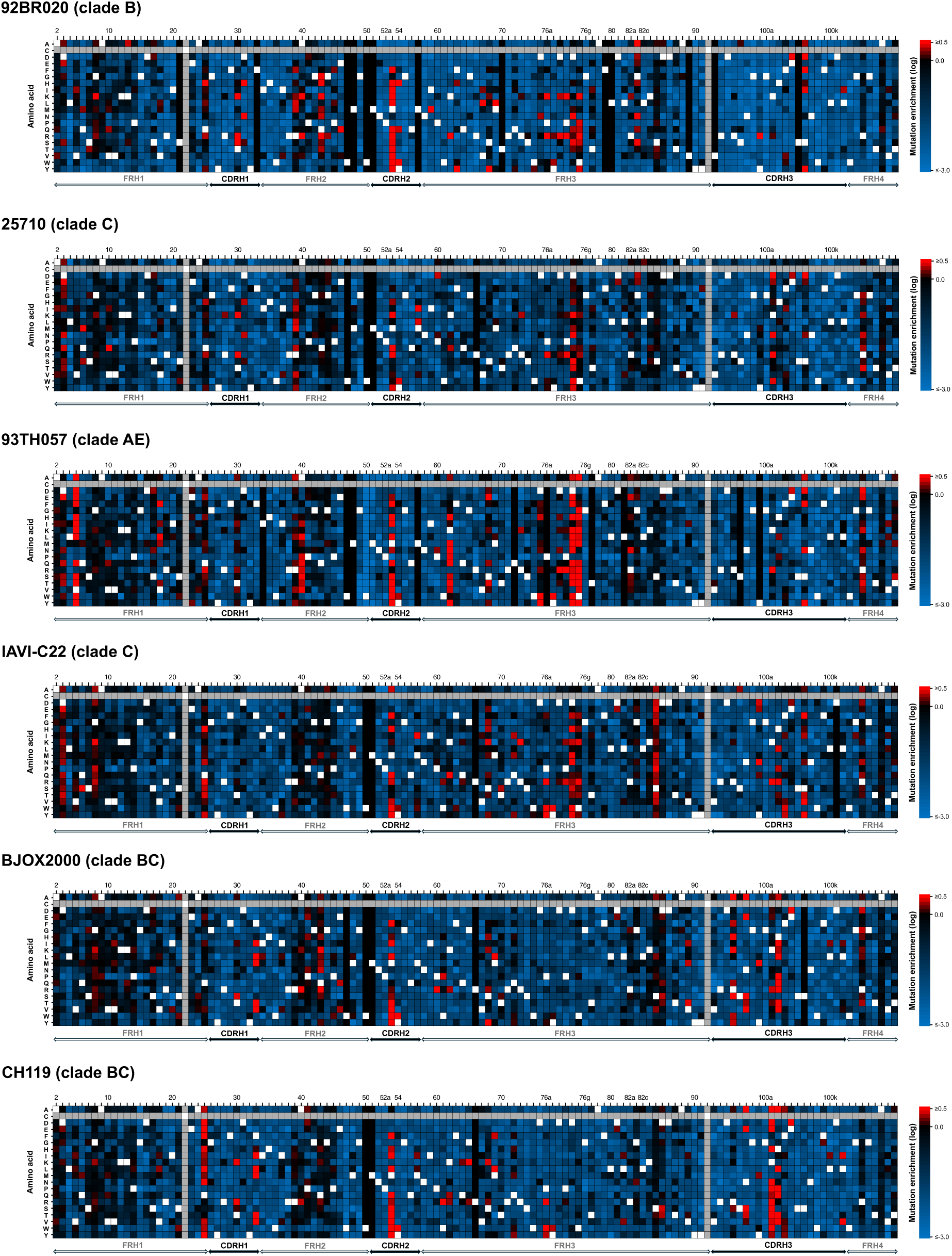
Heat maps showing amino acid enrichment across all VHv1 positions in the dbSMS library selected with six gp120s in parallel. Amino acids enhancing the gp120 interaction (red), diminishing it (blue), having a neutral effect (black), wild-type (white), and not sampled in the library (gray) are indicated. Amino acid positions are numbered according to the Kabat scheme.

**Supplementary Figure 11.**
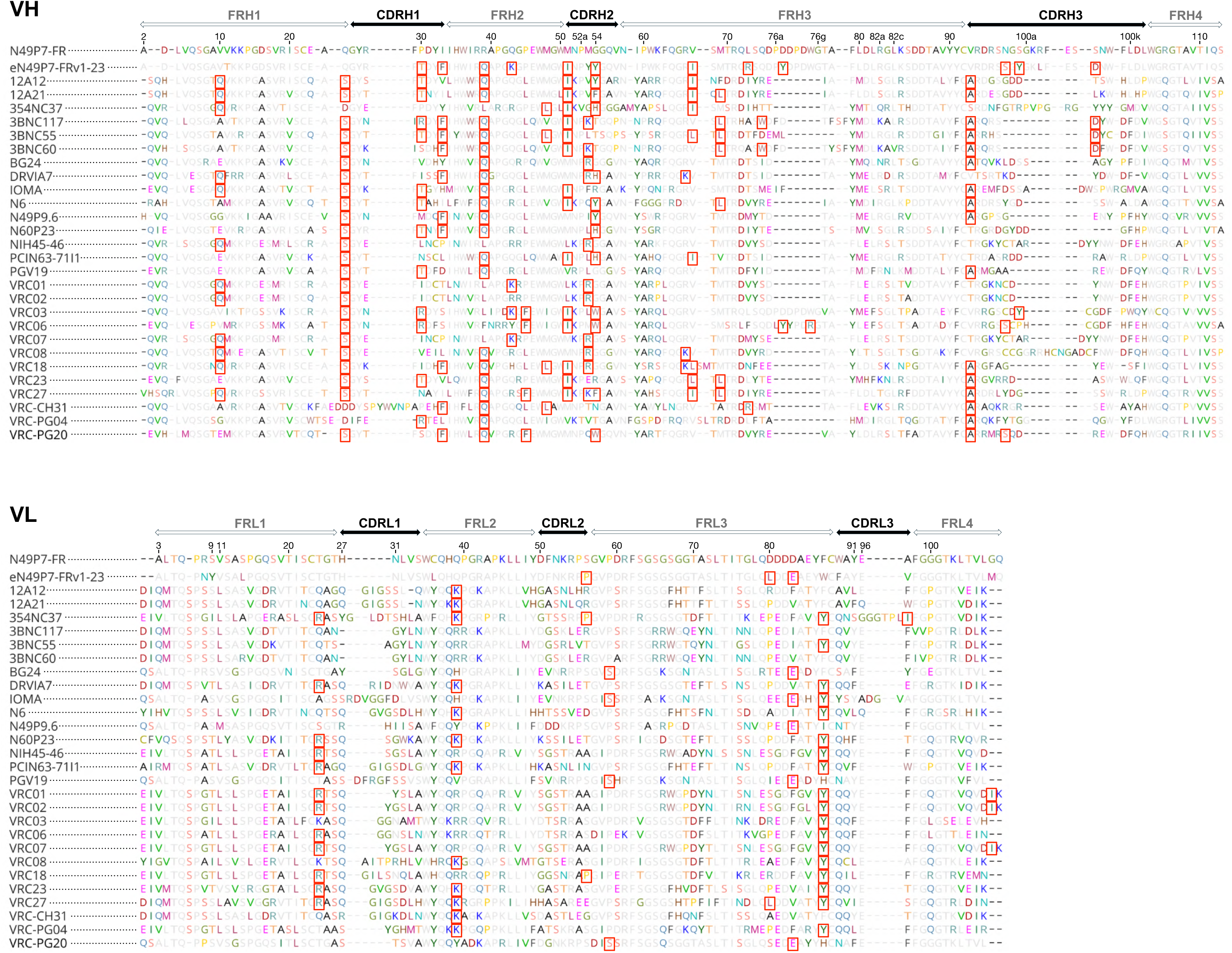
Alignment of VH and VL amino acid sequences of N49P7-FR, eN49P7-FRv1-23, and naturally occurring VRC01-class bnAbs. Amino acids highlighted with red frames represent residues that were found in at least one of the listed bnAbs and were sampled in our combinatorial libraries. The VH and VL chains of eN49P7-FRv1-23 contain 12 and 3 mutations, respectively, that are present in other VRC01-class bnAbs. Amino acid positions are numbered according to the Kabat scheme.

**Supplementary Figure 12.**
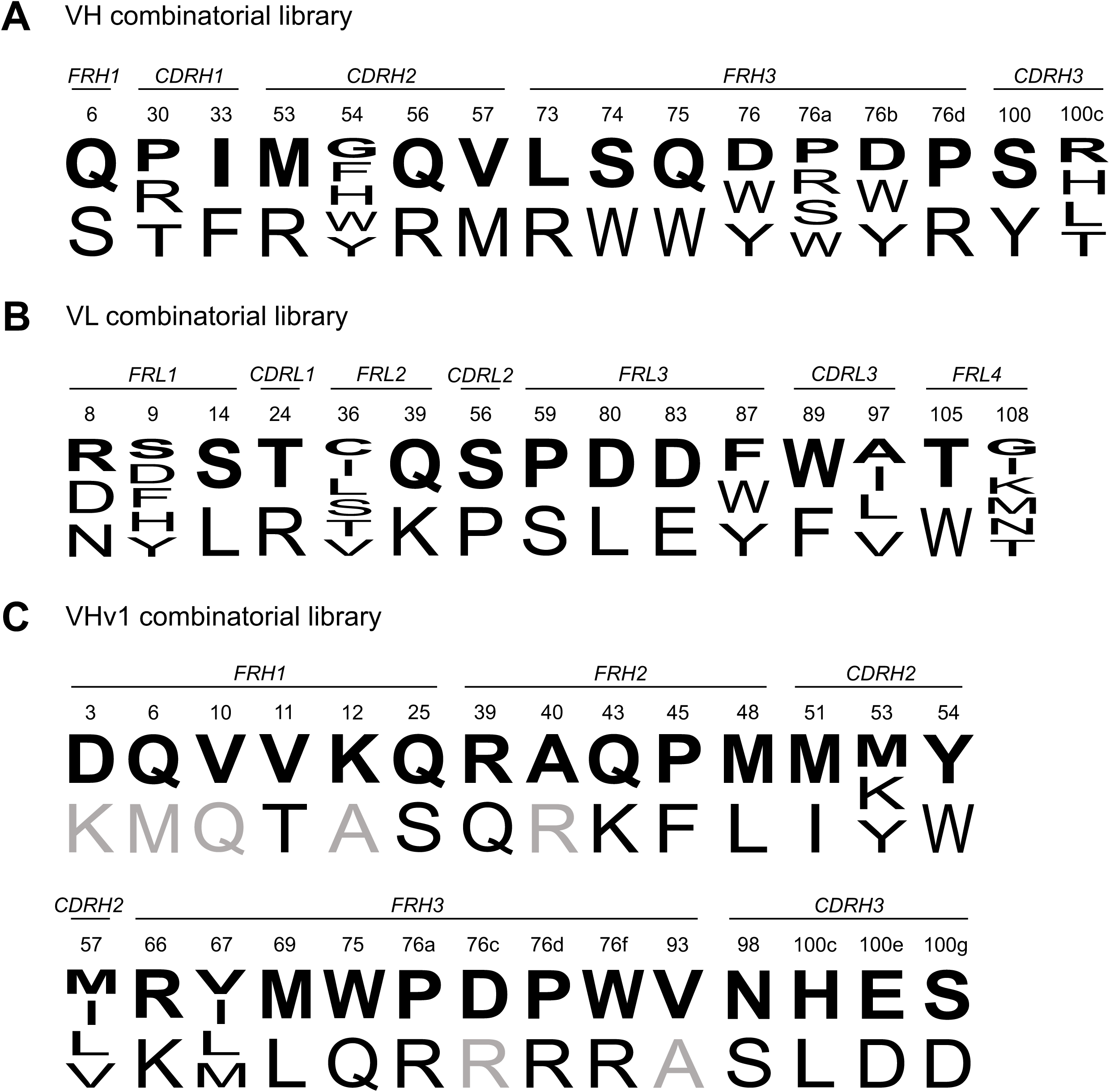
Amino acids sampled in the N49P7-FR combinatorial libraries. A total of 16 positions in VH (**A**), 15 positions in VL (**B**), and 28 positions in VHv1 (**C**) were sampled. For each position (numbered according to the Kabat scheme), the first amino acid (in bold) represents the wild-type residue, followed by 1–5 substitutions. Substitutions shown in gray in panel C indicate amino acids sampled only in a larger library, in addition to the substitutions included in a smaller library (Supplementary Fig. 6).

**Supplementary Figure 13.**
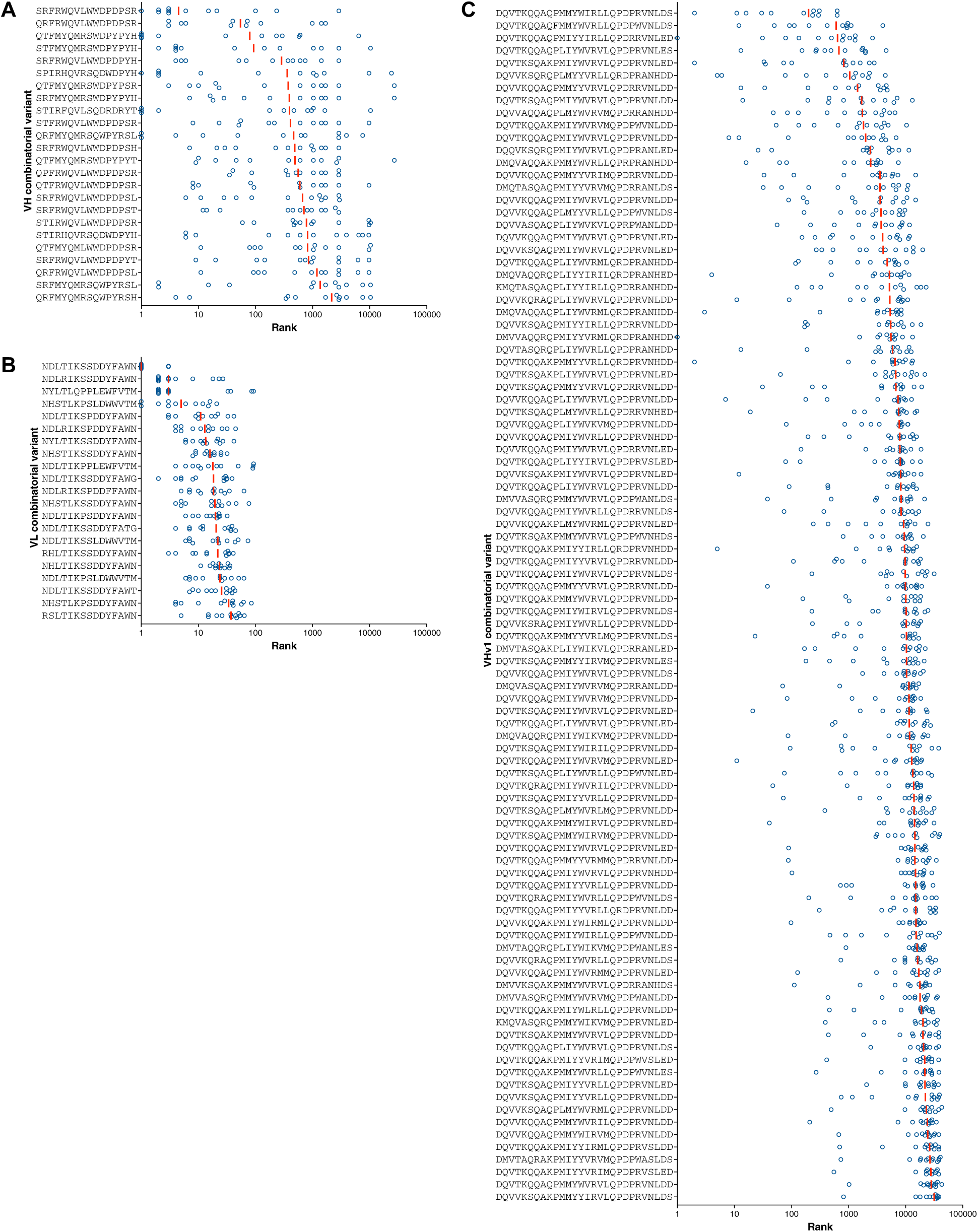
Ranking of the most enriched variants identified in the N49P7-FR combinatorial libraries screened against twelve gp120s. 24 VH variants (**A**), 21 VL variants (**B**), and 96 VHv1 variants (**C**) were reformatted as IgG1 and evaluated for functionality and developability to select lead candidates. For each variant, only the sampled amino acids are shown, ordered by their position. Each dark blue circle represents the frequency rank of a variant after screening with one gp120. Vertical red bars indicate median frequency across the different sorts.

**Supplementary Figure 14.**
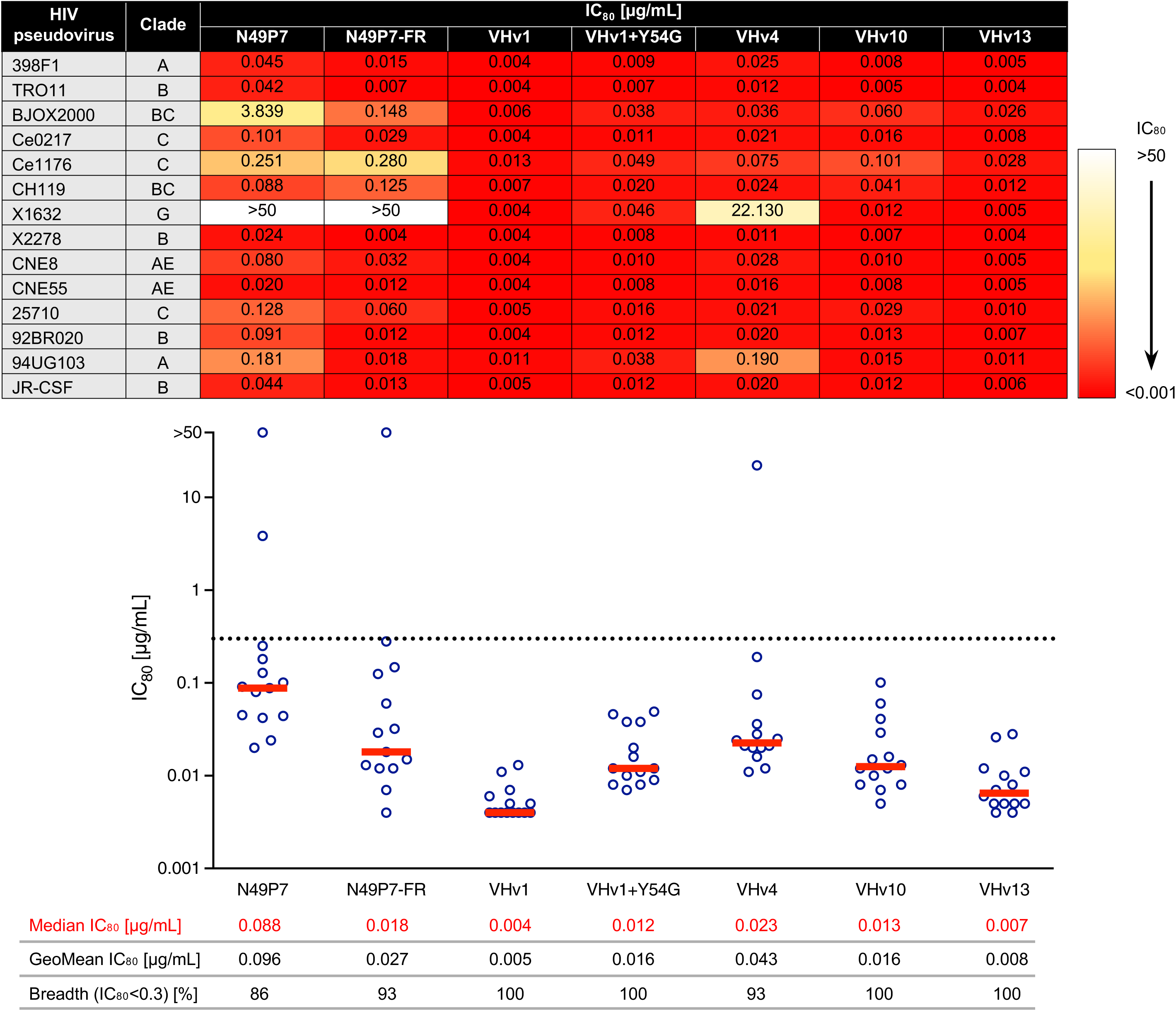
*In vitro* neutralization potency and breadth of N49P7-FR variants against selected HIV cross-clade pseudoviruses from the 12-virus and 6-virus panels. Each circle in the graph represents an IC_80_ value for one pseudoviral strain. Horizontal red bars indicate the median IC_80_ values, and the horizontal dotted line marks IC_80_=0.3 μg/mL. bnAbs were tested against each pseudovirus in duplicate.

**Supplementary Figure 15.**
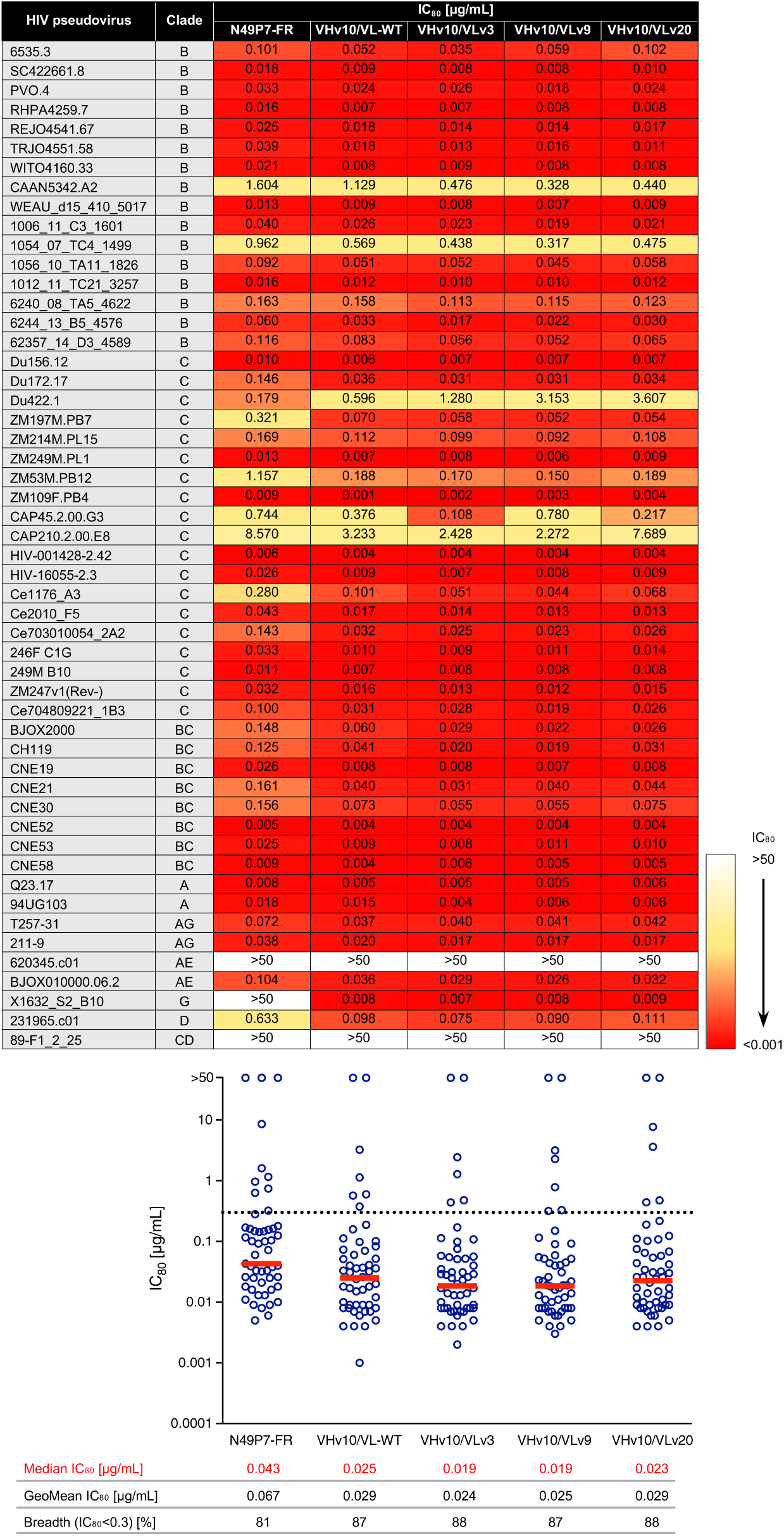
*In vitro* neutralization potency and breadth of N49P7-FR variants against selected HIV cross-clade pseudoviruses from the Seaman global panel. Each circle in the graph represents an IC_80_ value for one pseudoviral strain. Horizontal red bars indicate the median IC_80_ values, and the horizontal dotted line marks IC_80_=0.3 μg/mL. bnAbs were tested against each pseudovirus in duplicate.

**Supplementary Figure 16.**
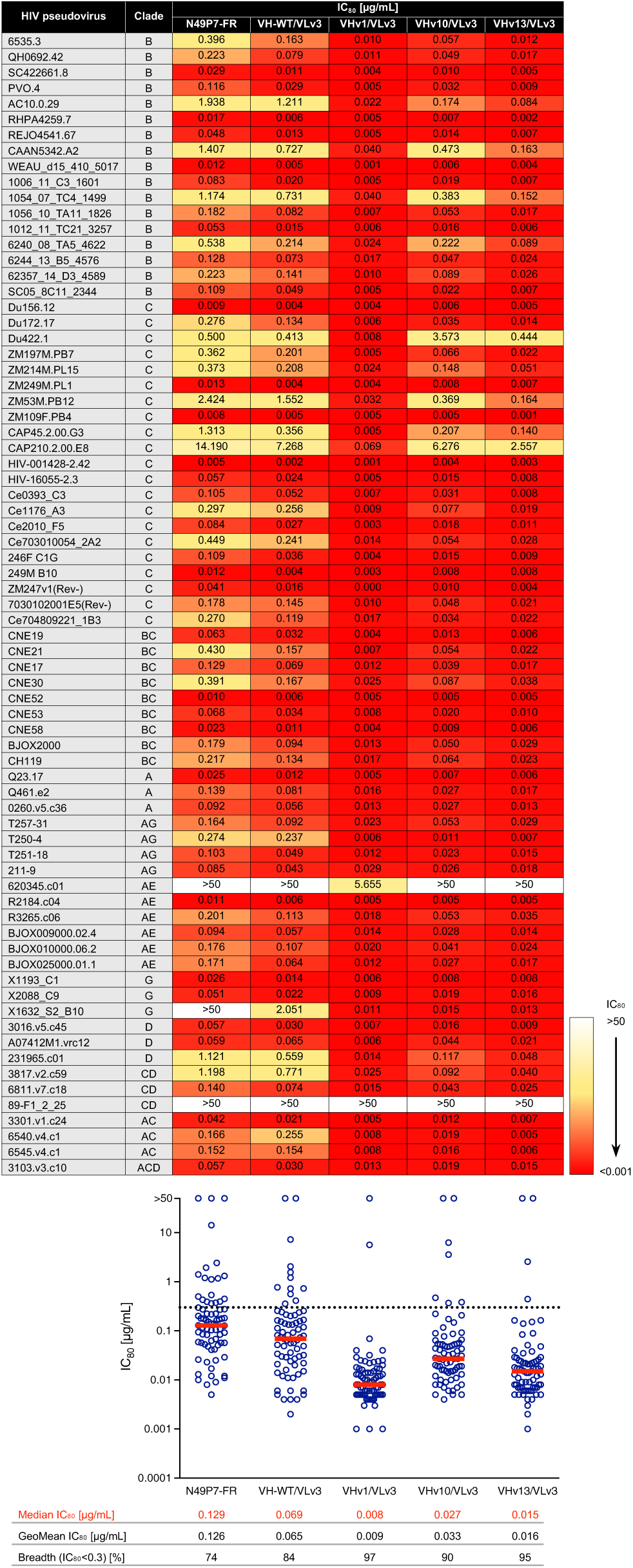
*In vitro* neutralization potency and breadth of N49P7-FR variants against selected HIV cross-clade pseudoviruses from the Seaman global panel. Each circle in the graph represents an IC_80_ value for one pseudoviral strain. Horizontal red bars indicate the median IC_80_ values, and the horizontal dotted line marks IC_80_=0.3 μg/mL. bnAbs were tested against each pseudovirus in duplicate.

**Supplementary Figure 17.**
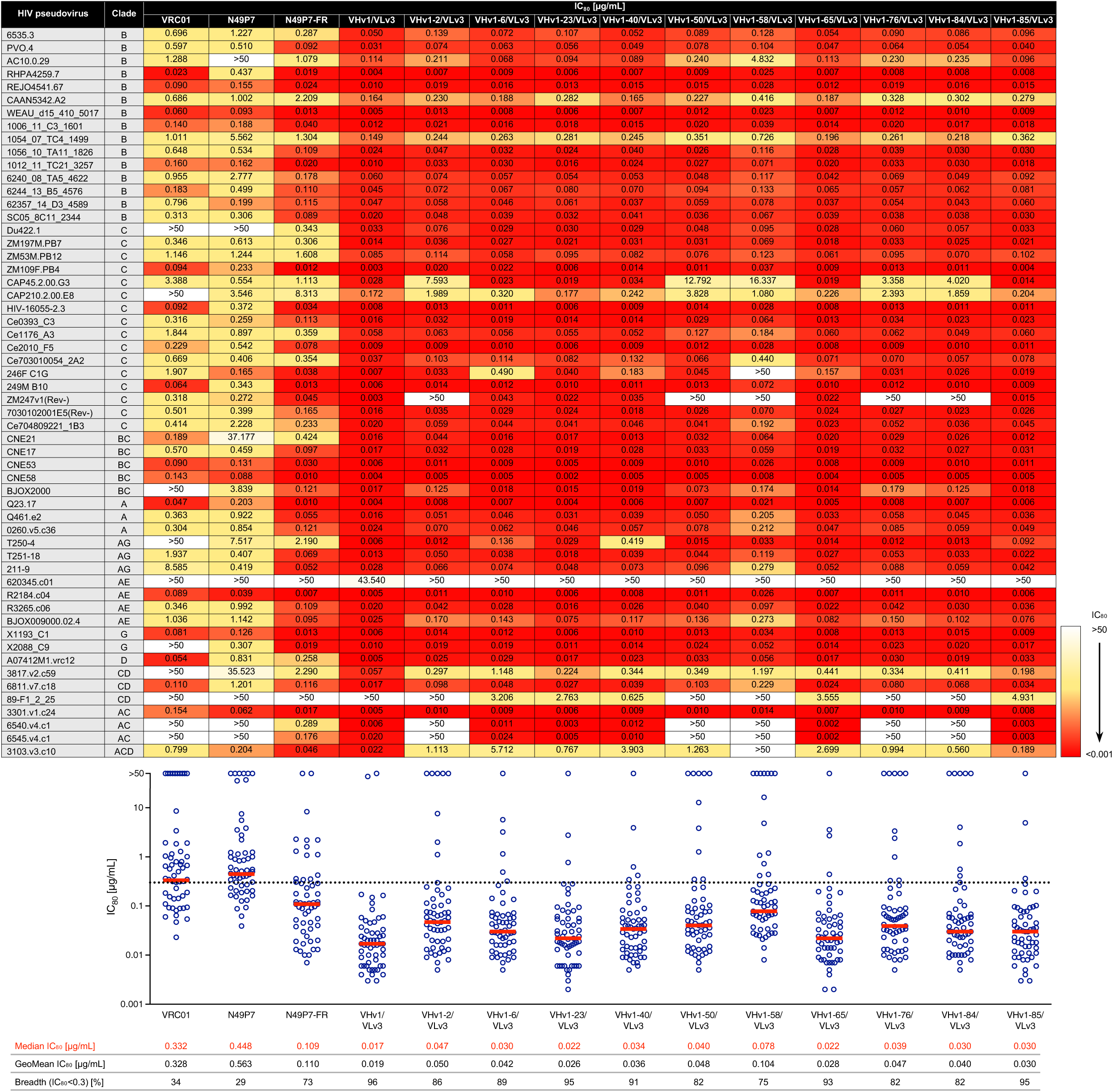
*In vitro* neutralization potency and breadth of VRC01 and N49P7 variants against selected HIV cross-clade pseudoviruses from the Seaman global panel. Each circle in the graph represents an IC_80_ value for one pseudoviral strain. Horizontal red bars indicate the median IC_80_ values, and the horizontal dotted line marks IC_80_=0.3 μg/mL. bnAbs were tested against each pseudovirus in duplicate.

**Supplementary Figure 18.**
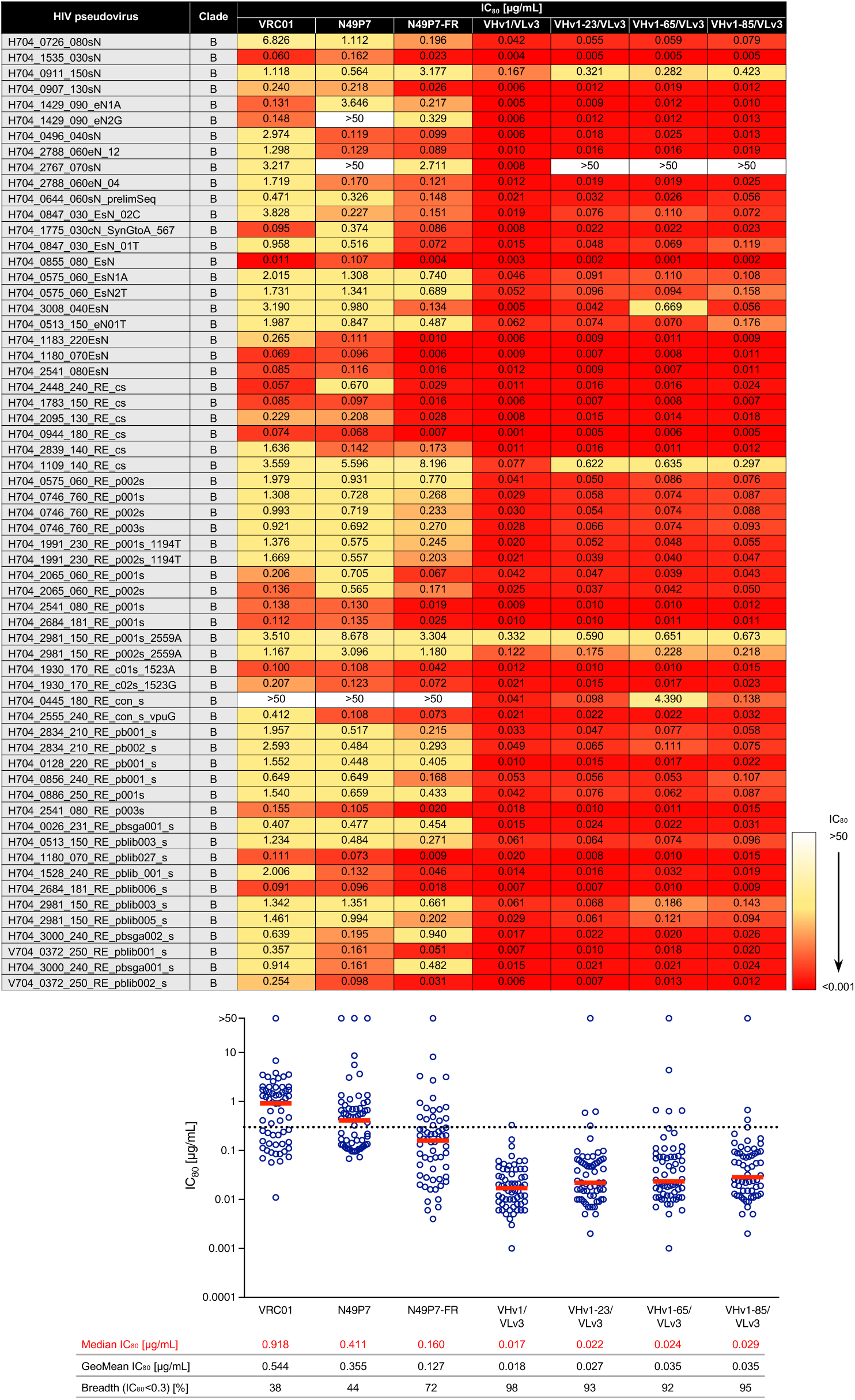
*In vitro* neutralization potency and breadth of VRC01 and N49P7 variants against HIV clade B pseudoviruses derived from the AMP trial placebo group viruses. Each circle in the graph represents an IC_80_ value for one pseudoviral strain. Horizontal red bars indicate the median IC_80_ values, and the horizontal dotted line marks IC_80_=0.3 μg/mL. bnAbs were tested against each pseudovirus in duplicate.

**Supplementary Figure 19.**
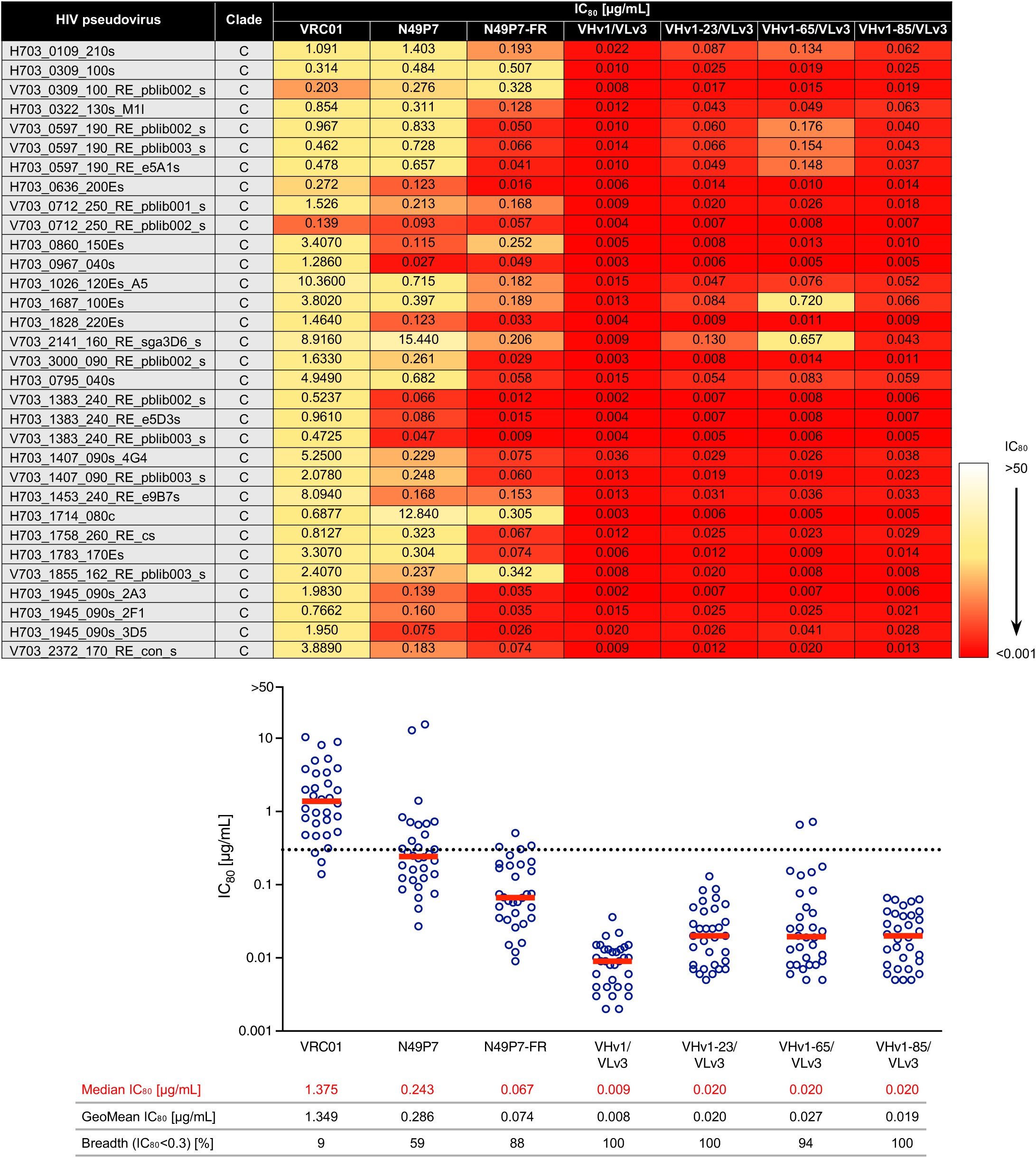
*In vitro* neutralization potency and breadth of VRC01 and N49P7 variants against HIV clade C pseudoviruses derived from the AMP trial breakthrough viruses. Each circle in the graph represents an IC_80_ value for one pseudoviral strain. Horizontal red bars indicate the median IC_80_ values, and the horizontal dotted line marks IC_80_=0.3 μg/mL. bnAbs were tested against each pseudovirus in duplicate.

**Supplementary Figure 20.**
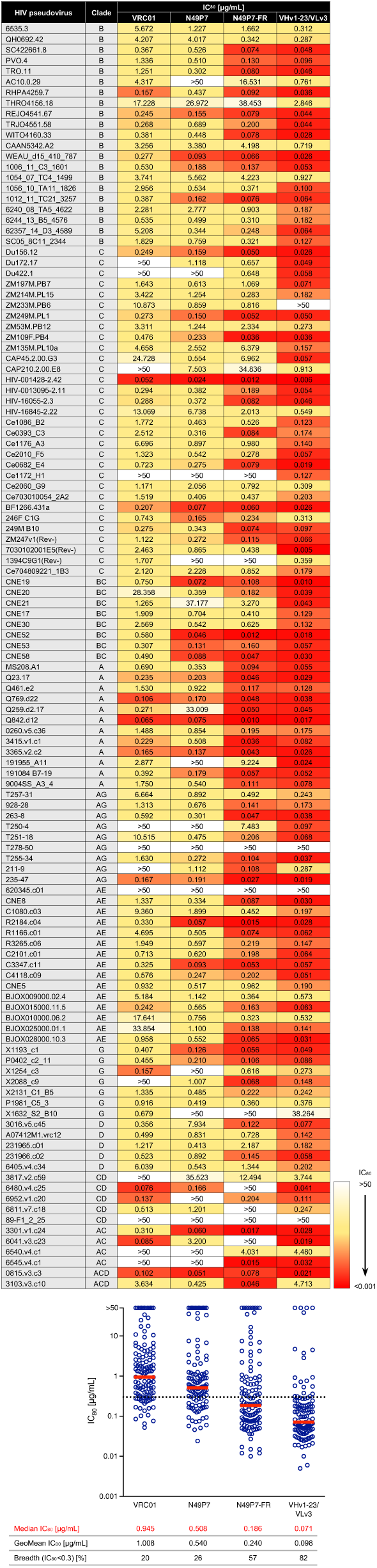
*In vitro* neutralization potency and breadth of VRC01 and N49P7 variants against HIV cross-clade pseudoviruses from the Seaman global panel. Each circle in the graph represents an IC_80_ value for one pseudoviral strain. Horizontal red bars indicate the median IC_80_ values, and the horizontal dotted line marks IC_80_=0.3 μg/mL. bnAbs were tested against each pseudovirus in duplicate.

**Supplementary Figure 21.**
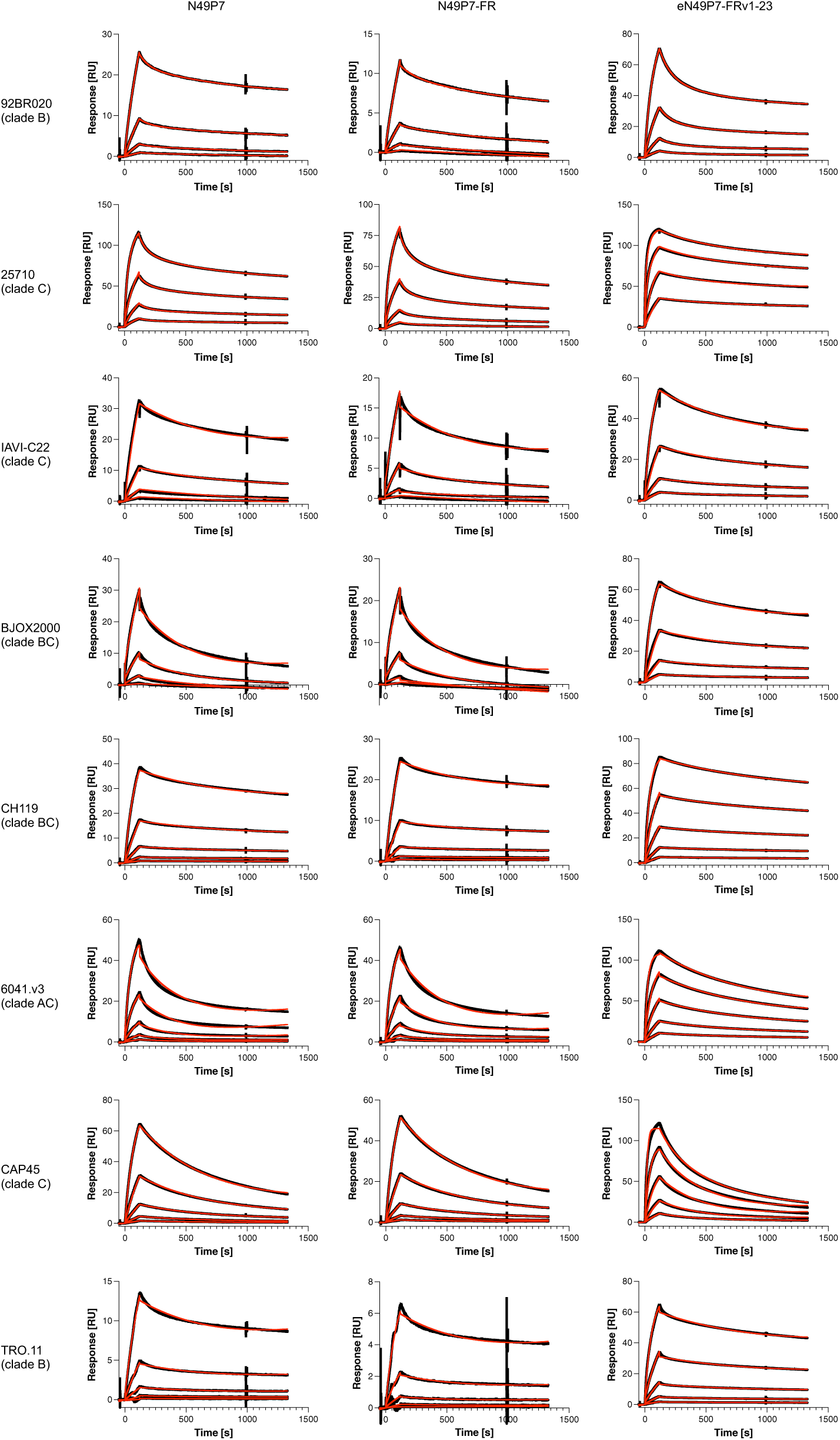
Surface plasmon resonance (SPR) sensorgrams for N49P7 variants binding to a panel of HIV cross-clade gp120s. Sensorgram series were globally fit to either a 1:1 Langmuir binding model or a heterogeneous ligand model (red fit curves). Complete kinetic parameters for all tested gp120s are summarized in Supplementary Table 9.

**Supplementary Figure 22.**
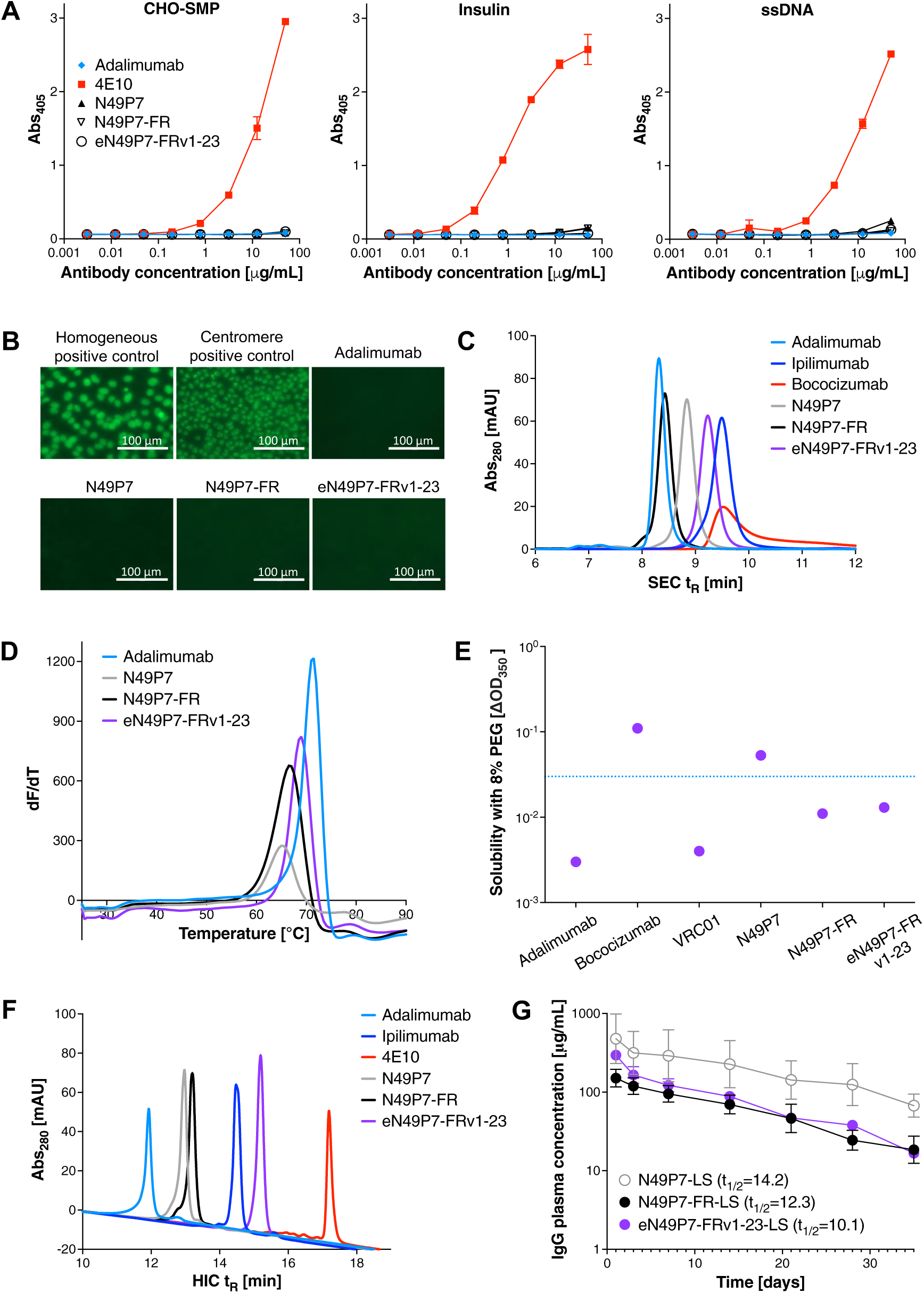
Developability characteristics of the optimized bnAb variant eN49P7-FRv1-23. **A**) Polyreactivity of N49P7 variants with CHO cell membrane proteins (CHO-SMP), insulin, and ssDNA, measured by ELISA in duplicate. Adalimumab (non-polyreactive) and 4E10 (highly polyreactive) were included as antibody controls. **B**) Reactivity of N49P7 variants with HEp-2 cells, assessed by indirect immunofluorescence. Controls included adalimumab (non-reactive) and polyclonal highly reactive antibodies (homogenous and centromere positive controls). The experiment was performed in duplicate. **C**) Analytical size-exclusion chromatography retention time (SEC t_R_) of N49P7 variants on a mAb column (30 cm TSKgel SuperSW mAb HR). Adalimumab and ipilimumab (favorable biophysical properties), and bococizumab (aggregation-prone) were included as antibody controls. The experiment was performed in duplicate. **D**) Thermal stability of N49P7 variants measured by differential scanning fluorimetry (DSF) in triplicate. Melting temperature (T_m_) values were calculated from the peaks of the melt curves, derived from the first derivatives of fluorescence intensity versus temperature. Adalimumab (a stable antibody) served as a control. **E**) Solubility of VRC01 and N49P7 variants in 8% PEG. The horizontal dotted line marks OD_350_=0.03, indicating the onset of visible precipitation. Antibodies with OD_350_>0.03 are predicted to be insoluble at high concentrations (>50 mg/mL). Adalimumab (solution-stable) and bococizumab (aggregation-prone) were used as antibody controls. The experiment was performed in duplicate. **F**) Hydrophobic interaction chromatography retention time (HIC t_R_) of N49P7 variants on a butyl column (10 cm TSKgel Butyl-NPR). Adalimumab and ipilimumab (favorable biophysical properties), and 4E10 (hydrophobic) were included as antibody controls. The experiment was performed in duplicate. **G**) Pharmacokinetics (PK) and plasma half-life (t_1/2_) of N49P7 variants with the LS mutations in hFcRn transgenic mice. Mean ± SD values for 4-7 mice per group are shown.

**Supplementary Figure 23.**
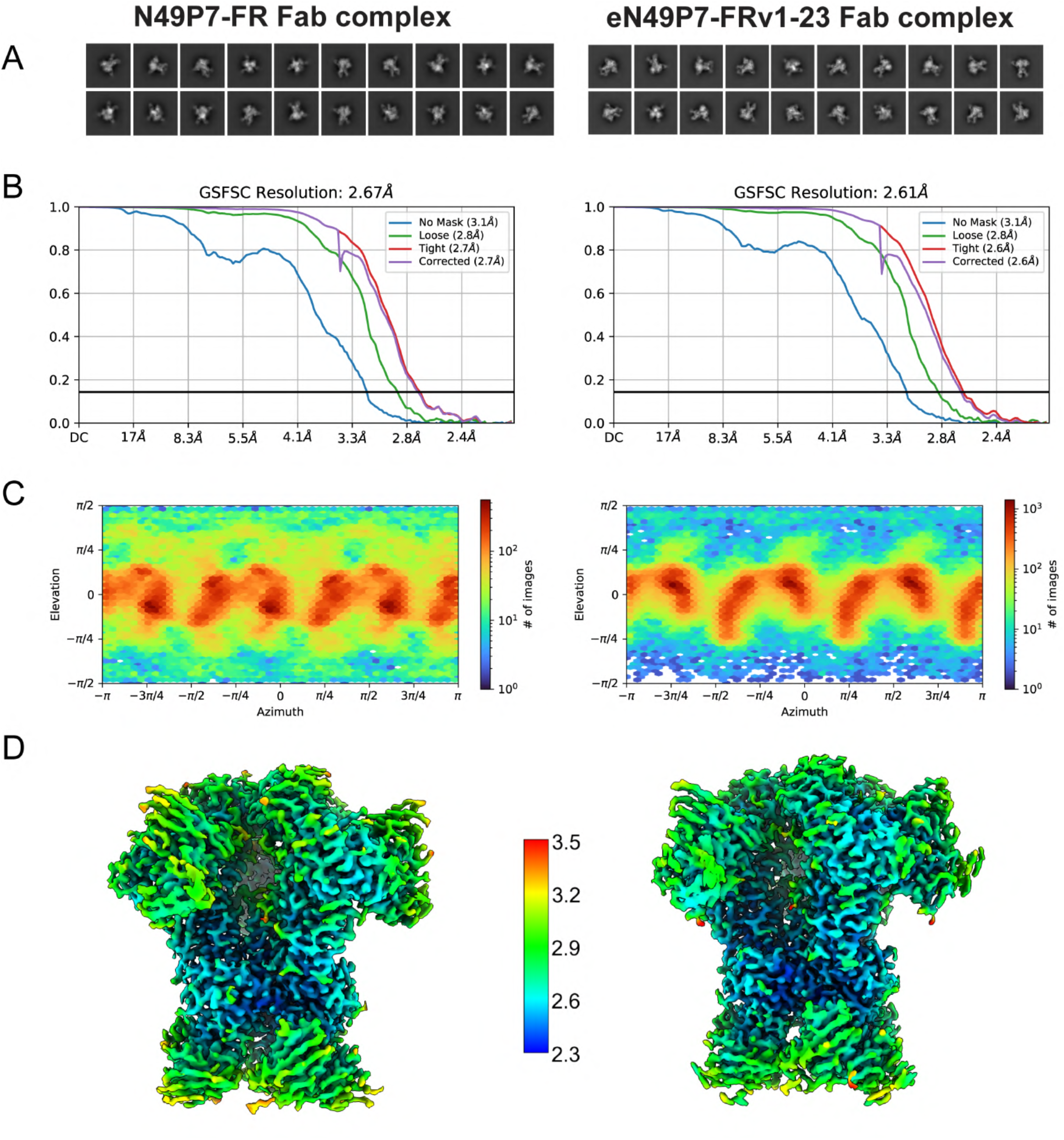
Cryo-EM data processing statistics. **A**) Representative 2D class averages, **B**) Fourier Shell Correlation (FSC) resolution estimates, **C**) angular distribution plots, and **D**) 3D reconstructions colored by local resolution estimates (in units Å) for the N49P7-FR and eN49P7-FRv1-23 cryo-EM complexes.

**Supplementary Figure 24.**
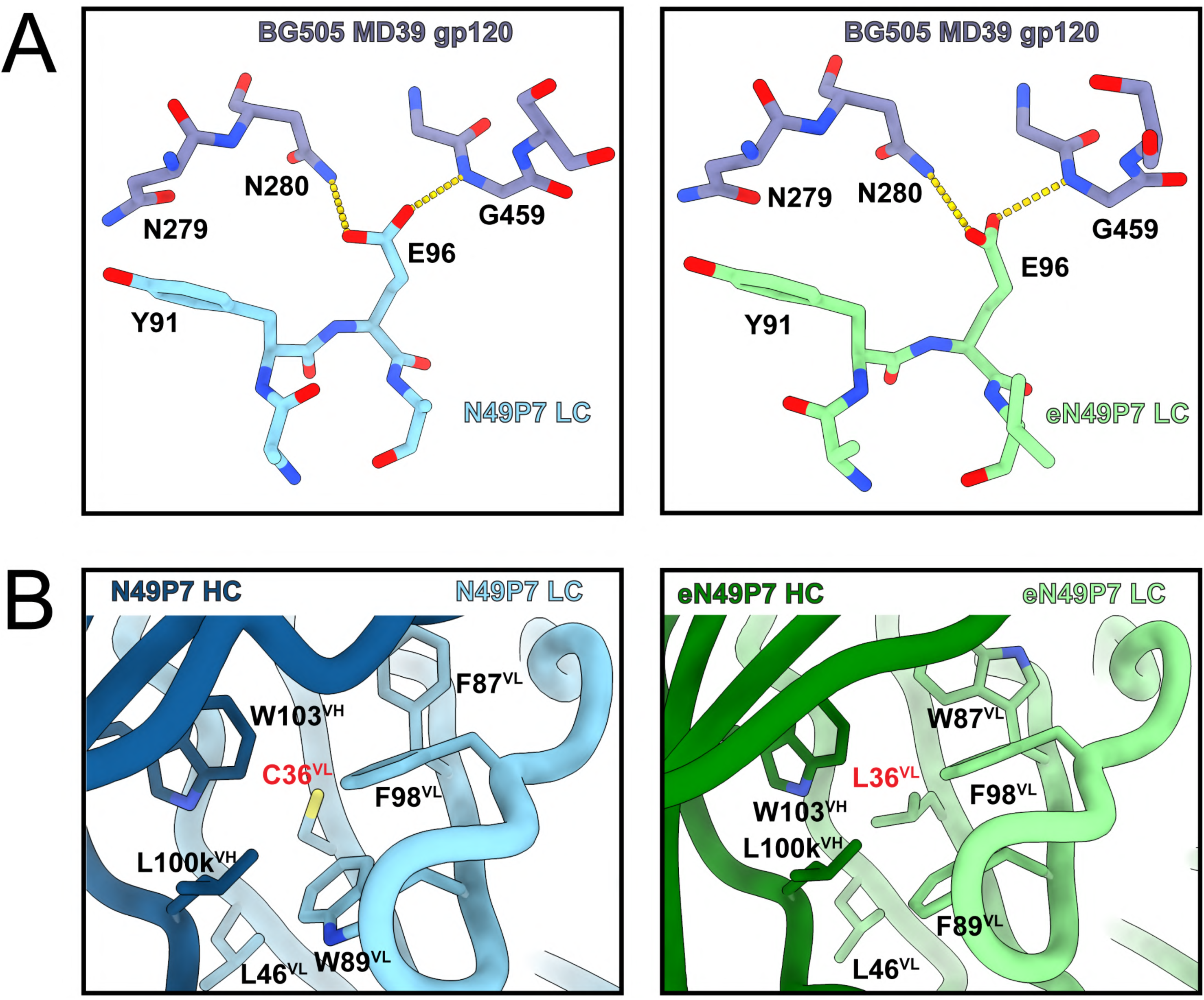
N497P7-FR and eN49P7-FRv1-23 light chain interactions. **A**) CDRL3 “YE motif” interactions with gp120 D-loop (N280) and V5 (G459), typical of VRC01-class antibodies, are conserved in N497P7-FR and eN49P7-FRv1-23. Dashed yellow lines represent putative hydrogen bonds. **B**) Local environment of FRL2 residue 36, which is an unpaired cysteine in N49P7-FR (*left*) and is mutated to leucine in eN49P7-FRv1-23 (*right*), providing better packing in the hydrophobic region composed of the labeled heavy and light chain residues.

**Supplementary Figure 25.**
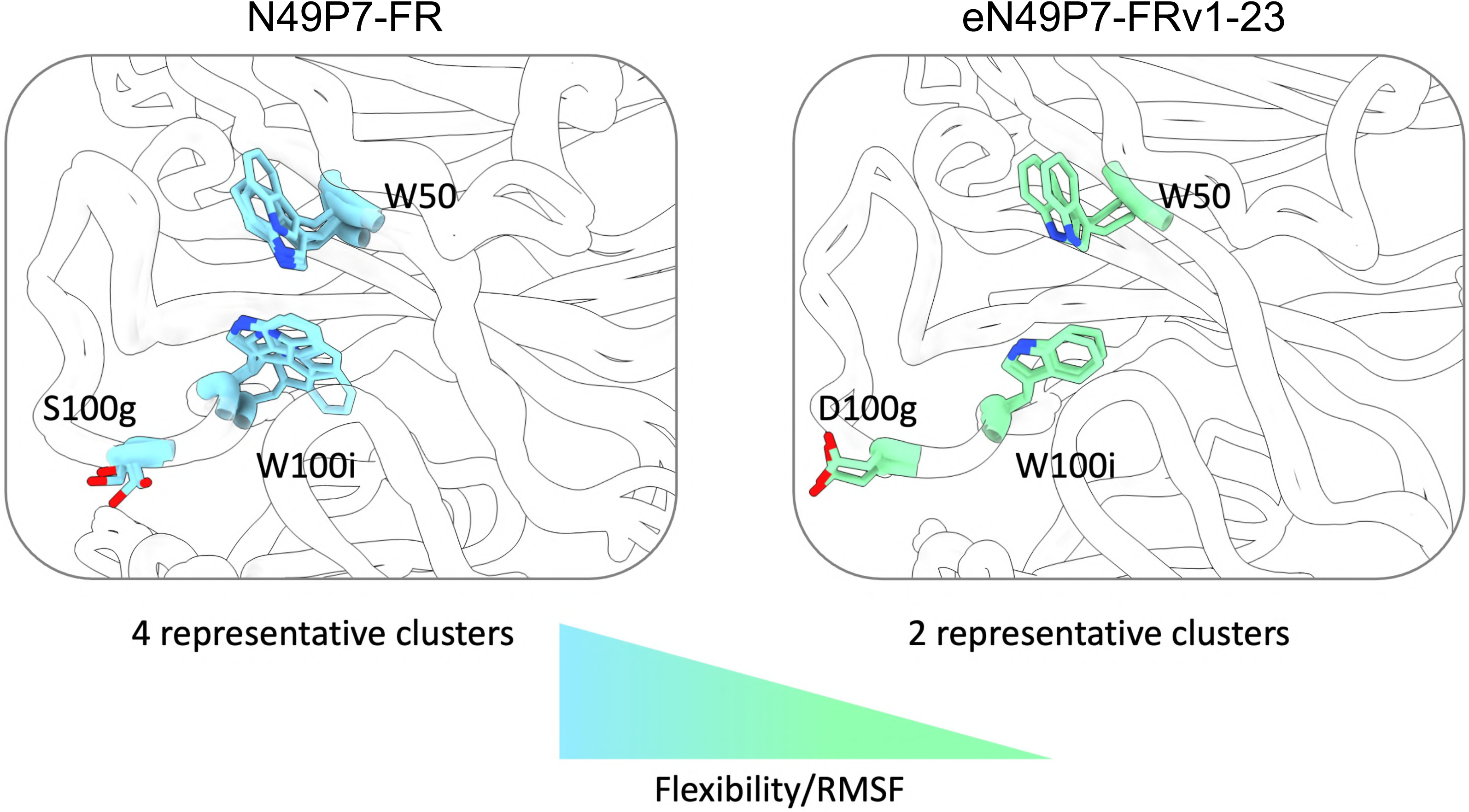
Flexibility characterization and cluster analysis of the antibody binding interface. The rigidification of S100gD^eVH^, W100i^eVH^, and W50^eVH^ in eN49P7-FRv1-23, relative to N49P7-FR, is reflected in decreased Root Mean Square Fluctuation (RMSF), indicating reduced flexibility, and in cluster analysis, which reveals fewer conformations of the binding site. Cluster representatives from MD simulations focusing on these residues show four distinct conformational states in N49P7-FR compared to two in eN49P7-FRv1-23, highlighting the additional stabilization of the binding interface.

## Table descriptions

**Supplementary Table 1.**
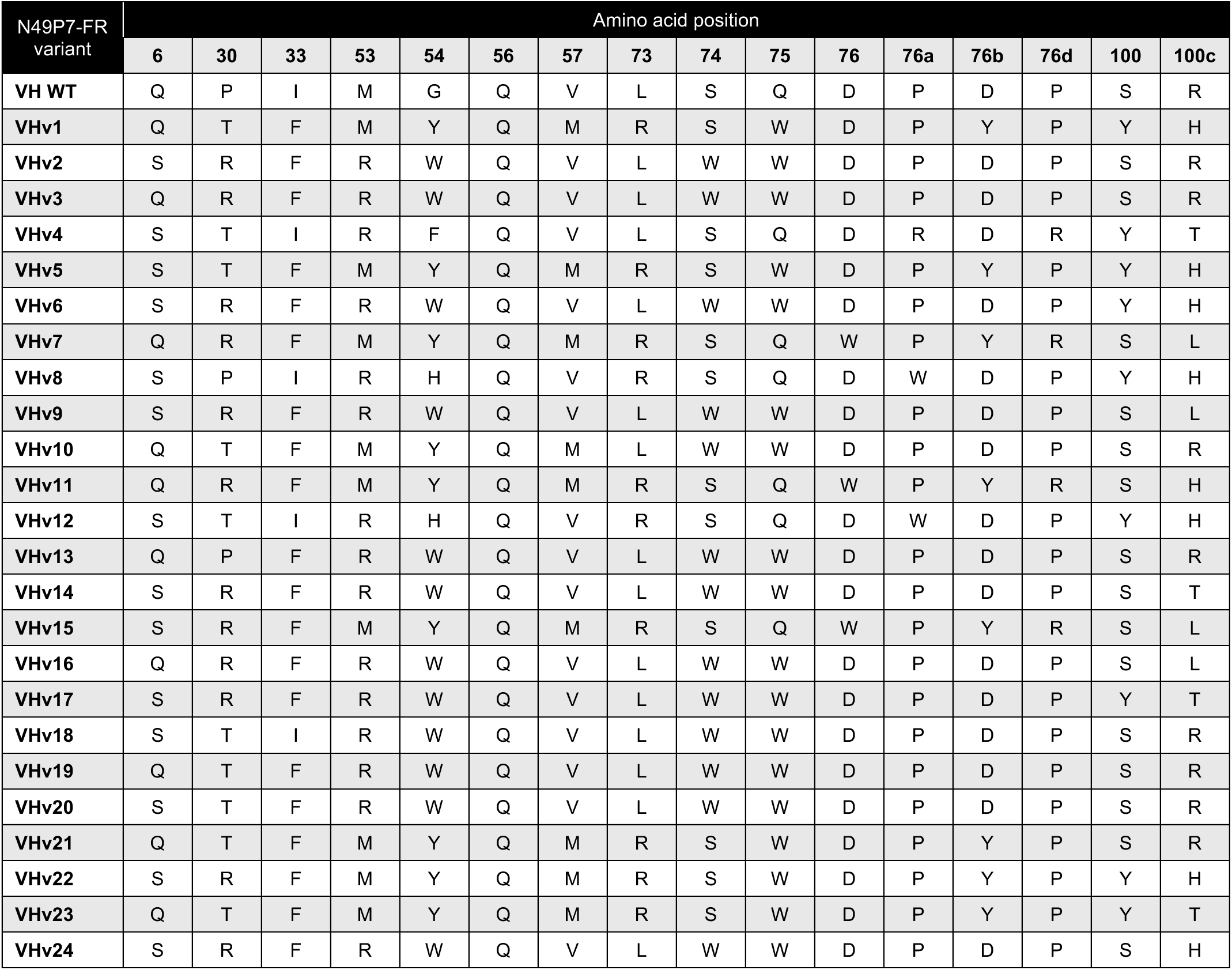
Most enriched combinatorial variants of N49P7-FR VH reformatted as IgG1 and evaluated for functionality and developability. For each variant, only the sampled amino acids are shown, ordered by position and numbered according to the Kabat scheme.

**Supplementary Table 2.**
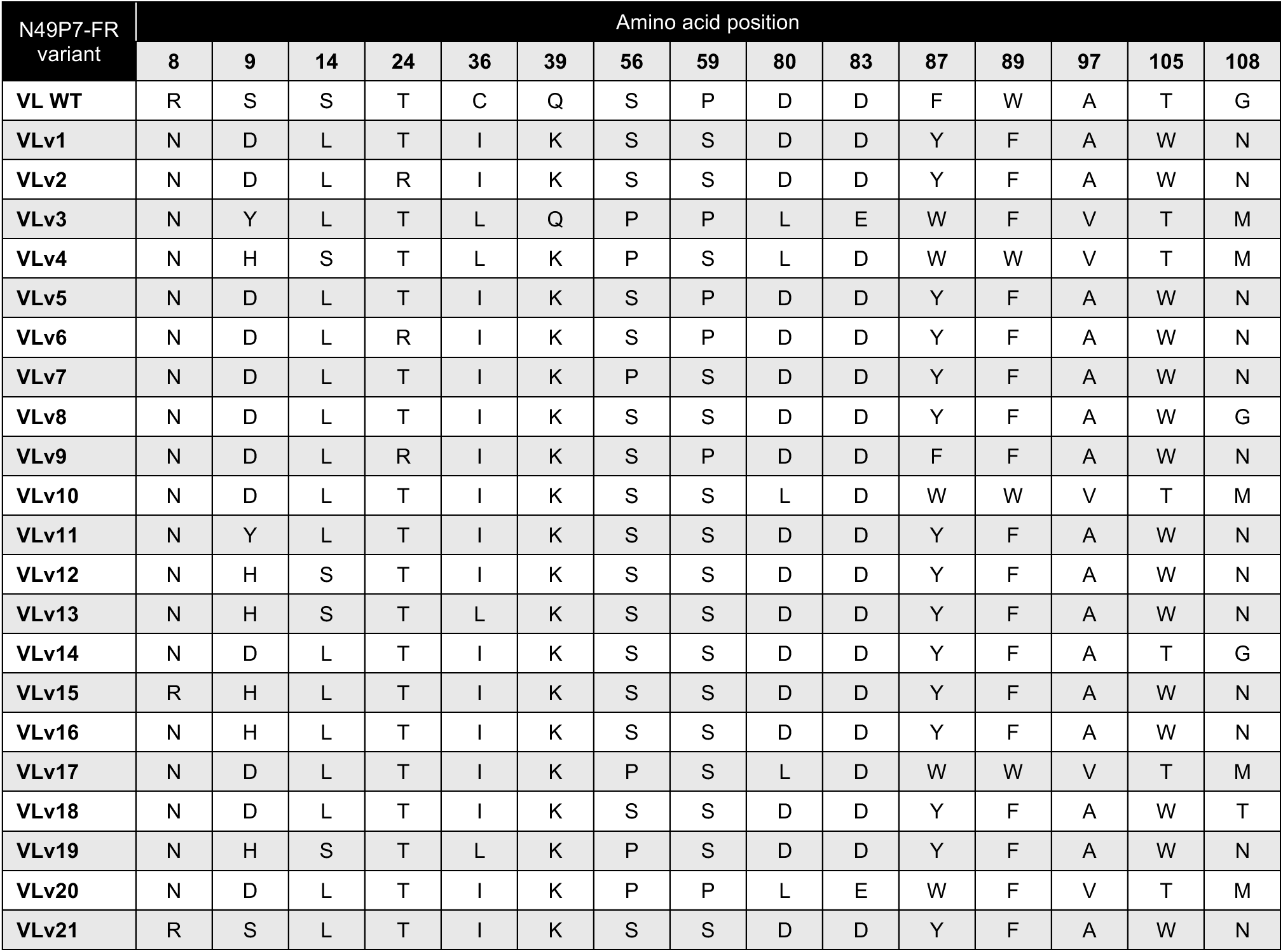
Most enriched combinatorial variants of N49P7-FR VL reformatted as IgG1 and evaluated for functionality and developability. For each variant, only the sampled amino acids are shown, ordered by position and numbered according to the Kabat scheme.

**Supplementary Table 3.**
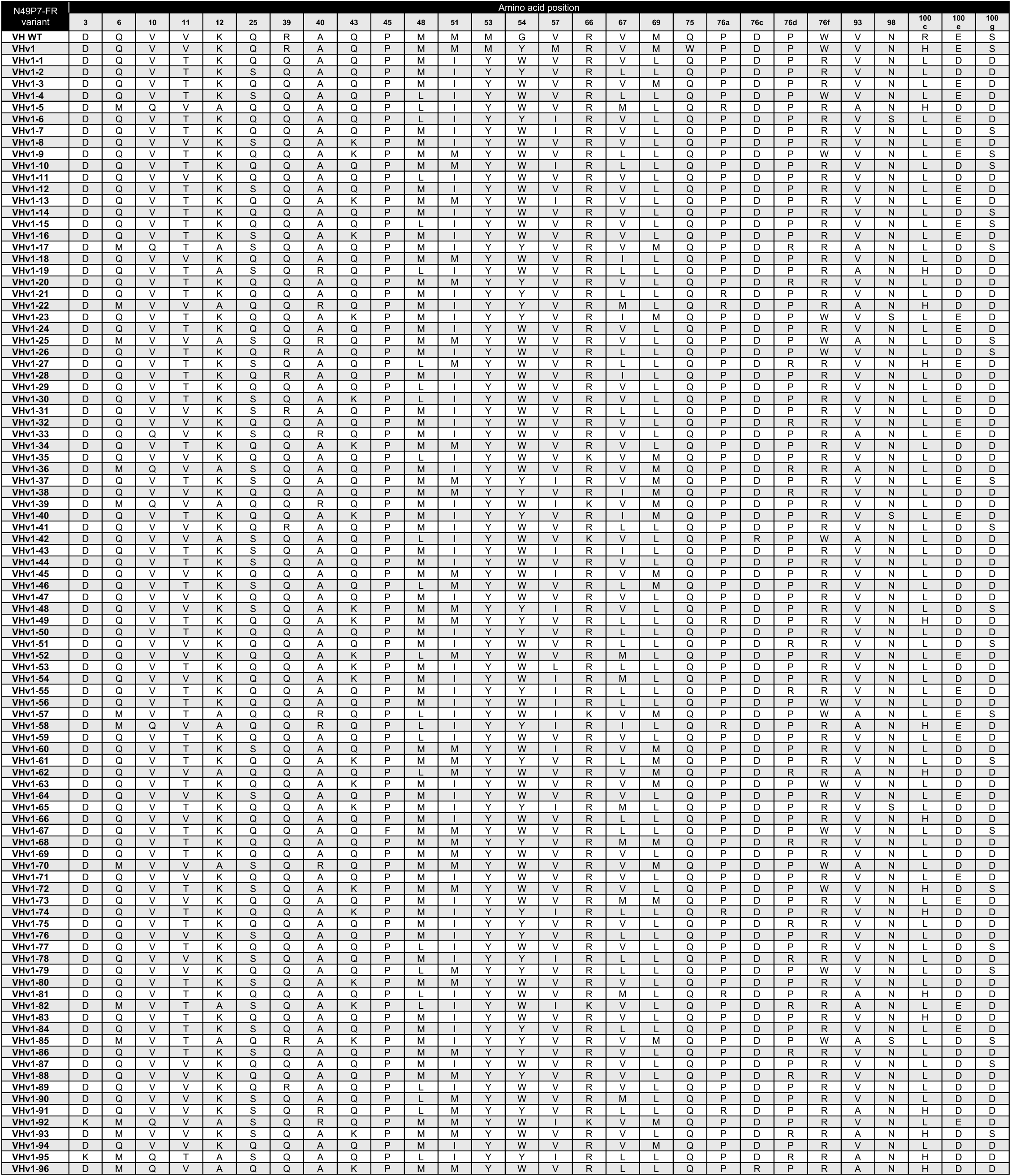
Most enriched combinatorial variants of N49P7-FR VHv1 reformatted as IgG1 and evaluated for functionality and developability. For each variant, only the sampled amino acids are shown, ordered by position and numbered according to the Kabat scheme.

**Supplementary Table 4.**
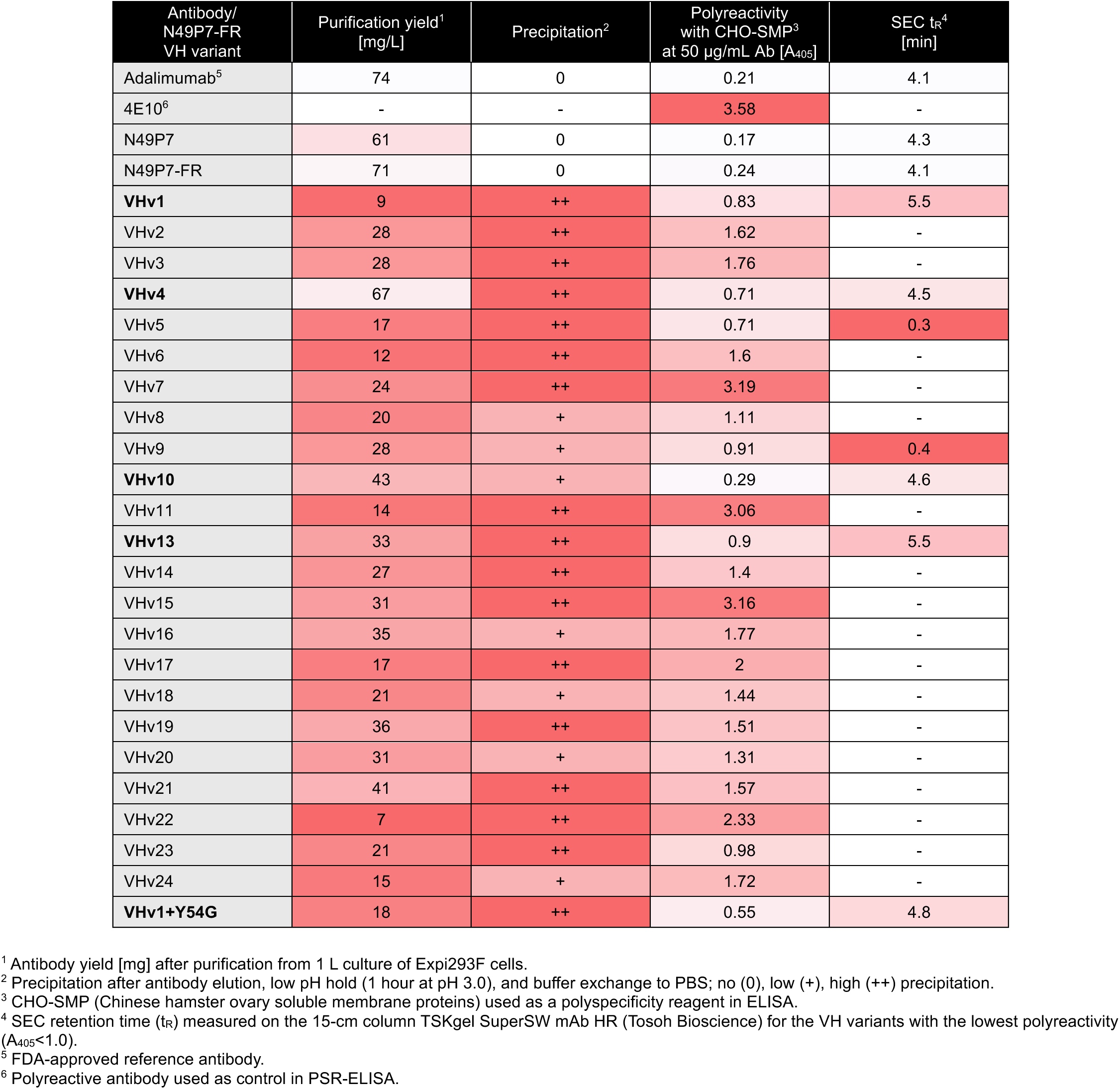
Developability characteristics of the most enriched combinatorial variants of N49P7-FR VH reformatted as IgG1. Red shading indicates developability liabilities, with color intensity proportional to severity. Variants in bold demonstrated the most favorable biophysical profiles and were selected for HIV neutralization analysis (Supplementary Fig. 14).

**Supplementary Table 5.**
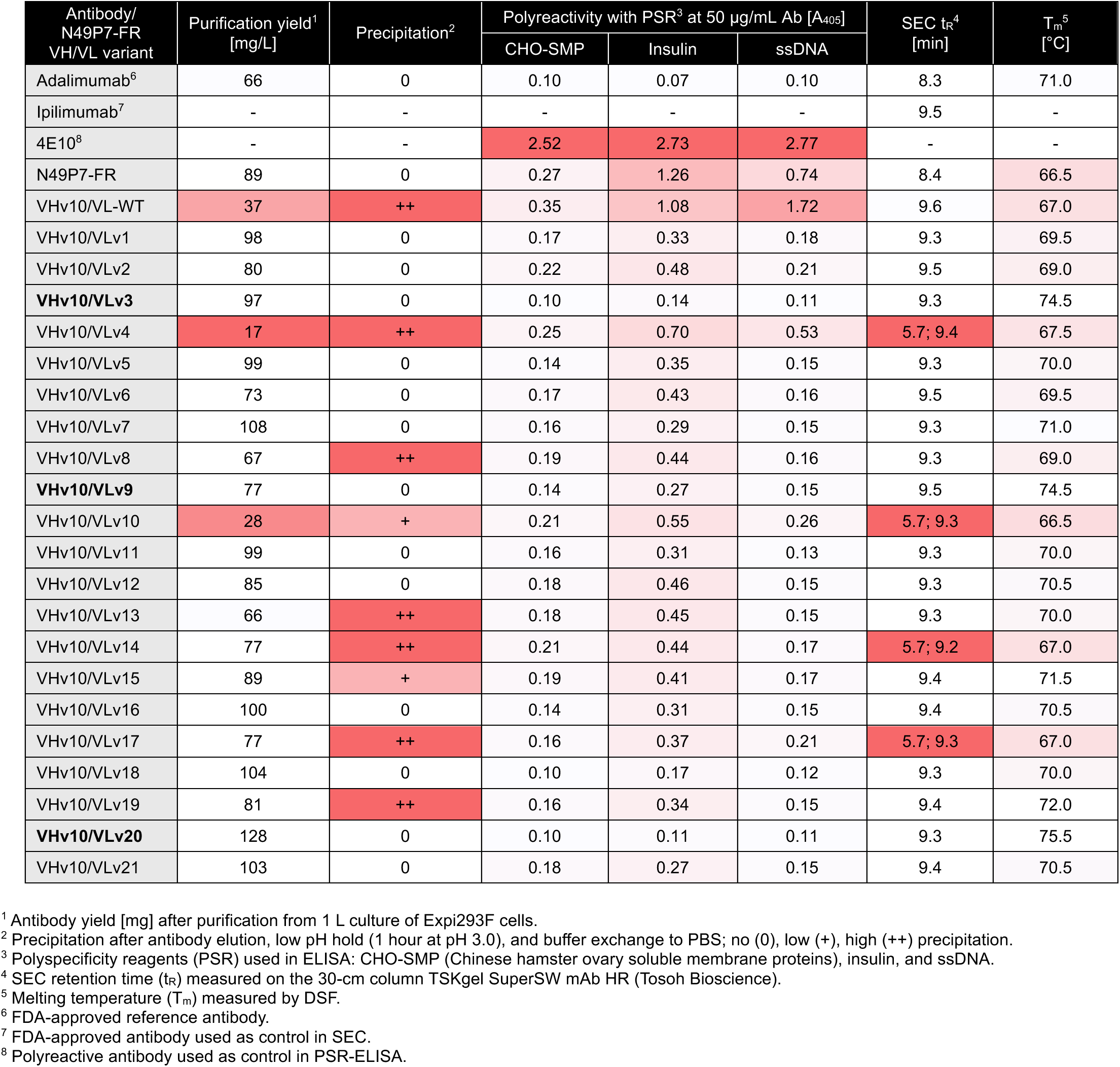
Developability characteristics of the most enriched combinatorial variants of N49P7-FR VL reformatted as IgG1. Red shading indicates developability liabilities, with color intensity proportional to severity. Variants in bold demonstrated the most favorable biophysical profiles and were selected for HIV neutralization analysis (Supplementary Fig. 15).

**Supplementary Table 6.**
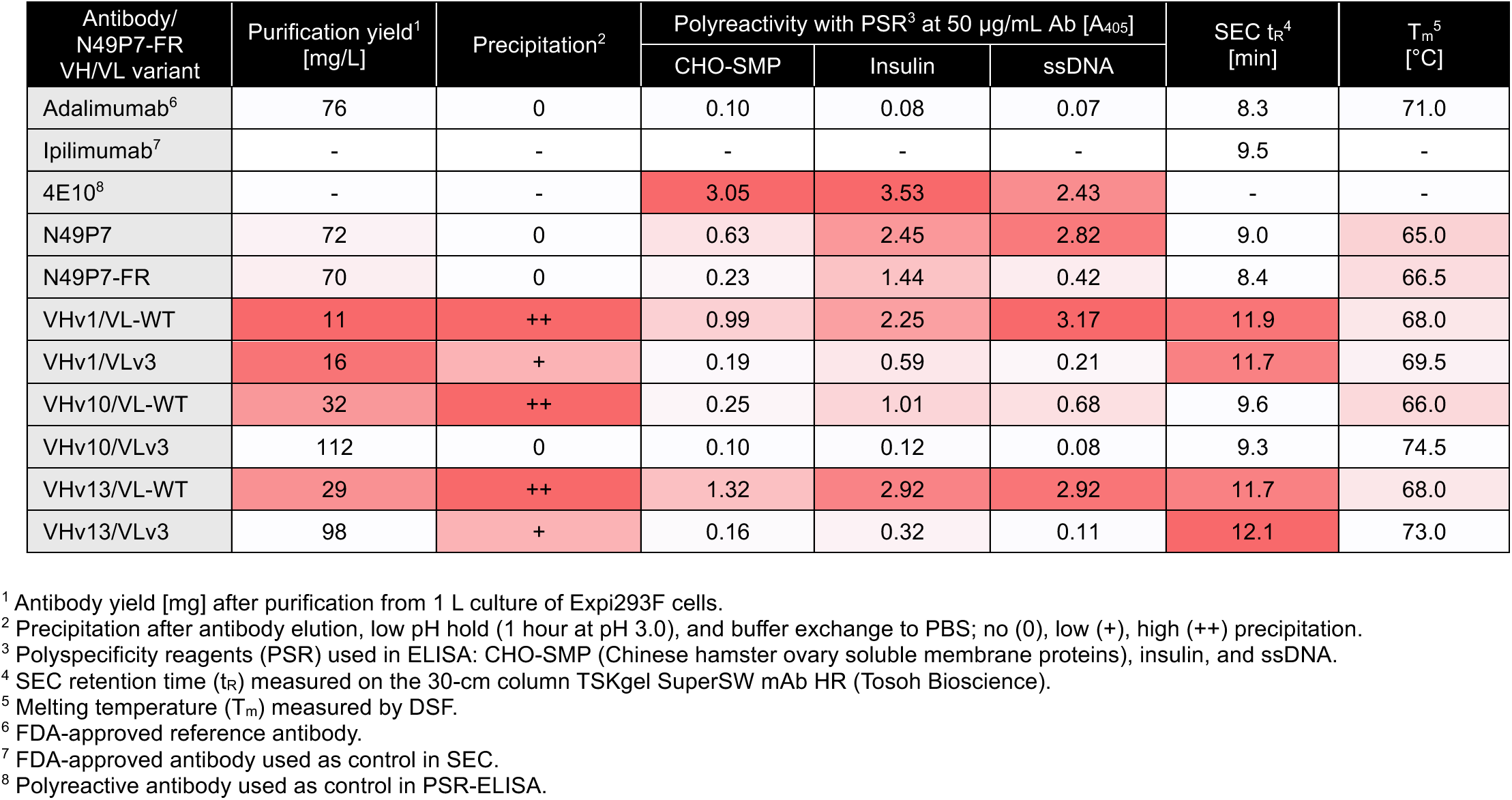
Developability characteristics of top N49P7-FR VH variants shuffled with VLv3. Red shading indicates developability liabilities, with color intensity proportional to severity. Variants were subsequently evaluated for HIV neutralization (Supplementary Fig. 16).

**Supplementary Table 7.**
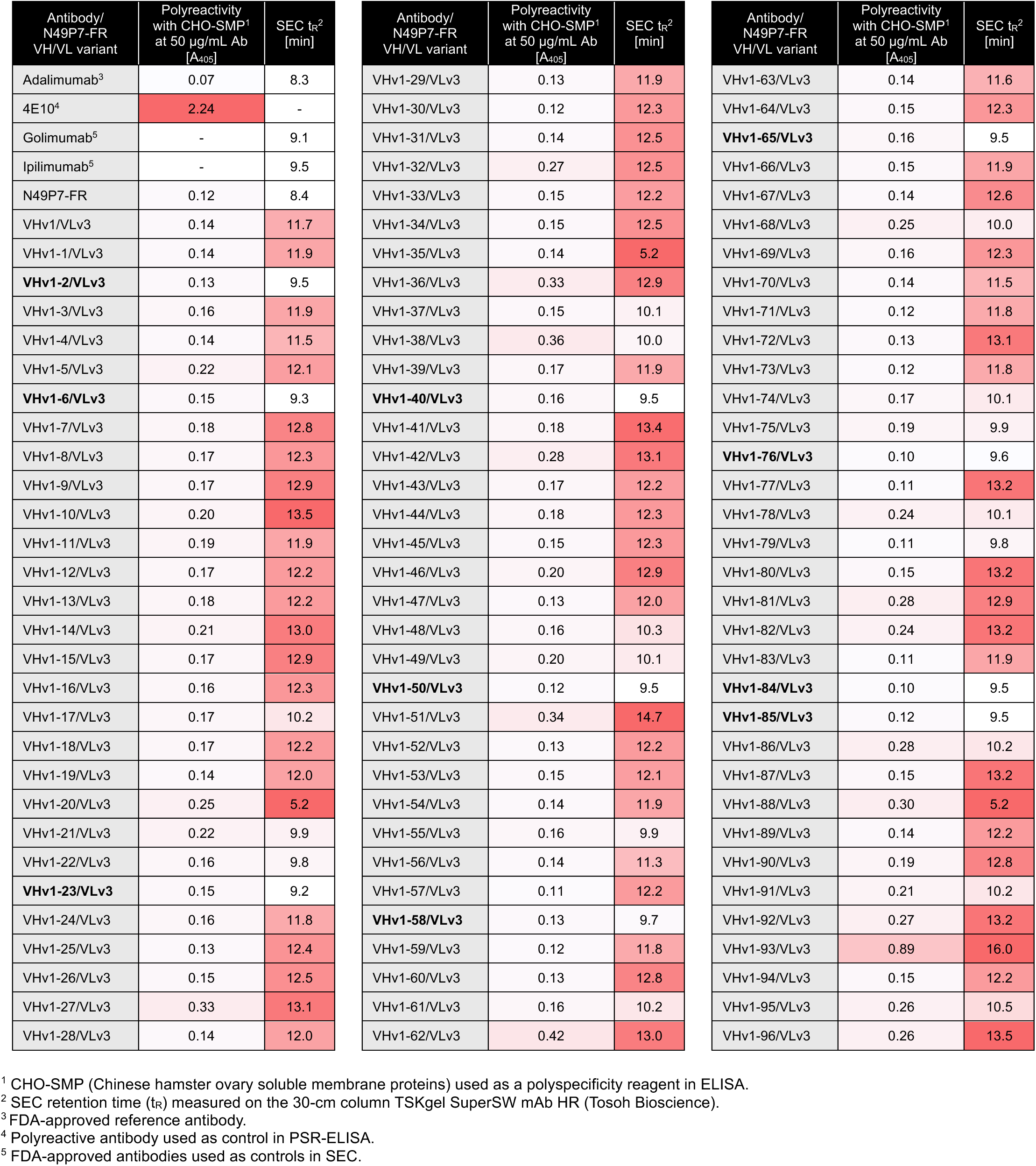
Developability characteristics of the most enriched combinatorial variants of N49P7-FR VHv1 paired with VLv3 and reformatted as IgG1. Red shading indicates developability liabilities, with color intensity proportional to severity. Variants in bold had the most favorable SEC profiles and were selected for HIV neutralization analysis (Supplementary Fig. 17).

**Supplementary Table 8.**
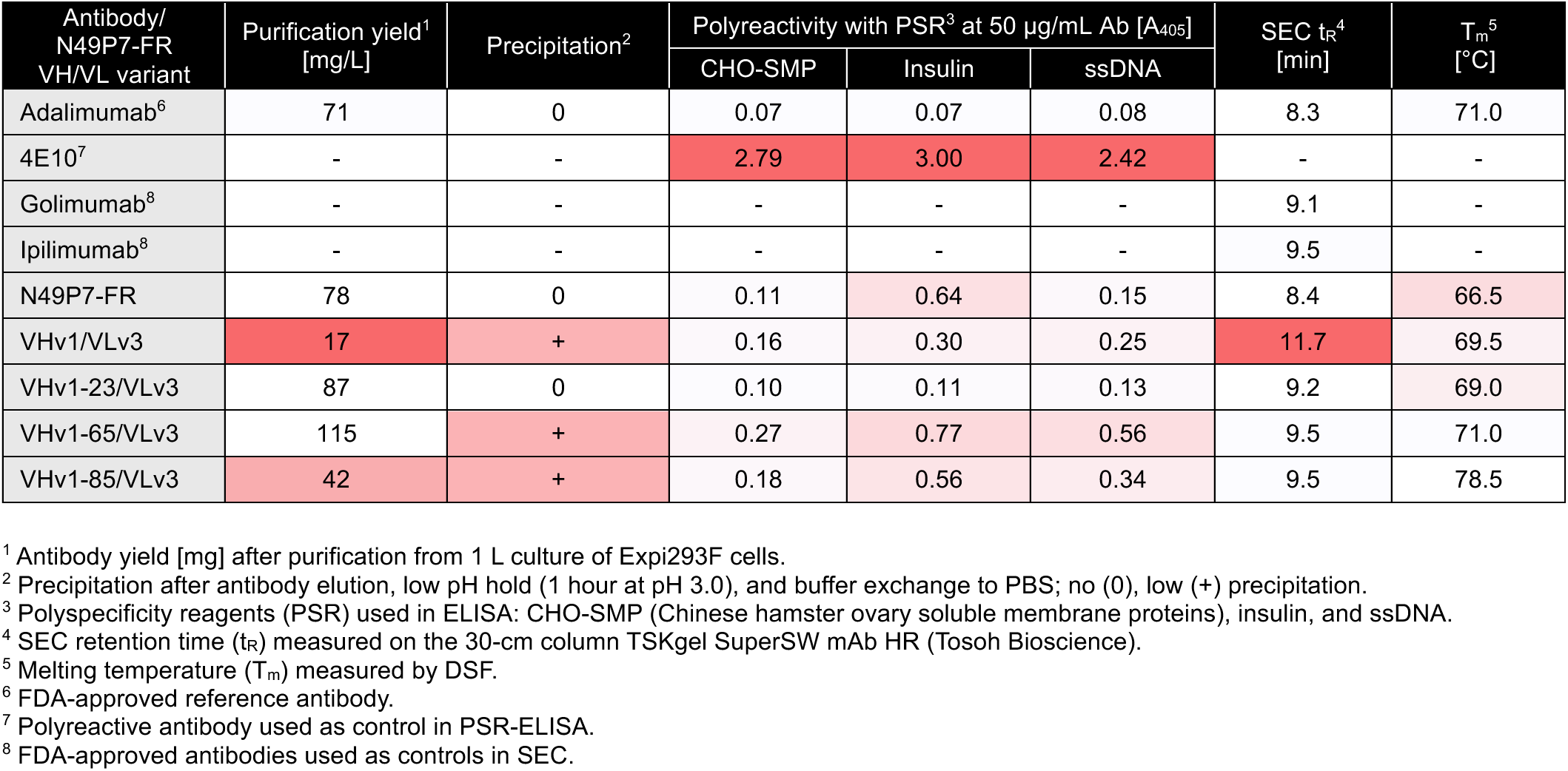
Developability characteristics of the most potent N49P7-FR VHv1 variants. Red shading indicates developability liabilities, with color intensity proportional to severity. Variants were subsequently evaluated for HIV neutralization (Supplementary Figs. 18–19).

**Supplementary Table 9.**
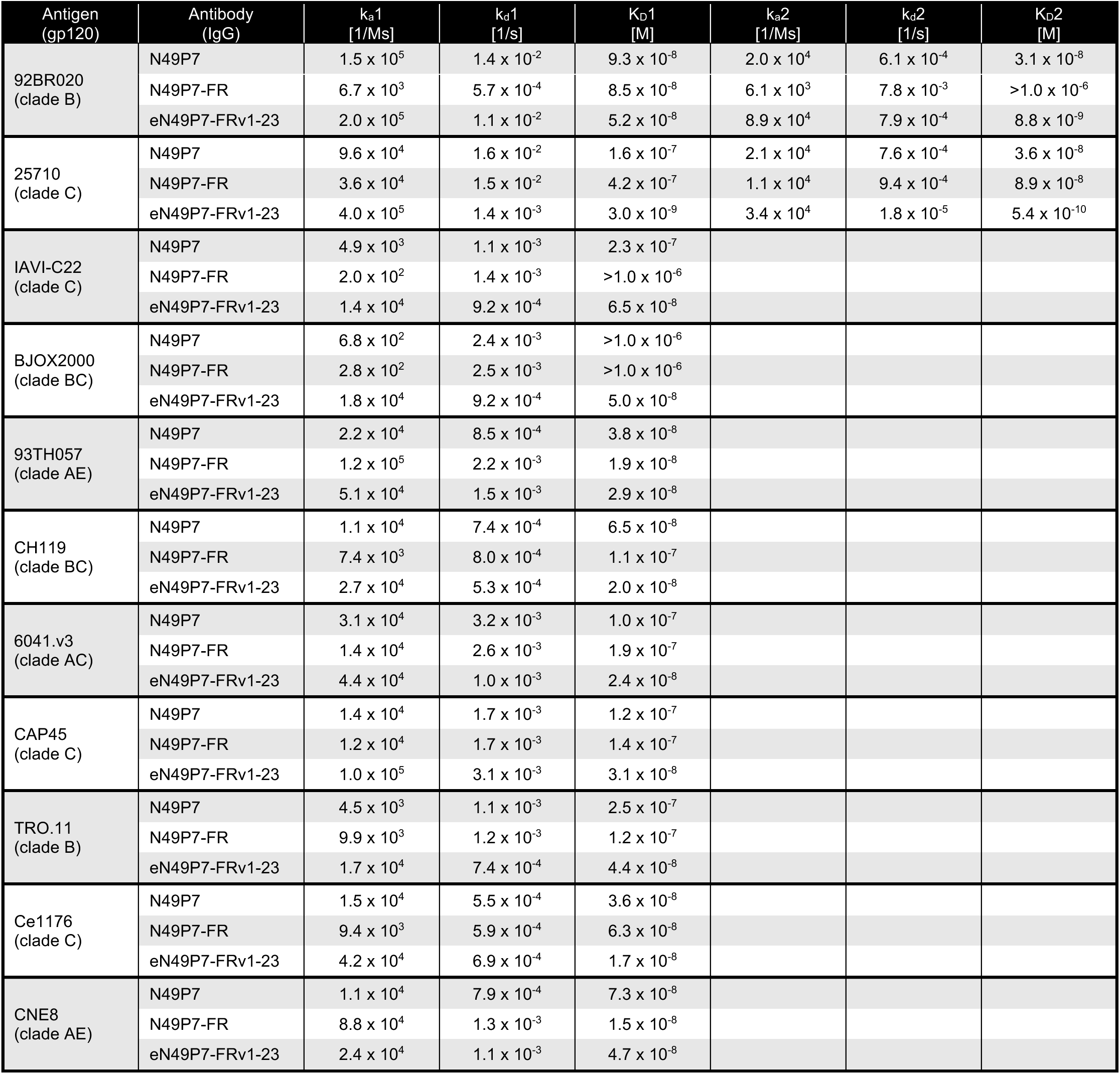
Kinetic parameters for N49P7 variants binding to a panel of HIV cross-clade gp120s. The association rate constant (k_a_), dissociation rate constant (k_d_), and equilibrium dissociation constant (K_D_) were derived from SPR sensorgrams globally fit to either a 1:1 Langmuir binding model (one k_a_, k_d_, and K_D_ value) or a heterogeneous ligand model (two sets of k_a_, k_d_, and K_D_ values). eN49P7-FRv1-23 exhibited improved binding kinetics relative to the parental N49P7-FR for 9 of the 11 tested gp120s (92BR020, 25710, IAVI-C22, BJOX2000, CH119, 6041.v3, CAP45, TRO.11, and Ce1176).

**Supplementary Table 10.**
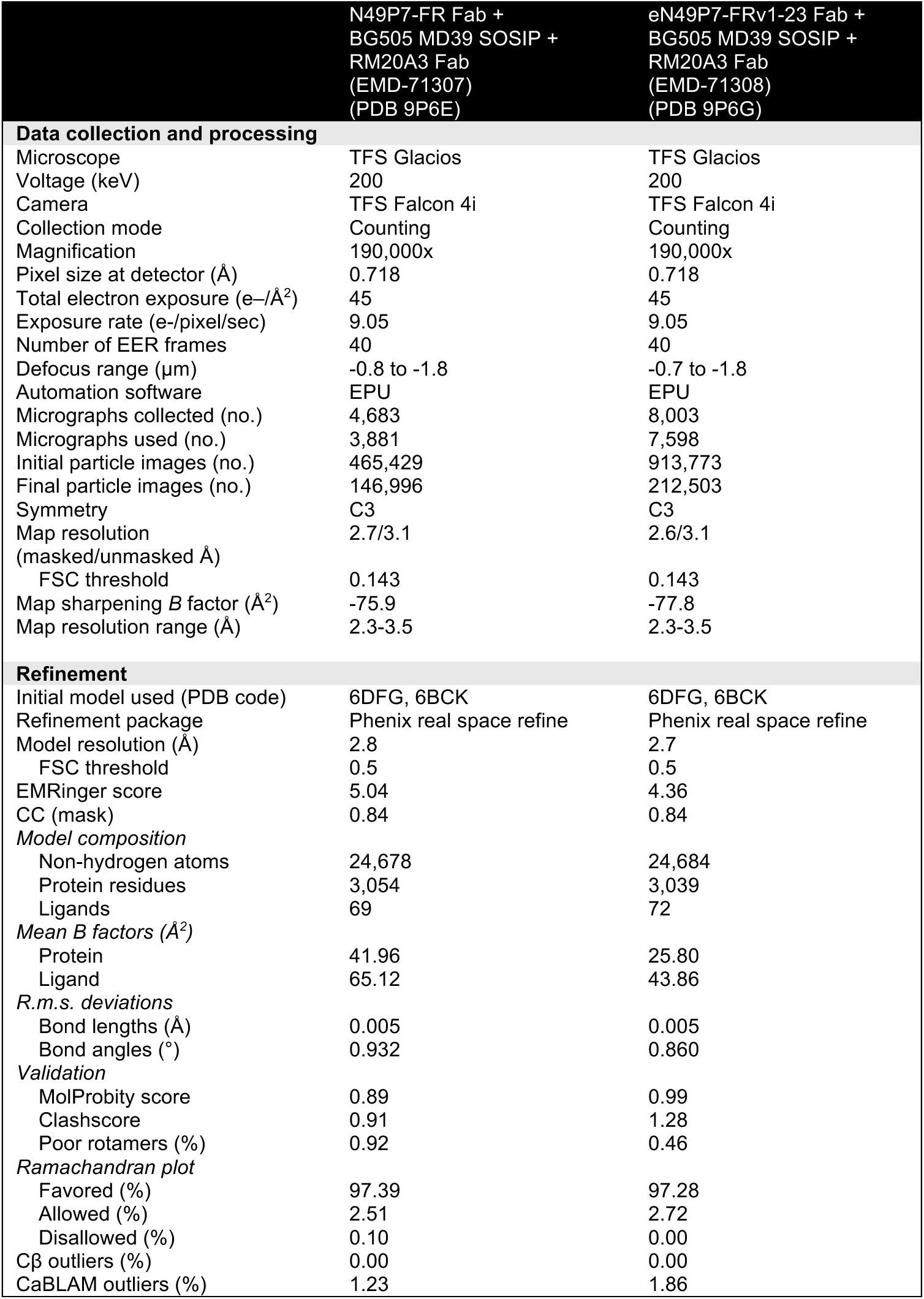
Cryo-EM data collection, refinement, and validation statistics.

**Supplementary Table 11.**
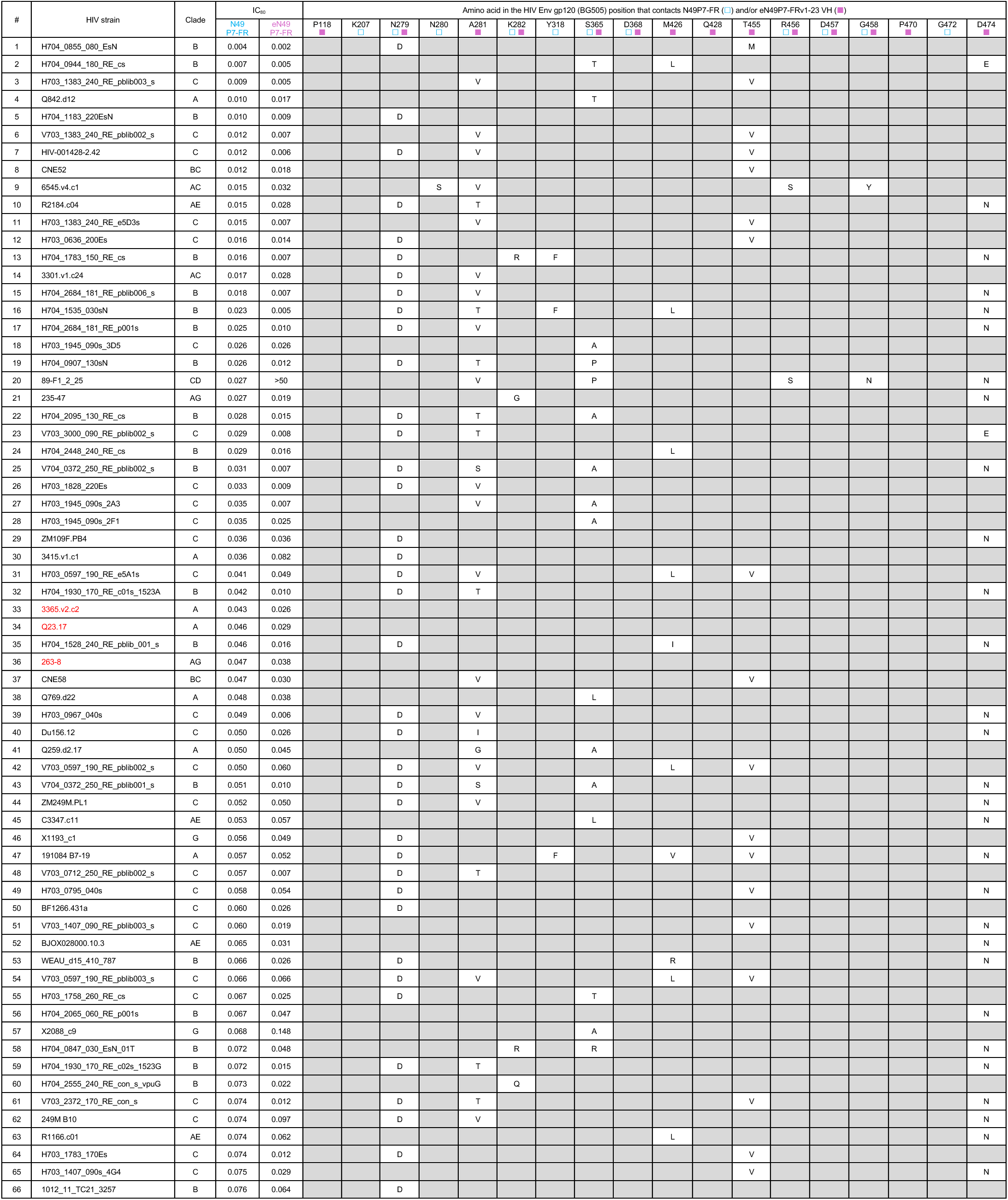
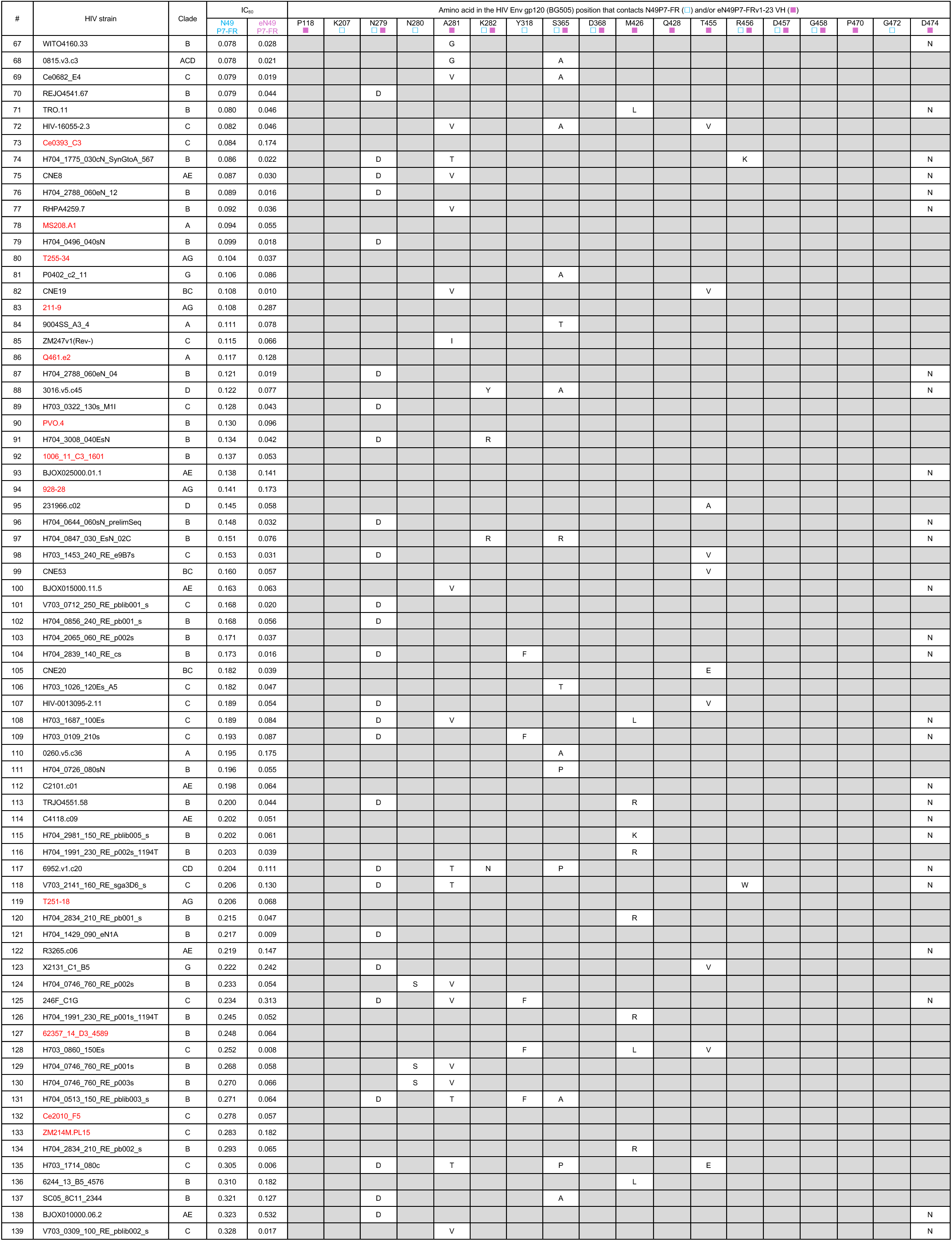
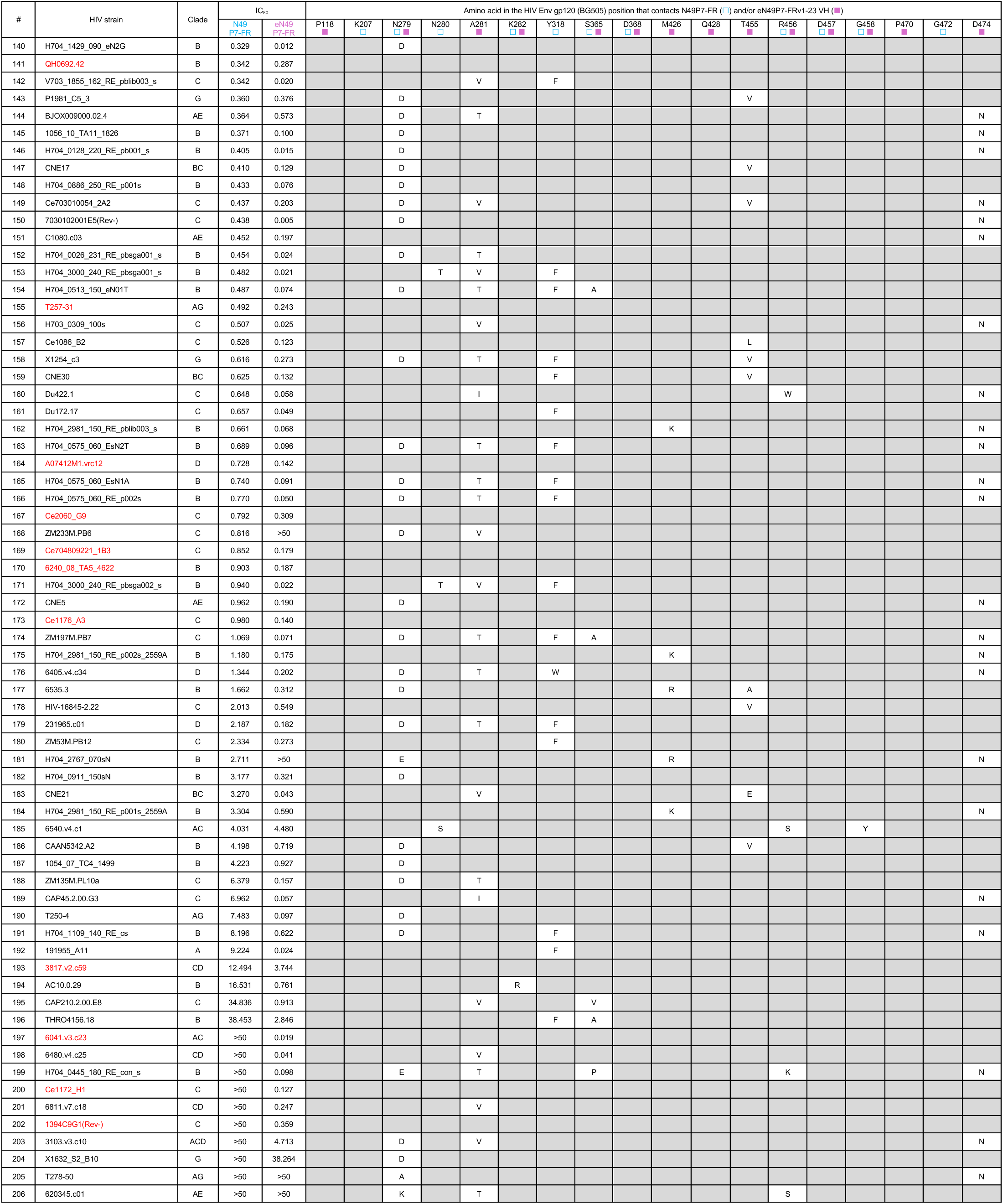
Amino acid variation at N49P7-FR and eN49P7-FRv1-23 contact sites in the HIV Env of 206 strains. Contact sites were determined by cryo-EM analysis of N49P7-FR or eN49P7-FRv1-23 in complex with BG505 MD39 SOSIP. For each strain, amino acid substitutions relative to BG505 MD39 SOSIP at these positions are shown; gray shading indicates no substitution. All Env sequences were derived from pseudoviruses used in the *in vitro* neutralization assay, for which the potency of both bnAbs is reported as IC_80_ values. Strains highlighted in red have no mutations at any of the identified bnAb contact positions.

